# Human Lymph Node Cellular Senescence Atlas Reveals Age-Dependent Alteration in Germinal Center B Cell Function and Niches

**DOI:** 10.64898/2026.04.02.716161

**Authors:** Negin Farzad, Archibald Enninful, Yao Lu, Fabio Parisi, Anthony Fung, Yumi Kwon, Yajuan Li, Margaux Labrosse, Mingyu Yang, Francesco Strino, Liang Chen, Junchen Yang, Mei Zhong, Fu Gao, Bo Tao, Joseph Cunningham, Zhiliang Bai, Haikuo Li, Fang Wang, Michael Stankewich, Dongjoo Kim, Mingze Dong, Lisa M. Bramer, Keyi Li, Meera R. Bhat, Evan Loe, Joseph Craft, Ljiljana Pasa-Tolic, Stephanie Halene, Lingyan Shi, Yuval Kluger, Mina L. Xu, Rong Fan

## Abstract

Immunosenescence, the age-associated decline in immune function, is a key feature of human aging. In human lymphoid organs, however, the specific immune cell populations that acquire senescence-associated phenotypes during aging and how they influence the surrounding tissue microenvironment remain poorly understood. A spatially resolved map of these senescence-associated immune states in human lymphoid tissues could help clarify their relationship with aging and their potential contributions to the progressive decline of immune function. Here, we integrated single-cell and spatial multi-omics to systematically characterize age-related senescence in human lymph nodes (LNs). Single-cell transcriptomics of lymphoid tissues from donors aged 18 to 100 years old identified 34 immune and stromal cell types and revealed age-associated upregulation of senescence signatures in specific populations. Spatial proteomic profiling of 99 LN sections from 51 donors (18–86 years) using high-plex immunofluorescence (∼20 million cells) mapped senescence markers (p16, p21, HMGB1, *𝛾*-H2AX) at single-cell resolution, revealing diverse senescent-like cell types (“senotypes”) and a stepwise shift from extrafollicular to germinal center (GC) localization with age. Notably, we observed focal clonal-like senescence in GC B cells in older donor LNs. Spatial transcriptomics, epigenomics, and metabolic imaging of selected samples further elucidate the multi-omics signatures and underlying mechanisms of functional impairment, metabolic remodeling, and distinct regulatory programs in senescent-like GC B cells. This study presents a comprehensive spatial atlas of senescence-associated immune states in human lymph nodes, revealing cell-type–specific and spatial heterogeneity that may contribute to immunosenescence and the decline of immune function during aging.

## INTRODUCTION

Immunosenescence, the age-associated decline in immune function, is a major contributor to increased susceptibility to infections, reduced vaccine efficacy, and the development of chronic inflammatory diseases in older individuals^1–5^. While cellular senescence^6^ has traditionally been defined as a stable state of cell-cycle arrest triggered by stressors such as DNA damage, oxidative stress, or telomere shortening, immune cells during aging often adopt senescence-associated phenotypes characterized by altered differentiation states and the production of pro-inflammatory factors resembling the senescence-associated secretory phenotype (SASP)^7,8^. This inflammatory secretory program can disrupt tissue homeostasis and contribute to the chronic low-grade inflammation associated with aging^9,10^.

Within the immune system, the accumulation of such senescence-like immune states has been implicated in impaired immune surveillance and dysregulated immune responses^1,2,4,9^. Because immune cells play central roles in maintaining tissue homeostasis across organs, these changes may broadly influence physiological function during aging^10^. Despite increasing recognition of immunosenescence as a key hallmark of aging, the cellular identities, functional states, and tissue contexts of senescence-associated immune cells in human tissues remain poorly characterized. Defining the heterogeneity and spatial organization of these cells in vivo in intact human tissues is therefore critical for understanding how immunosenescence emerges and contributes to age-related functional decline.

We present a spatial atlas of cellular senescence in human lymph nodes (LNs) from 51 donors across Group 1 (18-50 years old), Group 2 (50–70 years old), Group 3 (70–100 years old) generated by multi-modal spatial omics including CODEX^11^, DBiT-seq^12^, spatial epigenome sequencing^13^, CosMx^14^, mass spectrometry proteomics, and SRS metabolic imaging^15^. We mapped 34 immune and stromal cell types and revealed conserved senescence signatures that accumulate with age. Senescent-like cells marked by p16, p21, HMGB1, and *𝛾*-H2AX not only increased in abundance but shifted spatially from extrafollicular zones to the mantle zones and germinal centers with age. Senescence hotspots were observed in B cells including memory, naïve, and germinal-center subsets, showing focal, clonal-like accumulation. Spatial transcriptomic and epigenomic profiling uncovered diverse regulatory programs involving NF-kB, AP-1, and KLF transcription factors, with increased chromatin accessibility at *CDKN1A, CDKN2A*, and SASP genes. Metabolic imaging revealed altered lipid-to-protein ratios in senescent B cells, suggesting functional adaptation. Together, this atlas maps the cellular and molecular features of immune senescence in human lymph nodes and points to potential targets for modulating age-related immune decline.

## RESULTS

### Approach to a multimodal atlas of human lymph node

To investigate senescence-associated and age-related changes in human lymph nodes (LNs), we analyzed whole LN resections or needle biopsy tissue blocks from 65 donors (41 males, 24 females; 108 total samples) obtained through the Yale Pathology Tissue Service. LN tissue samples were stratified into three age groups: Group 1 (<50 years, n = 26), Group 2 (50–69 years, n = 15), and Group 3 (≥70 years, n = 10), with median ages of 32, 56, and 75 years, respectively (Figures 1, S1; Table S1). All tissues were histologically normal as confirmed by board-certified pathologists. To enable multimodal spatial profiling, both fresh-frozen (FF) and formalin-fixed paraffin-embedded (FFPE) tissues were included. FF samples were processed for DBiT-based spatial transcriptomics^12^ and spatial ATAC sequencing^13^ (chromatin accessibility), as well as spatially resolved mass spectrometry-based proteomics; FFPE samples were assayed using CODEX (multiplexed immunofluorescence)^11^, CosMx spatial molecular imaging^14^, FFPE-compatible Patho-DBiT sequencing^16^, and metabolic imaging via stimulated Raman scattering (SRS) microscopy^15^. Tissues were serially sectioned onto assay-specific slides to preserve morphology and ensure assay compatibility, including standard slides for H&E/IHC, poly-L-lysine (PLL) slides for DBiT-seq and CODEX, and Superfrost Plus slides for CosMx (Figure S1B). From these sections, we constructed a comprehensive spatial multi-omics atlas by integrating transcriptomic, epigenomic, and proteomic data, resulting in a multi-modal cellular senescence atlas. In parallel, we built a single-cell transcriptomics reference atlas by integrating publicly available scRNA-seq datasets from Tabula Sapiens and the Secondary Lymphoid Organ (SLO) Atlas, encompassing approximately 510,000 single-cell transcriptomes from 14 SLO samples and 25 peripheral blood donors ^17,18^ (Figure 2A). Senescent-like cells (SnCs) in tissues were identified in situ using CODEX with four canonical markers: p16, p21, HMGB1, and *𝛾*-H2AX (Figure 3A,B). The final integrated atlas comprises spatial data from over 20 million single cells from 56 individuals, including 18 whole lymph node samples and 81 tissue cores profiled with CODEX, 11 samples analyzed with DBiT spatial RNA transcriptomics, and 9 with spatial ATAC-seq. From these datasets, 1–2 samples of high interest were selected for in-depth characterization using CosMx, mass spectrometry-based proteomics, and stimulated Raman scattering (SRS), respectively (Figure 4, Figures S6–S9; Table S2). This resource enables high-resolution analysis of SnC abundance, spatial distribution, and microenvironmental context in human lymph nodes during aging, including investigation of senescence-associated secretory phenotype (SASP), DNA damage response (DDR), inflammaging, and immunosenescence pathways, as well as their connections to metabolic and epigenetic mechanisms.

**Figure 1.**
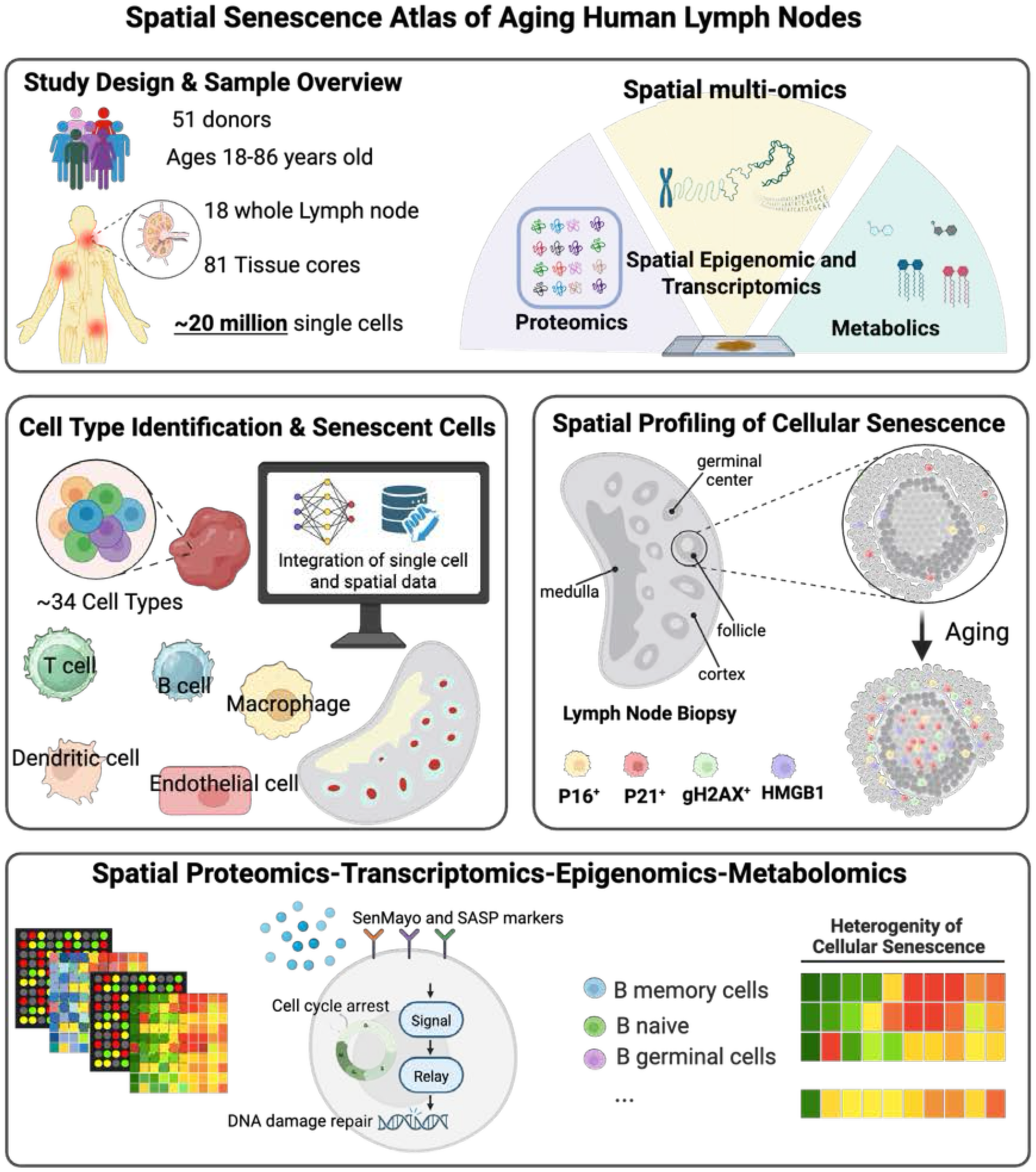
Study design: the human lymph node cellular senescence atlas across ages. Schematic of the study workflow, showing the collection of human lymph node samples from donors of various ages, followed by comprehensive multi-omics profiling, including spatial proteomics, epigenomics, transcriptomics, and metabolomics. Integration of single-cell and spatial data enables the identification and mapping of diverse immune and stromal cell types—such as T cells, B cells, macrophages, dendritic cells, and endothelial cells—within the lymph node microenvironment. Spatial mapping of cellular senescence reveals the localization and organization of senescent cells, as well as the cellular composition and specialized niches of lymph node follicles. Multi-modal analyses further allow quantification of senescence signatures and intercellular communication, highlighting the heterogeneity of senescence states among distinct B cell subpopulations, including memory B cells, naïve B cells, and germinal center B cells.

**Figure 2.**
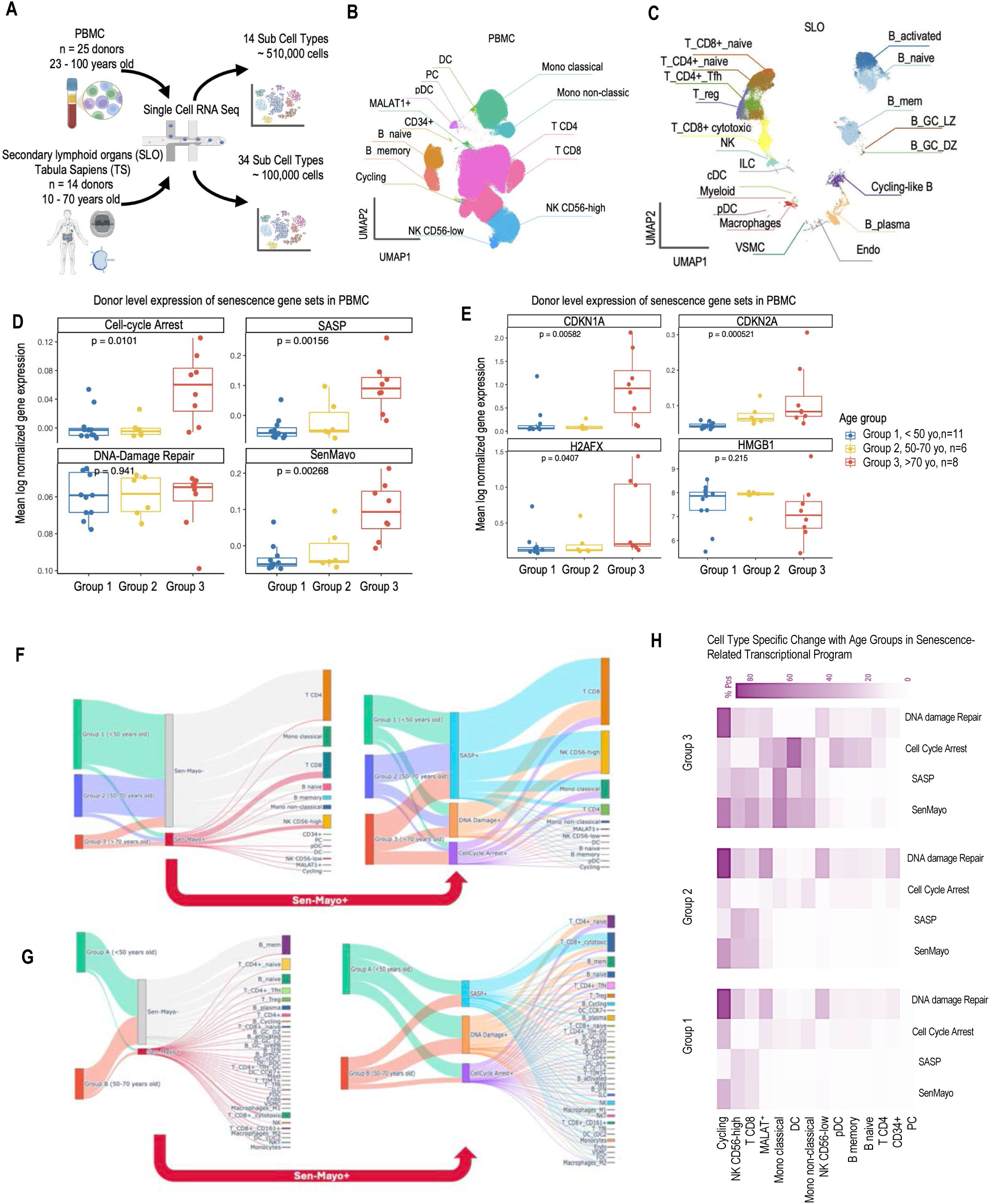
Single-cell transcriptome profiles from human immune cells and tissues across age groups. (A) Single-cell RNA sequencing (scRNA-seq) reference datasets for human PBMCs^19^ and lymphoid tissues curated from Tabula Sapiens (TS)^17^ and the integrated Secondary Lymphoid Organ (SLO)^18^ atlas. The PBMC collection includes 25 donors, and the SLO dataset includes 14 donors, stratified into Group 1: <50 years, Group 2: 50–70 years, and Group 3: >70 years for both sexes. (B & C) UMAP representations of the curated scRNA-seq reference datasets, showing distinct cell types in PBMCs and lymphoid tissues. (D & E) Boxplots showing mean log-normalized expression of senescence-associated gene sets in PBMC donors stratified by age group (Group 1: <50, Group 2: 50–70, Group 3: >70 years). (F & G) Sankey plots illustrating the contributions of age groups (PBMC: Group 1: <50, Group 2: 50–70, Group 3: >70; lymphoid tissue: Group A: <50, Group B: 50–70) to SenMayo⁻ and SenMayo⁺ populations (left) and to DNA Damage⁺, Cell Cycle Arrest⁺, and SASP⁺ compartments within the SenMayo⁺ population (right). Cells were classified as SenMayo⁺ if their SenMayo module score ranked within the top 20% of the dataset. Within the SenMayo⁺ subset, cells were further assigned as DNA Damage⁺, Cell Cycle Arrest⁺, or SASP⁺ if the corresponding module score ranked within the top 20% of scores computed for that program within the SenMayo⁺ population. (H) Heatmaps showing the percentage of cells positive for DNA damage repair, cell cycle arrest, SASP, and SenMayo signatures across PBMC cell types stratified by age group. Color intensity reflects the proportion of signature-positive cells within each cell type.

**Figure 3.**
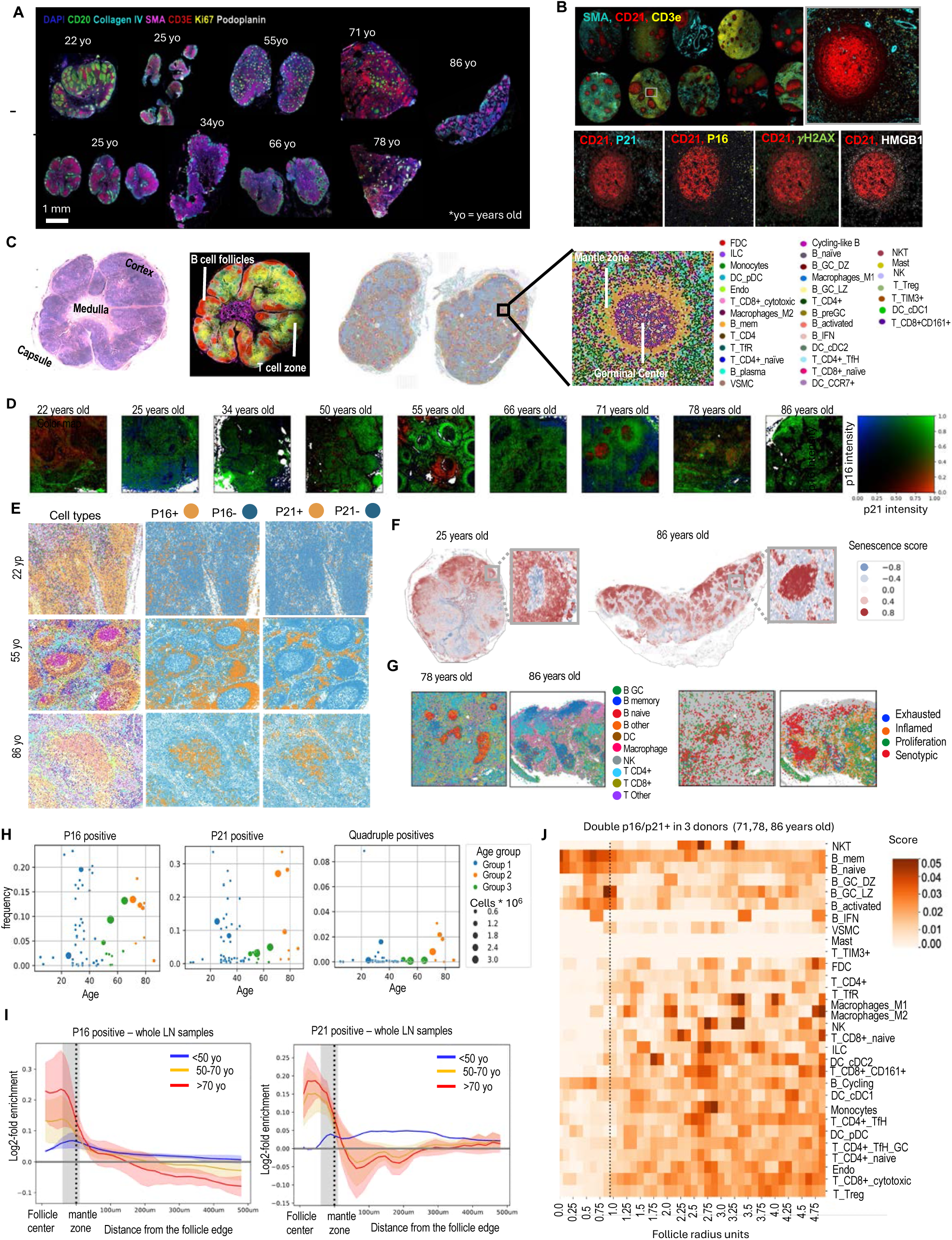
Spatial mapping of cell types and senescence via highly multiplexed immunofluorescence imaging. (A) Gallery of multiplexed immunofluorescence(mIF) images from nine whole lymph nodes, comprising 18 tissue sections, showing preserved anatomical structures. (B) Gallery of mIF images from selected cores in a human LN tissue microarray and zoomed-in view of a highlighted follicle showing senescence markers—HMGB1 (magenta), p16 (yellow), p21 (green), and *𝛾*-H2AX (red)—on a DAPI nuclear background (blue) and CD21 staining for follicles (red). p16 and p21 antibodies were validated using a p16⁺ nasopharyngeal carcinoma tissue section (Figure S3A). (C) Cell-type annotation of multiplexed immunofluorescence images using the MaxFuse pipeline identifies 34 distinct cell types. (D) Visualization of p16 and p21 expression across ages in zoomed-in follicular regions from individual donors. (E) Distribution of p16⁺ and p21⁺ cells within follicles across age groups (<50, 50–70, and >70 years). (F) Weighted senescence score (with greater weight assigned to p16) shown for representative young (25 years old) and aged (86 years old) samples. (G) Enlarged view of cell subtypes and their distinct functional states in a representative >70-year-old sample. (H) Quantification of p16 and p21 protein expression across all 111 tissue section samples, highlighting their spatial distribution and abundance change with age. (I) Radial distribution of p16⁺/p21⁺ cells relative to follicle radius, quantifying spatial localization patterns across age groups. (J) Heatmap of p16⁺/p21⁺ double-positive cells identifying the most enriched cell types within follicles in samples from donors >70 years old.

**Figure 4.**
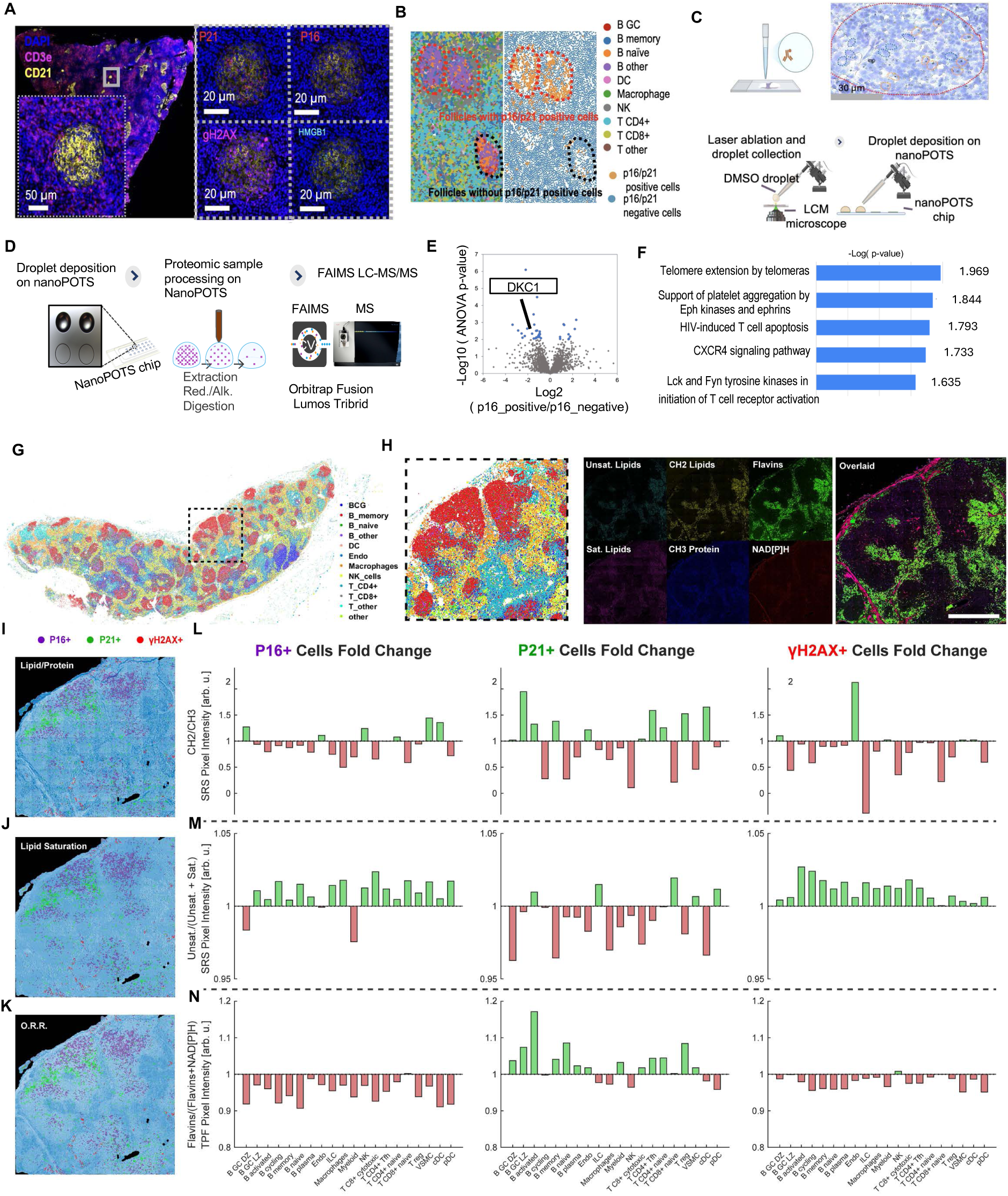
Spatial mass spec proteomics and optical metabolic imaging for characterization of cellular senescence. (A) Multiplex CODEX image of a 78 yo. human lymph node stained for CD3e (T-cells, red) and CD21 (follicular dendritic cells, yellow). Insets (right) show serial IF for the four core senescence markers p21, p16, *𝛾*-H2AX and HMGB1 within the boxed follicle (white box in overview). Scale bars, 50 µm (overview) and 20 µm (insets). (B) Cell-type map of the same node colored by single-cell annotation (legend, right). Follicles enriched (red dashed outline) or depleted (black dashed outline) for p16⁺/p21⁺ cells are highlighted; the accompanying binary map (right sub-panel) indicates the distribution of double-positive (orange) and double-negative (blue) cells. (C) Laser-capture/nanoPOTS isolation. Individual p16-classified cells were ablated from PEN-membrane sections (30 µm scale bar) and ejected into 200 nL DMSO droplets on a nanoPOTS chip for downstream proteomics. (D)Workflow for spatial single-cell proteomics: Single-cell ablation followed by droplet deposition on-chip protein extraction, reduction/alkylation, and tryptic digestion FAIMS LC-MS/MS on an Orbitrap Fusion Lumos Tribrid. (E)Volcano plot displaying differential protein abundance between p16⁺ and p16⁻ cells (ANOVA, FDR < 0.05). Blue dots denote proteins enriched or depleted in p16⁺ cells; DKC1 (boxed) is the most significantly down-regulated factor. (F) Top five Reactome pathways enriched among p16-regulated proteins (–log₁₀ P values shown). Telomere-extension machinery is most strongly impacted, concordant with DKC1 loss. (G) Whole-node segmentation (left) with zoomed view of a p16-rich germinal-centre cluster (right inset, dashed box). Cell-type colours as in panel B. (H) Label-free metabolic imaging of the boxed region in G using multiplex stimulated Raman scattering (SRS) and two-photon fluorescence (TPF). Channels: unsaturated lipids (C=C), CH₂ lipids, flavins, saturated lipids, CH₃ protein, NAD(P)H, and merged overlay. Scale bar, 100 µm. (I–K) Metabolic hallmarks stratified by senescence markers. Left, SRS map of the indicated ratio in the region shown in H (scale bar, 100 µm). Centre and right, cell-type-resolved bar plots for (i) lipid/protein ratio, (ii) SRS lipid-saturation ratio, and (iii) TPF optical redox ratio (ORR). (L-M) p16⁺, p21⁺ and *𝛾*-H2AX⁺ cells metabolic sates presented by bars representing log₂ fold change relative to lineage-specific means (green = increase, red = decrease); dashed lines denote no change.

### Human lymphoid tissue single-cell transcriptomics reference atlas

We obtained publicly available peripheral blood single-cell transcriptome datasets^19^ from 25 healthy donors (ages 23–100), retaining approximately 510,000 peripheral blood mononuclear cells (PBMCs) spanning 14 annotated immune subtypes, including NK CD56^high/low^ cells, classical and non-classical monocytes, plasmacytoid dendritic cells (pDCs), cycling progenitors, and naïve/memory B and T cells (Figure 2A,B; Figure S2A,B). We also analyzed single-cell RNA-seq data from human secondary lymphoid organs (SLOs) obtained from the Tabula Sapiens (TS) database. Visualization by UMAP resolved 34 distinct cell types (Figure 2A,C), comprising 11 B-cell subtypes, 4 dendritic cell (DC) subtypes, and 10 T-cell subtypes, in addition to NK cells, macrophages (M1 and M2), mast cells, and monocytes. Notably, T cells were further subdivided into cytotoxic and naïve populations, as well as functionally distinct helper T-cell subsets, including T follicular helper (T_FH) and T follicular regulatory (T_FR) cells (Figure 2C).

To quantify aging signatures, we performed differential gene expression analyses using the age brackets applied across the single-cell datasets. In lymphoid datasets, donors were grouped into Group A (<50 years) and Group B (50–69 years), whereas in PBMC datasets which included older donors, samples were stratified into Group 1 (<50 years), Group 2 (50–70 years), and Group 3 (>70 years) (Figure S2A,B). At the PBMC donor level, senescence-associated gene programs increased with age. Cell-cycle arrest, cytokine-inflammatory/SASP, and SenMayo scores were all significantly higher in Group 3 compared to Groups 1 and 2 (p ≈ 0.04, 0.001, and 0.0016, respectively). Similarly, canonical senescence markers CDKN1A (p21) and CDKN2A (p16) were elevated in Group 3 versus the younger groups (p = 0.0058 and 0.000521, respectively), consistent with increased cell-cycle inhibition and accumulation of senescent cells with age^20,21^ (Figure 2D,E). H2AFX and HMGB1 show a modest increase in the oldest group (p = 0.0407 and p = 0.215, respectively), consistent with age-associated elevation of DNA damage signaling, although the effect size remains moderate, likely reflecting cellular heterogeneity. Notably, senescence-related HMGB1 biology is often mediated by protein relocalization and extracellular release rather than by large transcriptional changes, making such effects less apparent at the RNA level ^22,23^. Pathway-level Principal Component Analysis (PCA) further demonstrated that aging-associated variation is dominated by inflammatory and stress-response programs. TNF*α*/NF-kB signaling, inflammatory response, p53 signaling, unfolded protein response, ROS, and apoptosis pathways showed as the key altered pathways in older donors (Figure 2C). In contrast, proliferative and cell-cycle–associated pathways (E2F targets, G2M checkpoint, mitotic spindle, DNA repair) loaded on the opposing axis for the younger donors, suggesting a reorganization of cycling compartments with age (Figure 2C). Differential-expression analysis between the groups of SLO and PBMC donors identifies the heaviest age-associated transcriptomic burdens (≥ 3,000 DE genes). Across both datasets, subtypes of B cells including transcriptionally-defined proliferative B-cell pools (i.e, cycling-like B cells and B_plasma-cells) exhibit alterations with age presumably corresponding to the change in antibody production and immune response^24^ (Figure S2D,E).

The Sankey plots illustrate how different cell types contribute to the SenMayo*⁺* versus SenMayo*⁻* compartments across age groups (Figure 2F,G). In PBMCs, the majority of cells from all age groups remain in the SenMayo*⁻* pool; however, the relative contribution to the SenMayo*⁺* fraction increases with age, with donors ≥70 years contributing disproportionately to senescence compared to younger groups. These SenMayo*⁺* cells are primarily localized to CD4*⁺* and CD8*⁺* T cells, NK CD56^high^ cells, and classical monocytes (Figure 2F). In SLO, most cells from both age groups feed predominantly into SenMayo*⁻*, but Group B shows a larger proportional flow into SenMayo*⁺* (Figure 2G). Within this fraction, enrichment is observed in memory, naïve, and plasma B cells, as well as T-cell subtypes including T follicular helper (T_FH), activated CD4*⁺*, and CD8*⁺* T cells (Figure 2G). Across both PBMC and SLO datasets, MALAT1*⁺* stromal cells exhibit an age-associated increase in SenMayo activity, with the oldest donors contributing most prominently to the SenMayo*⁺* fraction, consistent with a late-life stromal senescence trajectory (Figure 2F–H). In the lymphoid tissue microenvironment, this pattern extends beyond MALAT1*⁺* stromal cells: additional stromal and vascular compartments—including follicular dendritic cells (FDCs), vascular smooth muscle cells (VSMCs), and endothelial cells—also display measurable SenMayo*⁺* contributions. These findings support the conclusion that senescence in lymphoid tissues involves not only hematopoietic cells but also broader remodeling of the tissue niches.

Aging signatures in the PBMC dataset are markedly elevated in Group 3 (≥70 years), with the most pronounced shifts observed in myeloid and cycling compartments (Figure 2H). Classical and non-classical monocytes, as well as dendritic cells (DCs), exhibit strong enrichment for SenMayo and SASP programs, accompanied by a notable increase in cell-cycle arrest, particularly in DCs. Cycling-like B cells (a.k.a., cycling in Figure 2H) consistently display high DNA-damage signals across all age groups, remaining among the most enriched populations overall (Figure 2H).

The single-cell RNA-seq data from secondary lymphoid organs (SLOs) and peripheral blood mononuclear cells (PBMCs) indicate that immunosenescence is relatively modest in donors under 50 and those between 50–70 years, but becomes pronounced around 70 years of age. Donors over 70 exhibit aging-associated immune remodeling across multiple cell subtypes. In PBMCs, this is reflected by a pronounced shift toward inflammatory and SASP-associated programs, as well as an expanded SenMayo*⁺* fraction, particularly in CD4*⁺* T cells, CD8*⁺* T cells, and NK cells. In contrast, SLO single-cell data suggest that senescence-like states in lymphoid tissue B cells are present in older donors that may also contribute to age-related impairments in humoral immunity—a phenomenon that remains understudied but is crucial for adaptive immune responses. Understanding these changes in SLO B cells is therefore a focus of this work.

### Functional heterogeneity of cells enriched for SenMayo across age groups

The top 10% of cells enriched for the SenMayo gene set were selected from the single-cell lymphoid tissue datasets and subsetted into clusters (Figure S3A,B). The genes highlighted are primarily involved in the senescence-associated secretory phenotype (SASP) and other pathways linked to cellular senescence and inflammation^20^. SASP factors, including CXCL3, CXCL2, CXCR2, IL10, and IL1B, are prominently expressed in cluster 2 (Th1/Th17-like)^25–29^. Clusters 4 and 5, corresponding to vascular endothelial cells and fibroblasts, exhibit elevated expression of GDF15 and members of the insulin-like growth factor-binding protein (IGFBP) family—IGFBP2, IGFBP3, IGFBP4, and IGFBP5—highlighting altered growth and apoptosis signaling in non-immune stromal compartments^30–33^. Clusters 3 and 4, marked by strong ICAM1 expression, may facilitate immune cell recruitment and adhesion, contributing to the inflammatory response associated with tissue senescence^34^.

The ligand–receptor map of lymphoid tissue donors aged >50 reveals that SASP-enriched cells reside within a highly interconnected signaling hub coordinating immune evasion, leukocyte recruitment, and checkpoint regulation (Figure S3C). High-probability LGALS9-CD45 signaling from migratory dendritic cells (cDC1/cDC2/CCR7*⁺*) may enforce Galectin-9–mediated checkpoint suppression of T and NK cell activation. Concurrently, widespread CLEC2C-KLRB1 (LLT-CD161) interactions across NK and CD161*⁺* T cells suggest a counterbalancing role in maintaining innate immune surveillance within the SASP niche^35,36^. Within the B cell compartment, a prominent CD22-PTPRC (CD45) axis, most pronounced in naïve, memory, activated, and germinal center B cells (both dark and light zones), provides ITIM-mediated inhibition of B cell receptor signaling, likely modulating responses in the inflammatory milieu□^37^. Endothelial and stromal activation is supported by significant APP-CD74 signaling, which promotes NF-kB activation^38^. Additionally, an extensive network of ICAM1/2-ITGAL/ITGB2 (LFA-1) interactions spans nearly all T, B, NK, monocyte, and dendritic cell populations, probably facilitating proinflammatory cytokine production (Figure S3C)^39^.

Pathway enrichment analysis reveals age-dependent shifts in cell–cell communication networks. In donors under 50, the CD99 signaling module is significantly enriched (Figure S3D)^40^. In contrast, donors over 50 show selective enrichment of the Semaphorin-4 (SEMA4) pathway, which promotes activation of B cells and dendritic cells (Figure S3D)^41^. In the younger group, SASP cells influence TIM3*⁺* dendritic cells (DCs) and germinal center T follicular helper (GC T_FH) cells through the ICAM pathway (Figure S3D). In older donors, the ICAM network exhibits increased connectivity among macrophages (M1 and M2), monocytes, and regulatory T cells (Tregs), reflecting interactions between immune-regulatory and innate immune populations and a shift toward immune suppression and “inflammaging”^10^. It has been shown that Age-associated B cells (ABCs)^42,43^, characterized by T-bet (encoded by TBX21), serve as hallmarks of immune alteration in aging and are linked to chronic inflammation and autoimmunity. In older individuals, T-bet*⁺* ABCs show upregulation of SenMayo genes, particularly in cycling-like B cells and plasma B cells. Moreover, elevated CDKN2A and H2AFX expression in cycling-like B cells and germinal center dark-zone B cells (B_GC_DZ) identifies these subtypes as major populations affected by senescence-like programs, which differs from classical ABCs and is consistent with spatial analyses (Figure S3E).

Together, these analyses highlight inflammatory genes and SASP-associated pathways, while also revealing senescence-related programs in proliferative populations such as cycling-like B cells and germinal center dark-zone B cells (B_GC_DZ). Age-stratified differential expression further indicates that B-lineage cells—particularly naïve, memory, cycling-like, and plasma B cells in lymphoid tissues—carry some of the greatest transcriptional burdens associated with aging. While it remains debated whether these proliferative cells should be classified as fully senescent, it is well known that immunosenescence is not defined strictly by cell-cycle arrest, but rather by age-associated declines in immune function, in part driven by senescence-related programs.

### High-plex protein imaging for mapping region- and type-specific cellular senescence

A 53-plex CODEX (CO-Detection by indEXing) assay was performed on human whole lymph node sections and two tissue microarrays (YTMA and CTMA) using an antibody panel that included cell type or state–specific markers such as CD20 (B cells), CD3e (T cells), Ki67 (proliferating cells), Podoplanin (lymphatic endothelial cells), and SMA (smooth muscle actin), together with additional markers including four senescence markers (Figure 3A,B). This panel enabled examination of follicular architecture as well as age-associated functional states of cells, including reduced proliferation and increased exhaustion in older donors (Figure S5A,B). Each of the four senescence markers (p16, p21, HMGB1, and *𝛾*-H2AX) reports on a distinct senescence-associated pathway. Across multiple follicles in lymph node sections and TMAs from young (18–50 years old) to mid-aged (50–70 years old) donors, cells positive for p21 and HMGB1 were observed primarily at the periphery of follicles, whereas *𝛾*-H2AX- and p16-positive cells were distributed across both follicular peripheral zones and interfollicular regions. Notably, relatively few p16-positive cells were detected (Figure 3B). To quantify cell type–specific senescence, we integrated single-cell transcriptomic reference data with CODEX imaging data using MaxFuse^44^, enabling the transfer of annotations for 34 major cell types (Figure 3C, S5E). These included key immune populations such as naïve, memory, and activated B cells; CD4*⁺*, CD8*⁺* cytotoxic, and regulatory T cells; M1/M2 macrophages; plasmacytoid and conventional dendritic cells; as well as stromal and other immune cell types, including endothelial cells, monocytes, and NK cells. Multiplexed CODEX imaging of senescence markers revealed increasing co-expression of p16 and p21 with age (Figure 3D,E). In individuals aged 50–70 years, these markers were predominantly localized in the mantle zone, whereas in the 86-year-old sample they shifted toward the follicles, suggesting altered germinal center (GC) B cell states during aging (Figure 3E, S5C, S7A). To quantify spatial senescence patterns, we computed a senescence score based on the spatial expression of p16, p21, HMGB1, and *𝛾*-H2AX. This analysis revealed distinct age-associated distributions: younger samples (e.g., 25 years old) exhibited higher scores in interfollicular regions, whereas older samples (e.g., 86 years old) showed elevated scores within follicles (Figure 3F, S5B). Integrating the cell type map with spatial distributions of functional states enabled identification of cell populations acquiring senescent phenotypes with age (Figure 3G). Notably, senescent-like cells aggregated within follicles of older samples and displayed pronounced spatial heterogeneity across lymph node regions (Figure S5A,D). Marker-positive populations—including single-positive (e.g., p16*⁺*), double-positive (p16*⁺*p21*⁺*), triple-positive (p16*⁺*p21*⁺𝛾*-H2AX*⁺*), and quadruple-positive (p16*⁺*p21*⁺ 𝛾*-H2AX*⁺*HMGB1*⁺*) cells—all showed significant increases in older donors (Figure S6A,B). Notably, some senescence markers were also detected in younger individuals, potentially reflecting transient cellular stress responses associated with conditions such as acute infection or autoimmune activation. We further mapped spatially distinct immune cell states, including exhausted, inflamed, proliferating, and senescent-like (“senotypic”) phenotypes (see Methods). Older samples exhibited greater accumulation of senescent-like and exhausted-like cells, consistent with the notion of a shift toward non-proliferative and functionally impaired immune populations during aging (Figure S5A). Age-stratified analyses confirmed a marked increase in senescence marker frequency across samples, with a pronounced rise in p16*⁺* and p21*⁺* cells in donors over 70 years old and quadruple-positive cells observed almost exclusively in this age group (Figure 3H, S6A,B).

While vascular and stromal cell senescence has been documented during aging, whether key immune populations such as follicular B cells undergo senescence—and with what consequences—remains poorly understood. To quantify the spatial distribution of senescence across cell types, we identified lymphoid follicles in each sample using canonical markers. Cells with high CD20 and CD21 expression (top 10–25%) were designated as mature B cells of interest in these lymph node samples, with thresholds adjusted based on sample quality. Each follicle was modeled using a single-component Gaussian mixture to derive its centroid (µ) and covariance matrix (□), capturing its shape and orientation. We then quantified radial abundance of senescence markers using two complementary metrics: Mahalanobis distance within follicles to account for their geometry, and Euclidean distance outside follicles to assess marker localization in surrounding tissue. Ten Mahalanobis-based annuli within one follicular radius and 25 Euclidean annuli spanning 1–500 µm from the follicle center (both ∼20 µm wide) were used. Fold-enrichment in each annulus was calculated as the density of marker-positive cells relative to background, normalized by area and stabilized with pseudocounts. This dual-metric framework enabled comparison of intrafollicular and peripheral senescence patterns across variable follicle morphologies (Figure S7A). In 18–50 years old lymph nodes, p16- and p21-positive cells were broadly distributed, extending into the paracortex and forming a wide dome-shaped profile. By 50 years old, the peak contracted toward the germinal center and mantle zone. In >70 years old donors, the signal became concentrated within the follicular core, with surrounding regions near background, indicating focal accumulation of senescent-like cells in germinal centers (Figure 3I, S6B). This centripetal shift was mirrored in the double-positive map, where >70 years old samples showed a sharp central peak (Figure 3J). Collectively, these data reveal an age-dependent increase in senescence-marker frequency and a progressive spatial shift from diffuse distribution in youth to germinal center confinement in >70 years old samples. Spatial heatmaps of p16*⁺*/p21*⁺* double-positive cells further confirmed that in >70 years old lymph nodes, senescence is largely restricted to the follicular compartment (x-axis < 1.0, dashed line; Figure S6A). The strongest p16*⁺*/p21*⁺* signal occurred in B-lineage subsets—including memory, naïve, GC (DZ and LZ), activated, and IFN-responsive B cells—reaching 3–5% of the local cell pool. Beyond the follicle (radius > 1.0), double-positive cell density declined sharply across all lineages, indicating that age-related immunosenescence in lymph nodes is follicle-centered and primarily affects antigen-experienced and proliferative B-cell populations (Figure 3J, S7A). Spatial protein mapping of aged (>70 y.o.) lymph node follicles showed that HMGB1 was the most highly expressed—though likely less specific—senescence marker across B cell subtypes, followed by *𝛾*-H2AX, p21, and p16. Co-expression of multiple markers, including triple-positive (p16*⁺*p21*⁺ 𝛾*-H2AX*⁺* or p16*⁺*p21*⁺*HMGB1*⁺*) and quadruple-positive (p16*⁺*p21*⁺ 𝛾*-H2AX*⁺*HMGB1*⁺*) profiles, was largely confined to B_memory and B_naive cells, suggesting that these subsets harbor the most advanced or multifaceted senescence signatures in aged follicles (Figure S7B).

Here we observed a pronounced age-related increase in the expression and co-expression of key senescence markers (p16, p21, *𝛾*-H2AX, and HMGB1), particularly within follicular B cell compartments. With aging, senescent-like cells shift from a relatively diffuse tissue-wide distribution to concentrated hotspots within germinal centers, reflecting a centripetal remodeling of the senescent niche. This spatial reorganization predominantly involves antigen-experienced and proliferative B cell subsets—including memory, naïve, germinal center dark-zone, and germinal center light-zone populations—suggesting that senescence-like programs may increase their vulnerability to age-associated B cell dysfunction, for example in responses to infection or vaccination.

### Spatial proteomic and metabolomic characterization of cellular senescence

Using high-sensitivity mass spectrometry for single-cell proteomic profiling of p16*⁺* lymph node cells could uncover senescence-associated protein networks and post-transcriptional modifications that underlie immune cell senescence during aging. On a fresh frozen biopsy obtained from a 78-year-old donor, spatially resolved laser-capture isolation of individual p16 positive cells in follicles followed by nanoPOTS single-cell proteomics yielded a median of ∼740 protein groups per cell^45,46^ (Figure 4C-D, S8A, B). Comparative analysis of p16*⁺* versus p16*⁻* follicular cells identified 150 proteins with significant differential abundance (ANOVA FDR < 0.05), among which the telomerase co-factor Dyskeratosis congenita 1 (*DKC1*) was one of the most strongly depleted in the senescent cohort (log*₂*FC ≈ -3.2; Figure 4E). *DKC1* (log*₂*FC ≈ -3.2; Figure 4E), a core component of the H/ACA telomerase complex, indicating compromised telomere-maintenance machinery in senescent cells. Its depletion shows the senescent state of the p16+ follicular cells^47,48^. Additional downregulated proteins included EIF3K, RPL3/RPLP1B, and RPRD2, consistent with suppression of translational initiation and RNA polymerase II–associated transcriptional regulation^49,50^ (Figure S7B). SFXN1 reduction further suggested impaired mitochondrial serine transport and one-carbon metabolism^51^(Figure S7B). Conversely, proteins enriched in p16*⁺* cells included RAB1B, associated with ER stress and autophagosome formation, and MARC2, a mitochondrial enzyme linked to redox regulation^52^. Pathway enrichment analysis identified “Telomere extension by telomerase” as the top altered pathway, alongside CXCR4 signaling and Lck/Fyn-mediated T-cell receptor initiation (Figure 4F), collectively indicating telomere instability, stress adaptation, and rewired immune-receptor signaling (Figure 4F). Collectively, unbiased proteomic profiling could unveil molecular mechanisms of cellular senescence in human lymph nodes such as telomere-stabilizing deficits alongside rewired immune-receptor signaling, reinforcing the findings from spatial mapping that link p16 positivity to potential functional decline and chronic inflammatory signaling in aged human lymphoid tissue.

To determine whether the changes observed in p16*⁺* cells translate into functional metabolic alterations in situ, we applied stimulated Raman scattering (SRS) metabolic microscopy with two-photon autofluorescence (TPF) and second harmonic generation (SHG) imaging ^53–55^ to this tissue sample. Following SRS imaging, the same tissue section was stained and imaged with CODEX for cell typing^56,57^. Co-registration of CODEX and SRS data enabled correlation of metabolic profiles with specific cell types (Figure 4G,H). Ratiometric images revealed distinct metabolic patterns in cells positive for senescence markers (Figure 4I–K). By intersecting senescence marker-positive cells with cell type identities, we quantified the proportion of each cell type within different senescence classes (senotypes). Fold changes in metabolic indicators between senescence marker–positive and –negative cells were calculated across cell types (Figure 4L–N), and the mean metabolic indicator values for all cells in the region of interest are plotted in Figure S8D. We observed that the lipid*-*to*-*protein ratio was significantly elevated in p16*-* and p21*-*positive GC B cells (B_GC). By comparison, endothelial cells exhibited only modest lipid/protein increases in p16*-* or p21*-*positive cells, but a pronounced rise in *𝛾*-H2AX*-*positive cells (Figure 4L). As expected for senescent populations, the lipid*-*saturation ratio (CH*₂*/CH*₃*) increased in most p16*-* and *𝛾*-H2AX*-*positive cells. *𝛾*-H2AX*-*positive cells, which localize predominantly to the perifollicular or paracortical zone, displayed the most consistent and pronounced rise in lipid saturation (Figure 4M). The optical redox ratio (an indicator that typically falls as cells enter a more oxidized, senescent state) was indeed lower in p16*-* and *𝛾*-H2AX*-*positive populations with a clearer pattern compared to lipid*-*to*-*protein and lipid*-*saturation ratio due to its clear profile in follicular zones. P21*-*positive cells did not exhibit this decline, likely because they cluster predominantly in the interfollicular (perifollicular) zone, where many cells remain highly proliferative and therefore sustain a more reduced redox balance (Figure 4N).

These findings underscore that senescence marker expression is highly cell type–specific within the lymphoid compartment, as well as senescence marker dependent. B-memory cells emerge as the predominant senescent population, accounting for the majority of p16*⁺* (∼75%) and *𝛾*-H2AX*⁺* (∼60%) cells, suggesting they are a plausible reservoir of advanced or irreversible senescence (Figure S8C, D). This division implies that different senescence markers capture distinct senotypes as well as stages, trajectories, or mechanisms of cellular aging, with B cells occupying different positions along the senescence continuum.

### Spatial whole transcriptome sequencing for characterization of cellular senescence

Spatial whole-transcriptome sequencing was performed on both FFPE and fresh-frozen (FF) human lymph node tissues from 11 donors, focusing on selected regions of interest to investigate the mechanisms underlying cellular senescence using DBiT-seq^16^. Unsupervised clustering revealed distinct transcriptomic clusters in each sample, corresponding broadly to major cell types and anatomical regions, including the cortex, vessels, and B and T cell zones (Figure 5A). Spatial UMAP projections closely aligned with histology or CODEX images of adjacent sections, enabling precise spatial mapping of cell type–specific transcriptomic signatures and cluster annotation based on canonical cell-type markers (Figure S9A). Notably, the B cell cluster within primary follicles of the 86-year-old donor exhibited strong enrichment for multiple senescence-associated pathways, including NF-kB signaling, p53 signaling, and cell cycle checkpoints, with moderate enrichment for DNA damage response (DDR) and SASP pathways (Figure S9A). Using iStar, we generated super-resolved transcriptome-wide expression maps by integrating spatial transcriptomics with high-resolution tissue histology, achieving near–single-cell resolution of gene expression across entire tissue slides (Figure S9B)^58^. Comparison with protein-level expression measured by CODEX on serial sections revealed that markers such as p21 and HMGB1 localized within follicular peripheries but also dispersed across broader tissue areas at the transcriptomic level. In contrast, p16 protein expression was more robust and widespread than its transcript-level counterpart, which appeared sparse (Figure S9B). HMGB1 displayed strong protein-level enrichment in peripheral regions, consistent with its transcriptomic distribution. These observations highlight the heterogeneity of senescence profiles with aging, emphasizing the increased prominence of *𝛾*-H2AX- and p21-driven senescence in tissues from donors over 70 years old.

**Figure 5.**
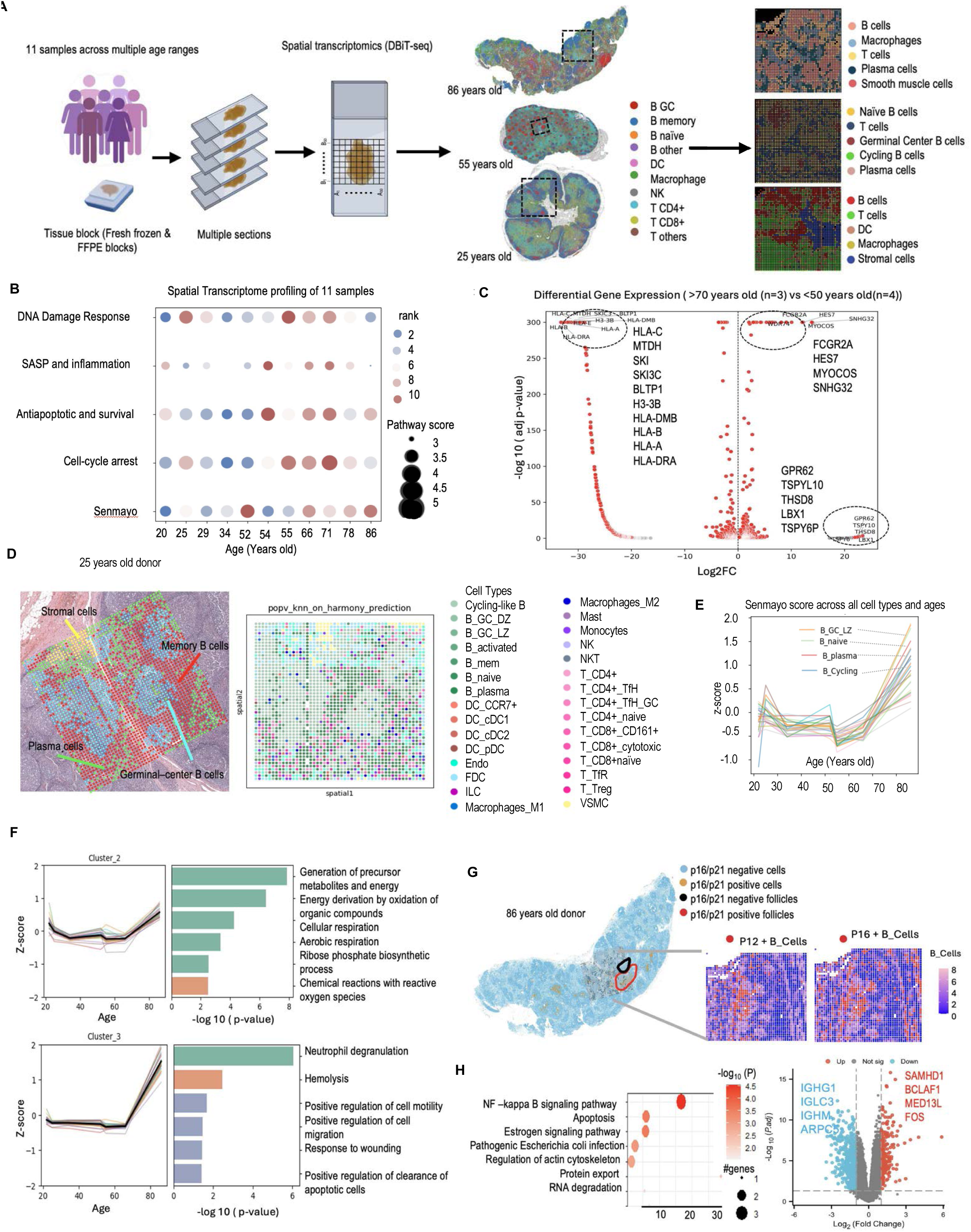
Spatial transcriptomic atlas of cellular senescence in human lymph nodes. (A) Schematic overview of spatial transcriptomic profiling using DBiT-seq performed on 11 human lymph node samples spanning ages 20 to 86 years. Representative spatial maps from 25-, 55-, and 86-year-old donors are shown, with cell types annotated based on single-cell reference datasets. (B) Bubble plot showing pathway scores for key senescence-associated processes—including DNA damage response, SASP/inflammation, anti-apoptotic/survival signaling, cell-cycle arrest, and the SenMayo signature—across all profiled samples. Scores are ranked by magnitude for each sample and pathway. Increased anti-apoptotic/survival signaling and SenMayo activity are evident in samples from donors older than 70 years. (C) Volcano plot showing genes differentially expressed in lymph node samples from donors >70 years versus <50 years, ranked by log₂ fold change (log₂FC). Red points indicate genes with an adjusted p-value < 0.05. (D) High-resolution cell type annotation of a representative mid-age (55-year-old) lymph node section, visualized at both low and high magnification. Annotation leverages single-cell reference data to resolve 34 cell types, highlighting the spatial localization of germinal center B cells, memory B cells, stromal cells, and plasma cells. (E) Line plot showing the kinetics of SenMayo gene set z-scores across age among 34 annotated cell types. Individual cell-type trends (colored lines) and the global mean (black line) demonstrate a marked increase in senescence signature expression with advancing age, particularly after 70 years. (F) Genes were clustered according to their expression trajectories across age across all cell types. Left panels show mean z-score trajectories across age for each cell type (colored lines) with the global mean in black. Right panels display the top enriched pathways (□log₁₀ p-value), colored by database (green: GO Biological Process; orange: KEGG; blue: Reactome). Both clusters show a pronounced increase in expression during late life, particularly after age 70. (G) Spatial mapping of p16 and p21 double-positive and double-negative B cells within a follicle, integrating high-dimensional proteomic (CODEX) and spatial transcriptomic (DBiT-seq) data from an 86-year-old donor. Insets show magnified views of B-cell regions highlighting p16/p21 expression status. (H) Volcano plot and pathway enrichment analysis showing differentially expressed genes and key pathways distinguishing p16/p21 double-positive versus double-negative B cells within the DBiT-seq region of interest.

The average expression of genes from key senescence-associated pathways was assessed across all 11 samples, spanning donors aged 20 to 86 years (Figure 5B). Among the pathways examined, the SenMayo signature exhibited the most pronounced age-related pattern, showing consistently higher activity in samples from donors ≥66 years, with the exception of one 52-year-old sample that displayed relatively low activity across multiple pathways. Other pathways, including DNA damage response, cell-cycle arrest, and SASP/inflammation, also tended to be elevated in donors ≥66 years, albeit with greater variability. Given that spatial transcriptomic measurements capture multiple cell types across distinct anatomical regions (Figure 5A), this pathway activity profiles likely represent a composite of cell-type–specific and architecture-dependent signals. To assess the effects of age, senescence-pathway activity was compared between younger and older groups using a cutoff of 53 years. A pathway was considered active (True) if its score exceeded the cohort median. Older samples exhibited a greater number of active senescence-associated pathways per sample (mean 3.33) compared with younger samples (mean 1.67), a difference that was statistically significant (one-sided Mann–Whitney U test, U = 30.0, p = 0.029) (Figure S9C). Notably, this result remained consistent after removing overlapping genes between the SASP/inflammation and SenMayo signatures, indicating that the observed trend was not driven by gene-set redundancy. Differential gene expression between the two age groups was visualized using a volcano plot (Figure 5D). Multiple HLA class I and II genes (*HLA-A, HLA-B, HLA-C, HLA-DRA, HLA-DMB*) were among the most significantly upregulated in samples from donors older than 53 years, reflecting age-associated immune remodeling. Additional genes, including *FCGR2A* and *HES7*, were also elevated in older samples, whereas a smaller subset of genes (*GPR62, TSPYL10, THSD8,* and *LBX1*) was relatively enriched in younger tissues (Figure 5D).

### Cell-type-specific transitional programs in senescence

Annotation of cell types in the spatial transcriptomic data was done using the MaxFuse algorithm to integrate the spatial dataset with our lymphoid tissue single-cell reference dataset^59^. This integration enabled more comprehensive cell-type annotations across the spatial landscape of the tissue (Figure 5D). Using the integrated cell and gene matrix, we assessed the enrichment of senescence markers to identify cell types exhibiting transcriptomic senescence signatures (Figure 5F). Cell type–resolved analysis revealed a progressive, age-dependent increase in *CDKN2A* (p16), *CDKN1A* (p21), *H2AX*, and *HMGB1* expression in naïve, plasma, and germinal center B cells, as well as in vascular smooth muscle cells, endothelial cells, T follicular helper cells, and dendritic cells (Figure S10A). These results reflect both the heterogeneity and tissue-wide accumulation of senescence and stress signatures in aging lymphoid tissues. Integrative analysis of the SenMayo gene set across annotated cell types showed a pronounced rise in senescence programs after 60 years of age, with a sharp increase in z-scores in nearly all cell types among individuals over 70 (Figure 5E). Notably, although most lymph node cell types exhibited significant age-associated enrichment for the SenMayo signature—including B_GC_LZ, B_naive, B_plasma, and cycling-like B cells—other pathways, such as DNA damage response, cell cycle arrest, anti-apoptotic signaling, and SASP, displayed more variable patterns (Figure S10C).

To map age-associated gene expression dynamics and functional remodeling in human lymph nodes, we clustered genes according to their expression trajectories across the aging continuum and performed pathway enrichment analysis for each cluster (Figure 5F, S10C). Clusters 0–1 captured signatures related to age-related declines: Cluster 0 exhibited stable-to-decreasing expression and was enriched for classical complement activation and translational programs, including peptide chain elongation and viral mRNA translation, while Cluster 1 showed a sharp early-life drop followed by a plateau and was enriched for RNA processing and splicing, consistent with reduced mRNA maturation with age (Figure S10C). Clusters 2–3 captured signatures related to late-age reprogramming: Cluster 2 displayed a modest early decline followed by recovery in older donors and was enriched for energy metabolism and precursor metabolite generation, whereas Cluster 3 was selectively upregulated in late life and enriched for pathways related to neutrophil degranulation, ferroptosis, cell motility, and wound response. Collectively, these patterns indicate metabolic adaptation and the emergence of inflammatory and tissue-remodeling transcriptional programs during lymph node aging (Figure 5F).

### Pathway analysis in B cells of >70-year-old lymph node follicles

Pathway enrichment analysis of B cell clusters from spatial transcriptomics of an 86-year-old lymph node revealed pronounced age-associated transcriptional shifts. Most significantly enriched pathways were upregulated and involved immune activation, cytokine signaling, NF-□B activation, and stress responses. Focusing on B cells within primary follicles using CODEX and DBiT-seq (Figure 5G, S9E), we identified distinct p16/p21-positive and -negative populations. Differential expression analysis showed that p16/p21-negative B cells expressed higher levels of genes such as *ATF4, TMEM154, CANX,* and *ELK4*, which are associated with active cellular states, immune regulation, protein homeostasis, and cell cycle progression within the lymphoid microenvironment (Figure S9E)^60,61^. Several SASP-related genes show higher expression in p16/p21-positive B cells, indicative of the senescent phenotype including *CD63, MAVS, IL6ST, CXCR4, CYTH1* and *KTN1*^62,63,64,65^. The volcano plot shows differential gene expression between p16/p21 double-positive and double-negative B cells in primary follicles of an 86-year-old lymph node (Figure 5H). Senescent (double-positive) B cells upregulate genes like *SAMHD1, BCLAF1, MED13L*, and *FOS*, linked to cell cycle control, stress responses, and immune signaling. Conversely, genes such as *IGHG1, IGLC3, IGHM,* and *ARPC5* are downregulated, indicating reduced antibody production and cytoskeletal activity (Figure 5H). The pathway enrichment plot highlights multiple biological pathways enriched in p16/p21-positive B cells, reflecting their senescent phenotype, with NF-kB signaling – key to inflammation, immune responses, and cellular stress – showing the highest fold enrichment^66,67^. Apoptotic pathways were also upregulated during senescence as part of the response to stress and damage^68^. To further characterize cellular senescence at the single-cell level, we used the 6,000-gene CosMx panel to identify p16- and p21-positive cells across major cell types (Figure S9D), which revealed heterogeneity in senescence marker expression, with z-scores indicating specific enrichment within follicular structures. Focusing on B cells, we aligned the CosMx data with CODEX multiplex immunofluorescence using STalign, identifying senescent cells based on their p16/p21 protein profiles^69^ (Figure S9D). Differential gene expression analysis between senescence-adjacent and distant B cells identified differential expression of key SASP-related genes, including *CCL21*, *COX1*, and *COX2* ^70,71^

### Spatial epigenomics sequencing for characterization of cellular senescence

Epigenomic alterations are also central to the development of cellular senescence, involving DNA damage–induced changes in chromatin accessibility, DNA methylation patterns, etc. These changes are also closely linked to age-related diseases and have informed the development of epigenetic clocks as biomarkers of biological aging. Thus, spatial epigenomics may capture early events or onset of senescent-like cellular states^72,73^. Global chromatin landscape changes associated with cellular senescence were profiled in nine fresh-frozen lymph node (LN) tissues spanning ages 20 to 86 using spatial ATAC-seq, enabling spatially resolved epigenomic mapping within LN tissues (Figure 6A, S11A). A representative sample from a 54-year-old donor showed unsupervised clustering corresponding to major histological structures, delineating B and T cell regions. Integration of spatial ATAC-seq data with a single-cell reference atlas (Figure 2) generated a UMAP representation with distinct B- and T-cell subtypes (Figure 6B, C). Chromatin-based gene activity scores indicated high transcriptional potential in B- and T-cell zones, consistent with IHC staining for CD20 and BCL2 on adjacent sections. Accessibility of PAX5 and CD3D was spatially localized to B- and T-cell regions, respectively (Figure 6D). Differentially accessible regions (FDR < 0.05, log*₂*FC ≥ 0.25) revealed senescence-associated chromatin changes across distinct histological regions (Figure 6E). Notably, gene activity scores for key senescence markers—including *CDKN1A, CDKN2A, H2AFX,* and *HMGB1*—were enriched in B-cell and activated T-cell zones, indicating increased accessibility of senescence-associated loci during aging (Figure 6E). We next predicted regulatory interactions between promoter regions and candidate enhancers through peak-calling–based analysis, identifying dynamically regulated promoter–enhancer interactions associated with key senescence genes, including *CDKN2A* (FDR < 0.05, log*₂*FC ≥ 0.1), across spatial clusters, highlighting the potential regulatory role of p16 in driving cellular senescence (Figure 6F). Motif enrichment analysis was then applied to identify transcription factor (TF) binding motifs associated with differential chromatin accessibility and gene activity scores across clusters (Figure 6G). The motif enrichment heatmap revealed distinct, cell type–specific regulatory programs relevant to senescence. Fibroblast, B-cell follicle, and endothelial clusters showed enrichment of developmental and stress-responsive TF motifs, including *HOXA7, NFIX, SMAD4, NFE2L1*, and *MEIS1*, suggesting activation of differentiation and stress programs within structural and stromal compartments. In contrast, activated T cells exhibited stronger enrichment of immune activation–associated motifs such as *SPIB* and *EGR3*, consistent with potential SASP-related signaling. Top-ranked motifs associated with transcription factors regulating *CDKN1A, CDKN2A, HMGB1*, and *H2AFX* included *ZKSCAN5, FOS:JUN, TCFL5, FOSL2:JUN, FOSL1:JUNB, NHLH2, FOS:JUND*, and *ATF6*.

**Figure 6.**
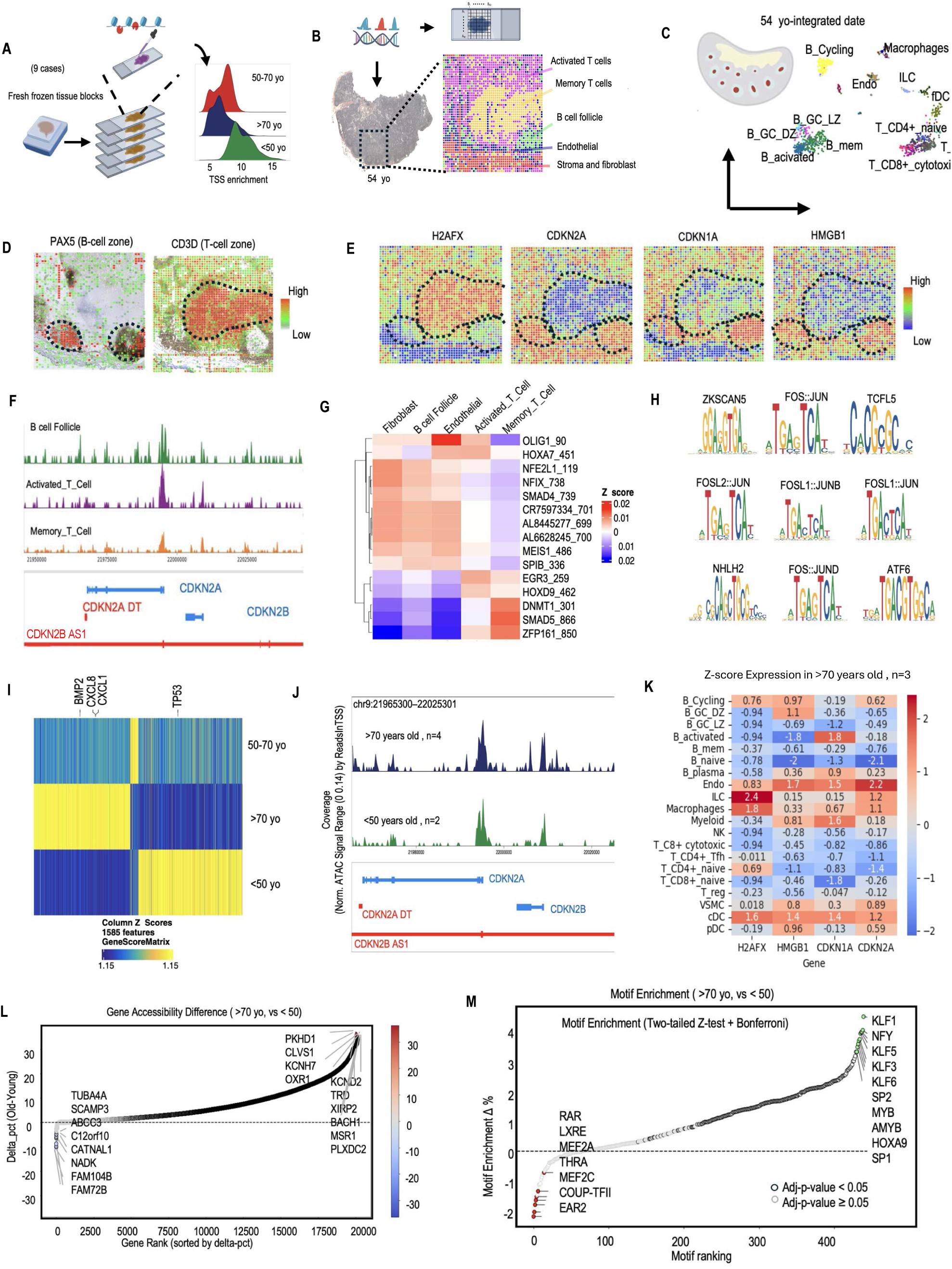
Spatial chromatin accessibility landscape in aging human lymph node tissues. (A) Spatial ATAC-seq profiling of chromatin accessibility in nine human lymph node samples spanning three age groups (<50, 50–70, and >70 years) using fresh-frozen tissue sections. Right, representative transcription start site (TSS) enrichment profiles demonstrating data quality. (B) Overview of the spatial chromatin accessibility analysis workflow. A representative lymph node section is shown with cell-type annotation and high-resolution spatial mapping. (C) UMAP clustering following integration of spatial ATAC-seq data with single-cell RNA-seq reference datasets, revealing major immune and stromal cell populations based on chromatin accessibility profiles. (D) Spatial distribution of *PAX5* accessibility (B-cell zones) overlaid with IHC staining for CD20 (highlighting germinal centers), and *CD3D* accessibility (T-cell zones) overlaid with IHC staining for BCL2 (highlighting the mantle zone), illustrating spatial correspondence between chromatin accessibility and histological features. (E) Spatial maps of chromatin accessibility for *H2AFX, CDKN2A, CDKN1A*, and *HMGB1*, highlighting region-specific variation across the tissue. (F) Genome browser tracks showing chromatin accessibility peaks in B-cell and T-cell regions at the *CDKN2A* locus. (G) Heatmap showing scaled transcription factor motif enrichment (Z-scores) across major cell clusters. (H) Top nine enriched transcription factor motifs identified in cells with high scores for the four senescence-associated markers, highlighting regulatory features of differentially accessible chromatin regions. (I) Heatmap displaying age-associated differences in gene activity scores across the three donor age groups. (J) ATAC-seq signal tracks at the chr9:21965300–22025301 locus, showing chromatin accessibility and annotated gene models for *CDKN2A, CDKN2B*, and their antisense transcripts in samples from >70- and <50-year-old donors. (K) Heatmap showing Z-score–scaled gene activity scores of key senescence-associated genes (*H2AFX, HMGB1, CDKN1A*, and *CDKN2A*) across annotated cell types. (L) Genome-wide comparison of gene accessibility between >70- and <50-year-old samples. Rank plot showing the difference in the proportion of cells with accessibility at each gene (□pct) between the two age groups. Positive values indicate a higher proportion of cells with accessible chromatin at that gene in >70-year-old samples. P values were calculated using a two-tailed z-test with Bonferroni–Dunn correction. Genes with adjusted p-value < 0.05 are outlined in black; non-significant genes are shown in grey. (M) Comparison of HOMER-defined motif enrichment between ATAC-seq peaks from >70- and <50-year-old donor groups. Negative values indicate motifs enriched in younger samples (red dots), whereas positive values indicate motifs enriched in >70-year-old samples (green dots). Motifs are ranked by enrichment in older peaks. P values were calculated using a two-tailed z-test with Bonferroni–Dunn correction. Motifs with adjusted p-value < 0.05 are outlined in black; non-significant motifs are shown in grey.

Several of these motifs correspond to the AP-1 transcription factor complex (e.g., *FOS:JUN*, *FOSL2:JUN, FOS:JUND*), which plays key roles in stress responses, inflammation, and senescence pathways. These findings further support the involvement of these gene regulatory networks in immune modulation and cellular senescence (Figure 6H).^30,74^

Analysis of age-associated senescence markers across all nine samples revealed a progressive activation of senescence-related programs. A heatmap shows increased accessibility of key senescence and inflammatory genes (*BMP2, CXCL8, CXCL1*) in samples from donors aged 50–70 and >70 years compared with those <50 years (Figure 6I). In contrast, *TP53* exhibited the highest chromatin accessibility in <50-year-old samples and declined with age, likely suggesting nuanced regulation of p53 activity through other mechanisms such as isoform usage and post-translational modifications that influence cell fate decisions between transient cell-cycle arrest, senescence, and apoptosis^75^. Looking across samples, donors younger than 50 years displayed greater chromatin accessibility at regulators of cell cycle progression and DNA repair, including *E2F1, BRCA2*, and *TP53* (Figure S11B). In contrast, samples from donors aged 50–70 and >70 years showed pronounced opening at canonical senescence genes (*CDKN1A, CDKN2A*) as well as SASP-related cytokines and growth factors (*CXCL8, BMP2, EGF, HGF*), indicating a shift toward senescence-associated inflammation and tissue remodeling (Figure S11B). Chromatin accessibility profiling at the *CDKN2A/B, CDKN1A*, and *H2AFX* loci revealed markedly higher and broader ATAC peaks in older lymph node samples, consistent with increased accessibility with age, in contrast to the lower and narrower peaks observed in samples from donors younger than 50 years (Figure 6J, S11C). Correspondingly, strong positive correlations were observed between the gene activity scores of key senescence and DNA damage markers (*CDKN1A, H2AFX*) and genes involved in chromatin regulation and related cellular processes (Figure S11D). Furthermore, the co-regulation of *CDKN1A* with nearby genomic elements, such as *RAB44*, suggests the presence of a broader regulatory network influencing cell-cycle control and cellular signaling at this locus (Figure S11D)^21,76^. *CDKN2A* accessibility is correlated with *CDKN2B*, supporting their coordinated roles in cell*-*cycle arrest (Figure S11D). The co*-*localization of *H2AFX* with neighboring genes, including *HMBS, HINFP, VPS11,* and *DPAGT*, indicates possible regulatory interactions modulating the DNA damage response and chromatin state (Figure S11D). These neighboring genes are linked to key processes such as histone modification (*HINFP*), porphyrin biosynthesis (*HMBS*), vesicular trafficking (*VPS11*), and glycoprotein biosynthesis (*DPAGT1*), collectively highlighting the regulatory landscape that may govern H2AFX*-*mediated DNA repair and senescence^76,76,77^. Integration with single-cell reference datasets revealed the distribution of key senescence-associated genes across diverse cell types in older donors (71, 74, 78, and 86 years). B cell subsets, including cycling-like B cells and plasma B cells, displayed notably higher scores for several senescence-associated genes. Cycling-like B cells exhibited elevated H2AFX and HMGB1, while plasma B cells showed increased CDKN1A and CDKN2A expression (Figure 5K). In contrast, T cell populations generally exhibited low or negative z-scores for these markers, indicating relatively lower senescence-associated activity (Figure 5K).

Comparison of gene accessibility and motif enrichment between donor groups >70 years and <50 years revealed age-associated shifts at both the gene and regulatory motif levels. Several genes, including *PKHD1, CLVS1, KCNH7, OXR1, KCNK2*, and *MSR1*, showed increased chromatin accessibility in older samples (Figure 6L) and are implicated in processes relevant to aging. For example, *PKHD1* has been associated with oxidative stress responses, tissue remodeling, and inflammatory signaling^78,79,80,81,82^. In contrast, genes such as *TUBA4A* (tubulin), *SCAMP3* (membrane trafficking), *ABCG3* (transporter), and *FAM72B* were more accessible in tissues from donors younger than 50 years, reflecting gene programs associated with robust cytoskeletal dynamics, intracellular transport, and cellular renewal that tend to decline with age (Figure 6L). Pathway analysis of genes with increased chromatin accessibility in >70-year-old compared with <50-year-old samples revealed significant enrichment of senescence- and stress-response pathways, including DNA damage– and telomere stress–induced senescence, TP53-regulated DNA repair, cell-cycle checkpoints, oxidative stress responses, NF-kB signaling, and SASP-related transcriptional programs (Figure S11F). Motif enrichment analysis reveals significant gains in accessibility for motifs recognized by the KLF (Kruppel-like factor) family (including *KLF1, KLF3, KLF5, and KLF6*) as well as *NFY* in >70 years old samples. The KLF family of transcription factors is known to regulate cellular senescence and cell cycle regulation and stress signaling^83,84^.In addition, *SP1*, *SP2, and HOXA9* demonstrate the regulators for inflammation, exhaustion, and senescence pathways^85,86^. In contrast, motifs such as *MEF2A* and *EAR2* are more enriched in <50 years old sample tissues, corresponding to genes and pathways involved in cell growth, metabolism, and anti-inflammatory signaling, all of which are diminished in the aging microenvironment^87^. Regulator of metabolism and development; shown as *COUP-TFII* (NR2F2) , *THRA*, *RAR*, *LXRE* showed impaired lipid handling and regeneration in aged tissues^88,89^.

Together, these spatial epigenomic analyses reveal age-associated chromatin accessibility changes in human lymph nodes, including increased accessibility at loci linked to stress responses, inflammatory signaling, and cell-cycle regulation, alongside reduced accessibility at genes involved in genomic maintenance and cellular renewal. These observations provide a spatially resolved view of epigenomic remodeling during lymph node aging and highlight intriguing regulatory features that warrant further investigation into the epigenetic mechanisms underlying cellular senescence in lymphoid tissues.

## Discussion

This study uses spatial multi-omics to map cellular senescence in human lymph nodes across age, focusing on core markers (p16, p21, HMGB1, *𝛾*-H2AX) and the SenMayo gene set. By integrating transcriptomic, proteomic, and epigenomic data, we reveal age- and lineage-specific senescence patterns and microenvironmental shifts that underlie immune function decline with aging. Senescent-like cells exhibit substantial heterogeneity in senescence gene expression, SASP, and inflammatory signaling. Clustering based on the SenMayo gene signature revealed distinct subpopulations of senescent cells, including clusters enriched for SASP components and pro-inflammatory cytokines, suggesting that senescent cells contribute to a sustained and diverse inflammatory responses^25,90^. This heterogeneity suggests that different cell types adopt senescence uniquely, affecting immune responses and the microenvironment differently. In aged lymph nodes, subsets of B cells – particularly cycling and germinal center subsets – exhibit a terminal senescent state marked by strong SenMayo and anti-apoptotic enrichment. On the other hand, VSMCs adopt a stromal-inflammatory senescent profile, and T cells largely resist classical senescence but develop a stress-adapted, pro-inflammatory phenotype. All these cells together reshape the germinal center into a presumably dysfunctional inflammatory niche. These B cells as key players in the germinal center reaction show the most pronounced age-related transcriptional changes in single-cell transcriptome data and their rates of proliferation, somatic hypermutation, and differentiation may indicate they are vulnerable to replicative and genotoxic stress, the key drivers of cellular senescence^91,92^.

In studies of human immune aging, hallmark features of immunosenescence—such as elevated inflammatory and SASP-associated transcriptional programs—have been most consistently observed in circulating lymphocytes, including CD4*⁺* T cells, CD8*⁺* T cells, NK cells, and classical monocytes. These findings align with recent PBMC aging studies showing age-related loss of naïve CD8*⁺* and MAIT cells, a shift toward differentiated T-cell states, inflammatory skewing (even in naïve CD8*⁺* T cells), and monocyte remodeling. Aging-clock and transcriptomic-age models further link older immune states to increased monocyte inflammation and heightened senescence-associated programs (inflammatory, stress, and DNA-damage response pathways), supporting the conclusion that late-life PBMC aging couples inflammaging with canonical senescence pathways^93,94^. In contrast, lymph nodes exhibit a predominantly B-cell–centric senescence phenotype. Germinal center (GC) dark zone and light zone B cells, cycling-like B cells, naïve B cells, and plasma cells are strongly enriched for SenMayo signatures and cell-cycle–arrest programs across multiple modalities. This pattern suggests that the follicular adaptive niche acts as a major reservoir of senescent cells during aging, where the sustained demands of antibody production and high metabolic activity increase B-cell susceptibility to senescence programs. Spatial protein imaging revealed age-dependent patterns of senescence markers in lymph nodes: dispersed in young individuals, clustering in individuals aged 50–70, and predominantly accumulating in follicles of individuals over 70, particularly for p16 and p21. These spatial shifts reflect dynamic remodeling of senescence within the tissue architecture. The transition from interfollicular regions in youth to follicular accumulation in older adults marks a niche-specific remodeling in which B cells strongly activate stress-response and anti-apoptotic pathways, forming “SenSpots” within follicles that act as hubs of chronic inflammation and immune dysfunction. Complementary proteomic and metabolic analyses further indicate that aged germinal center niches undergo profound molecular reprogramming, including a shift toward lipid accumulation and a more oxidized cellular state. These cells also exhibit compromised telomere maintenance and mitochondrial dysfunction, reflecting active metabolic and structural rewiring rather than passive quiescence.

To explore the potential preliminary mechanism of immunosenescence in situ in human lymphoid tissues, spatial whole-transcriptome sequencing was performed on selected samples and regions identified through spatial protein mapping to investigate the mechanisms underlying spatial cellular senescence in human lymph nodes (LNs). We observed enrichment and differential expression of genes associated with key senescence markers—including genes corresponding to p16, p21, HMGB1, and *𝛾*-H2AX—across follicles, cortex, and interfollicular spaces at high resolution, highlighting both spatial and cellular heterogeneity. Senescent B-cell niches exhibited upregulation of stress- and immune-regulatory genes, such as *FOS/AP-1* and NF-kB–associated pathways, alongside downregulation of immunoglobulin transcripts, consistent with reduced antibody production. Epigenetic analyses revealed extensive age-associated remodeling in LNs, characterized by increased chromatin accessibility at SASP loci and inflammatory cytokine and growth factor genes. This state also emphasized stress- and senescence-regulatory factors, particularly AP-1 family members and KLF family transcription factors (Table S6). Consistent with transcriptomic patterns, B-cell subsets—including cycling-like B cells, plasma B cells, and GC B cells—exhibited the strongest senescence-associated accessibility signatures, whereas T cells showed comparatively limited changes, underscoring lineage-specific epigenetic trajectories during immune aging.

Our integrative multi-omics analysis—combining transcriptomics, proteomics, epigenomics, and metabolic imaging—provides a spatially resolved, cell type–defined, genome-scale view of cellular senescence in human lymph nodes across the lifespan. Although still preliminary, this work offers a quantitative atlas of senescent-like cell states, their single-cell and spatial heterogeneity, and the tissue niches they occupy, making the findings particularly intriguing. Senescence emerges as a complex, cell type- and niche-specific process, with cycling-like, memory, and germinal center B cells as the key cell types of interest in this study, accumulating in follicular “SenSpots” characterized by distinct metabolic, transcriptional, and chromatin signatures that reflect both terminal and transitional senescent states (Table S6). These observations redefine the lymph node as a dynamic immune organ during aging, shaped by spatial and molecular senescence patterns, and highlight the importance of understanding how senescent cells and senescence-associated programs contribute to immune aging. This preliminary atlas provides a framework for dissecting cell type–specific senescence states, their tissue niches, and their potential impact on immune function across the lifespan.

### Limitations of the study

This study provides a comprehensive spatial multi-omics analysis of cellular senescence in human lymph nodes across the lifespan, yet several limitations remain. While our cohort spanned a wide age range, the single-cell RNA-seq reference dataset for lymph nodes lacked samples from individuals aged 41–59, limiting our ability to capture gradual senescence-associated changes during middle age. Additionally, certain cell types were unevenly represented across age groups due to age-related shifts in lymph node composition, which may affect analyses of senescence marker expression and pathway activity. Sample availability also posed challenges: lymph nodes are rarely banked without clinical suspicion of malignancy, restricting access to tissue from fully healthy individuals. For instance, biopsies from younger patients, although histologically benign, were often collected in the context of unrelated clinical conditions that could elevate inflammatory and senescence-associated markers such as SASP and DNA damage response pathways, as reflected in our analyses. Despite these potential confounders, spatial senescence scores displayed consistent age-associated trends, whereas scores for inflammation, proliferation, and exhaustion were more variable, suggesting that cellular senescence represents a core aging mechanism relatively independent of other transient responses. To integrate spatial transcriptomic and epigenomic data with scRNA-seq, we employed Harmony, enabling cross-modality alignment and interpretation^130^. However, Harmony was originally developed for integrating scRNA-seq datasets and may not fully capture the complexities inherent to spatial transcriptomic technologies. Although integration of spatial multimaps with CODEX and SRS achieves near-cellular resolution, differences in sensitivity across modalities and samples can challenge data consistency, complicate co-registration, and potentially affect the accuracy of cell type annotation and senescent-like cell identification. Some CODEX lymph node samples also contained substantially more cells than the scRNA-seq data, which could introduce integration biases. Multi-omics analyses further depend on careful serial sectioning of tissue blocks, yet anatomical variation between sections complicates alignment even when using advanced tools such as STalign. In this study, we identify novel metabolic and epigenomic signatures of cellular senescence associated with p16 and p21 in B and T cells, as well as other immune cell types of interest. Nevertheless, further functional experiments are required to validate these findings. For instance, studies that selectively deplete, inhibit, or induce senescence in B and T cells would provide insight into not only changes in canonical senescence markers but also functional consequences under specific perturbations, clarifying their roles in the broader contexts of immunosenescence and inflammaging.

## ACKNOWLEDGMENTS

We thank the Yale West Campus cleanroom team for assistance with microfluidic wafer fabrications and the YPTS team for FFPE tissue sectioning and staining. Computational data analysis was conducted with Yale High Performance Computing clusters (HPC). We would like to thank the Yale Pathology team for their help in tissue handling. We would like to acknowledge the support received from the U.S. National Institutes of Health including grants U54CA274509, UH3CA257393, RF1MH128876, U54AG079759, U54AG076043, R01CA245313, RM1MH132648 (all to R.F.), and U01CA294514 (to R.F., M.X., and Z.M.). Z.M. is supported by NSF awards 2345215 and 2245575.

## Author Contributions

Conceptualization: R.F., Y.K. Methodology: N.F, A.E, Y.L, F.P, A.F, and Y.K Experimental Investigation: N.F., A.E.,Y.K, Y.L,L.C,M.Z, Z.B. Data Analysis: N.F, A.E, Y.L, A.F., F.P, M.L, M.Y., J,Y., and J.C. Data Interpretation: N.F., A.E., F.P., R.F., M.X. and Y.K. Resources and valuable inputs: N.F, A.E, Y.L,A.F,F.P., S.F,M.X and J.C. Data deposition: F.S, M.Y, Y.K. Original Draft: N.F., A.E., and R.F. All authors reviewed, edited, and approved the manuscript.

## Materials and Methods

### Tissue specimen bank (Lymph node resection and biopsy)

De-identified archival formalin-fixed paraffin-embedded (FFPE) human reactive lymph node tissue blocks, originally collected by physicians for diagnostic purposes, were obtained from the Yale Pathology Tissue Services (YPTS). The collection, retrieval, and distribution of tissue samples for research purposes were carried out under the approval of the Yale University Institutional Review Board, with oversight from the Tissue Resource Oversight Committee. Written informed consent for participation, including instances where identifying information was collected alongside specimens, was obtained from patients or their legal guardians in accordance with the Declaration of Helsinki. All samples were handled in strict compliance with HIPAA regulations, University Research Policies, Pathology Department diagnostic protocols, and Hospital by-laws. A total of 33 patient samples were collected. The biopsies were collected across a wide range of age groups (1 to 86 years old) from different sites in the body including the axillary, inguinal, groin, submental and neck. All diagnoses were reviewed for accuracy centrally by a board-certified pathologist. Each sample was confirmed as a non-malignant lymph node without significant anatomic pathology. This tissue bank includes frozen tissue blocks in OCT and Formalin-Fixed Paraffin Embedded (FFPE). All tissue preparation steps from harvesting to embedding in paraffin were done in RNase-, DNase-, and protease-free conditions (Supplementary Table 1).

### Sample handling and section preparation

Paraffin and fresh frozen blocks were sectioned at a thickness of 7-10 µM and mounted on the center of Poly-L-Lysine coated 1 x 3" glass slides in a cryo-chamber (-20 °C). Before sectioning, the frozen tissue block was warmed to the temperature of cryotome cryostat (-20°C). Serial tissue sections were collected simultaneously for DBiT-seq, Histological staining (H&E), immunohistochemistry (IHC), CODEX and spatial-ATAC-seq. Some sections in between were mounted on VWR Superfrost Plus Micro Slides (48311703) for NanoString CosMx. The sectioning of biopsy samples was carried out at YPTS. Paraffin and frozen sections were shipped in tightly closed slide boxes or slide mailers in dry ice and stored at - 80°C upon receipt until use. All tissue slides were directly kept at - 80°C if long-time storage is needed.

**Histopathological observation**

All original H&E slides and immunohistochemistry (IHC) studies of cases were reviewed by a board-certified pathologist with hematopathology subspecialty certification. One representative block per case was selected and retrieved for further analysis. FFPE and FF sectioning were performed at YPTS. These procedures adhered to Clinical Laboratory Improvement Amendments (CLIA)-certified laboratory protocols, where clinical-level IHCs were obtained. Sections were stained for CD20 (clone L26, no dilution; Dako). Formalin-fixed, paraffin-embedded tumor tissue sections were deparaffinized and rehydrated before antigen retrieval and primary antibody incubation. Subsequently, specimens were incubated with diaminobenzidine chromogen for primary antibody detection and counterstained with hematoxylin^95^.

### Proteomic analysis (CODEX spatial phenotyping using PhenoCycler-Fusion)

Spatial high-plex phenotyping of the adjacent FF and FFPE section were performed following the CODEX PhenoCycler-Fusion user guide^96,97^.A 53-plex panel was utilized for the CODEX imaging consistent with 44 conjugated antibodies purchased from AKOYA and in house conjugated antibodies (p21, p16, HMGB1, CXCL13, and CXCR5) were incorporated into this panel (Table S2). Antibodies were conjugated with a DNA based reporter using the AKOYA conjugation kit (SKU 7000009). The validation and quantification of the conjugated antibodies was performed by electrophoresis gel^96,97^ . The tissue sectioned on Poly-L-lysine (PLL) slides were obtained with the thickness of 5-7 um from both FF and FFPE blocks. The tissue preparation including deparaffinization, hydration, antigen retrieval and equilibration in staining buffer followed by antibody cocktail staining and post fixation following Akoya protocol^96,97^.The tissue on slide was attached to the flow cell and CODEX cycles were configured on the system following loading the prepared reporter, and the imaging process commenced. Upon completion of the imaging cycles, a final QPTIFF file was produced and visualized using QuPath V0.5.0102. Details regarding PhenoCycler antibody panels, experimental cycle design, and reporter plate volumes can be found in Supplementary Table 2.

Standard preprocessing of scRNA-seq data was performed using Scanpy, followed by integration with CODEX data based on linked features with high variability. Pivot cells, representing approximately 10% of the CODEX dataset, were identified through high confidence matches, using principal components and a smoothing weight during pivot matching. Cell type labels from scRNA-seq were transferred to these pivot cells, and a Support Vector Machine (SVM) model was trained on them to annotate the remaining non-pivot CODEX cells based on protein expression. The integration allows for the mapping of scRNA-seq-derived cell types onto spatially resolved CODEX images, providing a comprehensive spatial transcriptomic and proteomic landscape of the lymphoid tissue.

### Validation of Senescence markers and conjugated antibodies

A p16+ nasopharyngeal carcinoma tissue confirmed our conjugated senescence markers’ staining. A positive staining of the p16+ tumors within the tissue section was first confirmed by IHC. The conjugated P16 and P21 antibodies were confirmed on the serial section of this tissue using CODEX. We also observed subset of the tumor cells which stain positive for P16 and P2 (Figure S3).

### Spatial multi-omics (DBiT)

The DBiT-seq device was fabricated polydimethylsiloxane (PDMS) with 20µM channel width (Front Range Photomasks (Lake Havasu City, AZ) was prepared. A protocol to perform SU-8 photolithography, development, and hard baking for photomask was followed based on the manufacturer’s (MicroChem) and previous published protocol^12,98^ . DNA oligos for spatial barcodes A and B in this study were purchased from Integrated DNA Technologies (IDT, Coralville, IA) and the sequences were listed in Supplementary Table 3. Barcode (100 µM) and ligation linker (100 µM) were annealed at a 1:1 ratio in 2X annealing buffer and stored at 20°C until use^12,98^.

In spatial transcriptomic experiments, the FFPE tissue sections on a PLL slide were retrieved from the 80°C freezer and deparaffinized (following Patho-DBiT protocol, Supplementary Table 3)^16^. The tissue slide was then submerged in a boiling 1X antigen retrieval buffer for 20 minutes, followed by a 10-minute cool down to room temperature. After a quick rinse in distilled water, a scan from the intact tissue was captured using a 10X objective on the EVOS M7000 Imaging. The tissue was then permeabilized using 1% Triton X-100 in DPBS at room temperature for 10 minutes. In situ polyadenylation was performed on tissue by adding 60 µL of the Poly(A) enzymatic mix (38.4 µL nuclease-free water, 6 µL 10X Poly(A) Reaction Buffer, 6 µL 5U/µL Poly(A) Polymerase,6 µL 10mM ATP, 2.4 µL 20 U/µL SUPERase•In RNase Inhibitor, 1.2 µL 40 U/µL RNase Inhibitor) and incubated in a humidified box at 37C for 30 min following Patho-DBiT protocol. After removing the excessive reagents with DPBS for 5 min, 60 µL of the reverse transcription mix (20 µL 25 µM RT Primer, 16.3 µL 0.5X DPBS-RI, 12 µL 5X RT Buffer, 6 µL 200U/µL Maxima H Minus Reverse Transcriptase, 4.5 µL 10mM dNTPs, 0.8 µL 20 U/µL SUPERase•In RNase Inhibitor,0.4 µL 40 U/µL RNase Inhibitor) was loaded into the PDMS reservoir and sealed with parafilm. The sample was incubated at room temperature for 30 min and then at 42 °C for 90 min, followed by DPBS wash as described before^16^.

In spatial ATAC-seq experiments, the FF tissue sections on a PLL slide were fixed with formaldehyde (0.2% for 5 min) and permeabilized with 500 µl lysis buffer (10mM Tris-HCl, pH 7.4, 10mM NaCl, 3mM MgCl2, 0.01% Tween-20, 0.01% NP-40, 0.001% digitonin, 1% BSA) for 15 min. in situ transposition was applied by adding loaded Tn5 transposition mix (50 µl 2X tagmentation buffer, 33µl 1XDPBS, 1µl 10% Tween-20, 1µl 1% digitonin, 5µl transposase, 10 µl nuclease-free H2O ) on the tissue^98^. Finally, tissue was washed with EDTA, 500 µl 1xNEBuffer 3.1 and dip in water sequentially.

In the barcoding by DBiT-seq device, each fresh frozen and FFPE tissue sections were firstly attached to the first direction of PDMS to perform in situ barcodes A ligation. The chip includes tissue slide and first direction PDMS was imaged using a desired optical microscopy to record the positions of the 50 channels on the ROI for downstream alignment and analysis. After securing the PDMS on the PLL slide with an acrylic clamp, the annealed DNA barcodes A with the ligation linker 1 in a ligation mixture (The ligation mix, comprising 100 µL 1X NEBuffer 3.1, 61.3 µL nuclease-free water, 26 µL 10X T4 ligase buffer, 15 µL T4 DNA ligase, 5 µL 5% Triton X-100, 2 µL 40 U/µL RNase Inhibitor, and 0.7 µL 20 U/µL SUPERase RNase Inhibitor) at the ratio of 1:4 were added into each inlet at 5 ul volume. An adjusted vacuum was applied on the outlet of the chip to flow the reagents on the ROI through the channel. After a 30-minute incubation at 37 °C, the PDMS chip was removed, and the slide was washed with 50 mL DPBS. Afterwards, the PDMS for the second direction, containing 50 perpendicular channels to the first direction PDMS, was positioned on the slide and the region of interest (ROI). A bright-field image was then captured, and ligation of the 50 barcodes B were carried out following the first ligation. The washed and air-dried tissue slide with a defined ROI was scanned using an optical microscope to record a bright-field image for downstream analysis.

### Cell type annotation in DBiT-seq

We integrated DBiT-seq data with high-resolution H&E image using iStar and predict super resolution gene expression on the whole tissue section. To annotate our DBiT-seq data, Harmony and KNN are harnessed to integrate with our scRNA-seq reference dataset. Our DBiT- seq data is also co-registered with CODEX data. We then identify pixels enriched of B cells and senescence related markers. Differential gene expression and pathway analysis are then carried out between senescent and non-senescent B cells.

### CosMx(Single-cell level spatial transcriptomics on whole tissue)

One FFPE sample from a 86 years old donor was chosen from the TMC archive and sectioned on VWR Superfrost Plus Micro Slides (48311703), following the CosMx tissue handling protocol^99^. The 6000 human CosMx panel was applied on both slides using the CosMx SMI instrument at Yale Center for Genome Analysis (YCGA).

### Senescence scoring of CosMx dataset using the SenMayo gene set

To mitigate technical sparsity, cells were analyzed using the previously described ‘SenMayo’ gene set, which has been validated to identify cellular senescence. The expression of transcript 𝑖 in cell 𝑗, 𝑇_i_(𝑗), as normalized by dividing by the highest observed expression across all 𝑛 cells in the sample:

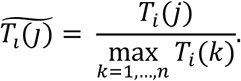

The overall expression score (OES) for the SenMayo gene set, 𝐺, in cell 𝑗, 𝑆(𝑗; 𝐺), was calculated by summing the normalized expressions of all genes contained in the gene set:

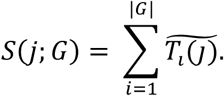

These scores were then standardized using z-normalization,

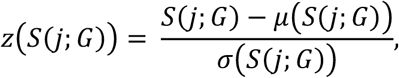

where 𝜇 and 𝜎 are the mean and standard deviation of the OES across all 𝑛 cells. For visualization, cells in tissue samples were visualized by plotting points corresponding to their spatial center; z-scored OES’s were mapped to a color gradient.

Similarly, we aggregate gene expression over Fields of View (FOV) to identify tissue regions with relatively higher or lower levels of p16 and p21 expression. Each FOV consists of a small, contiguous tissue section, which is independently sequenced using the CosMx protocol. The average expression of a gene, 𝑔, within an FOV, 𝐹, is computed by taking the mean expression across the 𝑛 cells in 𝐹,

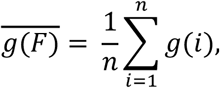

where 𝑔(𝑖) represents the expression of gene 𝑔 in the 𝑖-th cell. We standardize across all FOVs in a single tissue by taking the z-score in the standard way,

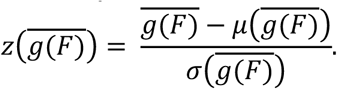

Here, 𝜇 and 𝜎 are the mean and standard deviation of the FOV mean gene expression estimator 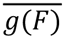. For visualization, as above, FOVs were visualized by plotting points corresponding to their spatial and the z-scored mean gene expression was mapped to a color gradient.

Raw counts were used for analysis. Scripts were implemented in Python 3.10. Numpy (v 1.26.4) and Pandas (2.2.2) were used for data processing; plots were generated using Matplotlib (v 3.9.1).

### Laser capture microdissection proteomics

Laser ablation of tissue sections. The robotic arm was installed on a PALM MicroBeam LCM system (Carl Zeiss MicroImaging, Munich, Germany) to precisely position the capillary tip at the center point between the laser source and microscope lens, ensuring optimal alignment for accurate sample collection. To maximize collection efficiency, the distances between dimethyl sulfoxide (DMSO) droplet and tissue slide or nanoPOTS chips for sample collection, deposition, and capillary cleaning were measured to properlySingle-cells were selected and isolated at 63□ magnification using PalmRobo software. The outlines of single-cell were guided by p16-IHC staining and cell morphology. The ‘AutoLPC’ function was used to ablate tissue sections and directly catapult the ablated tissue particles into the DMSO droplet within the nanoPOTS well. Collection was performed automatically using custom software. The collected samples were dried at 70 °C, nanoPOTS chips were covered with microscope slides, wrapped in aluminum foil, and stored at -20 °C until use.

Proteomic sample processing. A home-built nanoliter robotic liquid-handling platform was employed to dispense reagents into nanowell. In brief, 200 nL of cell lysis buffer (1 mM tris(2-carboxyethyl)phosphine, 0.1% n-dodecyl-ß D-maltoside, and 50 mM ammonium bicarbonate (ABC) buffer, pH 8.0) was added to each nanowell and incubated at 70 °C for 1 hour to facilitate cell lysis, protein extraction, and denaturation. Following this, 50 nL of alkylation buffer (10 mM 2-chloroacetamide in 50 mM ABC buffer) was introduced and left to react in the dark at room temperature for 30 minutes. Subsequently, 50 nL of digestion buffer (0.01 ng/nL Lys-C and 0.04 ng/nL trypsin in 50 mM ABC buffer) was added and samples were incubated at 37 °C for 10 hours to complete enzymatic digestion. Finally, to terminate the enzymatic reaction, 50 nL of 5% formic acid (FA) was added and incubated at room temperature for 30 minutes. After processing, the samples were dried in a desiccator and stored at -20 °C for subsequent LC-MS analysis.

LC–MS analysis. Samples were analyzed using a home-built nanoPOTS LC platform coupled to an Orbitrap Fusion Lumos Tribrid MS (Thermo Fisher Scientific) with an eld asymmetric ion mobility spectrometry (FAIMS) interface. A solid phase extraction (SPE) column (100 µm i.d., 4 cm long) packed with 5 µm, C18 packing material (300 Å pore size; Phenomenex) and an analytical column (50 µm i.d., 25 cm long) packed with 1.7 µm, C18 particles (BEH 130 Å pore size; Waters) were employed for on-line sample cleanup and separation of peptides, respectively. The analytical column was heated to 50 °C using a column heater (Analytical Sales and Services Inc., Flanders, NJ, USA). The mobile phases were composed of buffer A (0.1% FA in water) and buffer B (0.1% FA in acetonitrile) and delivered using an UltiMate 3000 RSLCnano System (Thermo Fisher Scientific). The dried samples were directly injected from nanoPOTS chips and transferred to the SPE column for a 5-minute cleanup using 5% acetonitrile in 0.1% formic acid at a flow rate of 3 µL/min. Subsequently, peptides were eluted and transferred to the analytical column for chromatographic separation at a flow rate of 100 nL/min. The separation utilized a gradient elution: initially, a 30-minute linear increase from 8% to 22% buffer B, followed by a 9-minute linear increase from 22% to 35% buffer B. To enhance sensitivity and accuracy of peptide/protein identification for low-input samples, we applied previously developed Transferring Identification Based on FAIMS Filtering (TIFF) method, which involves the use of a spectral library for identifying peptides detected in low-input samples. For the construction of the spectral library, we digested bulk samples derived from a whole slide section taken from the same lymph node block. The spectral libraries were generated by analyzing 50 ng of these samples using discrete FAIMS compensation voltages (CVs) of -45, -60, and - 75 V for each LC-MS analysis. For the analysis of single-cell samples, we used these same three FAIMS CVs (-45, -60, and -75 V) in a single LC-MS analysis. For the single-cell samples, the mass spectrometer was operated in positive ion mode with an electrospray voltage of 2.4 kV and an ion transfer tube temperature of 250 °C. Data acquisition was performed in data-dependent acquisition (DDA) mode with MS1 scan range set to 350-1600 m/z at a resolution of 120,000, automatic gain control (AGC) target of 1E6 ions, and a maximum injection time of 500 ms. For MS2 acquisition, precursor ions with charges ranging from +2 to +6 and intensities greater than 1E4 were isolated using a 1.4 m/z window and fragmented by 30% high-energy dissociation (HCD). Fragment ions were then detected in the ion trap with an AGC target of 2E4 and a maximum injection time of 150 ms. The cycle time was set to 0.8 seconds. For library samples, the maximum injection time was 118 ms for MS1 and 86 for MS2, with a cycle time of 2 seconds, while keeping all the other parameters the same.

Data analysis. LC-MS raw files were analyzed using FragPipe (Ver. 21.1) powered by MSFragger (Ver. 4.0) search engine, Philosopher (Ver. 5.1.0) and IonQuant (Ver. 1.10.12 ) for LC-MS feature detection, database searching, and protein/peptide quantification using TIFF method as previously described. Peptides from low-input samples were identified by matching to the reference library based on 3D features, including LC retention time, accurate m/z, and FAIMS CVs. The UniProt human (Homo sapiens) database (2023 release, UP000005640_9606) was searched with a peptide spectral match and protein-level false discovery rate (FDR) of 1%. Strict trypsin was set as the cleavage enzyme. The precursor and fragment mass tolerances were set to 20 ppm. Methionine oxidation (+15.9949) and protein N-terminal acetylation (+42.0106 Da) were set as variable modifications. Carbamidomethylation (+57.02146 Da) on cysteine was set as fixed modification.

### Stimulated Raman Spectroscopy (SRS)

The SRS were collected from an upright laser-scanning microscope (DIY multiphoton, Olympus), which was equipped with a 25x water objective (XLPLN, WMP2, 1.05 NA, Olympus). The synchronized pulsed pump beam (tunable 740-990 nm wavelength, 5–6 ps pulse width, and 80 MHz repetition rate) and stokes beam (wavelength at 1031nm, 6 ps pulse width, and 80 MHz repetition rate) from a pico Emerald system (Applied Physics & Electronics) were coupled and introduced into the microscope. Upon interacting with the samples, the transmitted signal from the pump and Stokes beams was collected by a high NA oil condenser (1.4 NA). A shortpass filter (950 nm, Thorlabs) was used to completely block the Stokes beam and transmit the pump beam only onto a Si photodiode array for detecting the stimulated Raman loss signal. The output current from the photodiode array was terminated, filtered, and demodulated by a lock-in amplifier at 20 MHz The demodulated signal was fed into the FV3000 software module FV-OSR (Olympus) to form images using laser scanning. SRS protein, lipid, unsaturated lipid and saturated lipid signal were obtained at vibrational modes 2930 cm-1, 2850 cm^-1^, 2880 cm^-1^ and 3015 cm^-1^ in a single tile of 512 x 512 pixels, at dwell time 40 µs and 100 tiles were stitched to generate the large tissue mapping. To quantify the unsaturated lipid ratio, the intensity of C=C vibrational modes (peak intensity at 3015 cm^-1^) was divided by the intensity of saturated vibrational modes (peak intensity at 2880 cm^-1^), resulting in ratiometric images. Minor intensity adjustment (brightness and/or contrast) due to vignetting in tile boundaries was performed using VISTAmap^100^.

### Optical redox imaging

The two-photon fluorescent microscopy was used to collect the NAD(P)H and FAD signals from the tissue. The NAD(P)H signal was excited by 780 nm, the emission wavelength was collected at 460 nm. The FAD signal was excited by 860 nm, the emission wavelength was collected at 515 nm. The ratiometric images were generated by FAD/NAD(P)H. The ratio was further quantified in ImageJ and plotted in GraphPad 10.

### Single cell RNA-seq reference data

To build a diversified single-cell RNA sequencing (scRNA-seq) reference dataset for lymph node analysis, two scRNA-seq datasets are integrated: Tabula Sapiens (TS)^17^ and an integrated secondary lymphoid organ (SLO) atlas^18^.The integration process begins with filtering cells that have fewer than 30 detected genes and genes present in fewer than 20°C ells. This is followed by total count normalization, scaling, and the selection of 4,000 highly variable genes. Harmony is then used to integrate the datasets and eliminate technical variations^101^. Since the SLO dataset offers more granular annotations, a KNN classifier is employed to propagate SLO annotations to the TS dataset, excluding cells with annotation probabilities below 0.8 from the final dataset. The resulting lymohoid scRNA-seq reference dataset comprises 95,389 cells, profiling 10,166 genes from lymph nodes (LN), spleen, and tonsil.

### Single cell RNA-seq data analysis

Publicly available scRNA-seq lymph node (LN) data from HuBMAP is collected and integrated with the reference scRNA-seq dataset using Harmony and KNN to establish the LN scRNA-seq dataset. This dataset comprises 108,518 cells across 8,156 genes from 14 donors, ranging in age from 10 to 79 years. The donors are divided into two groups (10-50 and 51-79 years old) for comparison of gene enrichment scores related to the SenMayo gene set and canonical senescence gene markers. Age-associated B cells (ABCs) are defined as B cells expressing the T-bet gene. The proportion of ABCs and the expression levels of canonical senescence gene markers are explored between ABCs and non-ABCs across the two age groups. To better understand the heterogeneity of senescence-like cells, the top 10% of cells with the highest SenMayo gene enrichment scores are selected as senescence-like cells. The overlapping genes between the SenMayo gene set and the LN scRNA-seq data are then used to further classify these senescence-like cells into subgroups. Differential gene expression (DGE) and pathway analysis are performed to investigate the genes and pathways in these senescence-like cell subgroups. For cell type annotation, we integrated the scRNA-seq dataset of lymph nodes from Tabula Sapiens (TS) with a well-annotated secondary lymphoid organ (SLO) atlas. Integration was performed using Harmony, followed by K-nearest neighbor (KNN) classification to assign TS cells to the closest SLO-derived cell types. Using this approach, we successfully annotated 34 immune and stromal cell types. For overall visualization, we applied a grouping strategy that merged closely related immune and stromal subtypes to improve clarity while preserving key distinctions among major cell populations. Specifically, B_GC_DZ and B_GC_LZ were combined into a single “B_GC” cluster, and cycling-like B, B_activated, and B_plasma were grouped as “B_other.” Dendritic cell populations (FDC, DC_CCR7⁺, DC_cDC1, DC_cDC2, and DC_pDC) were merged into a single “DC” group, while macrophage subsets (Macrophages_M1 and Macrophages_M2) were combined as “Macrophages.” T_CD4⁺_TfH, T_CD4⁺_TfH_GC, and T_CD4⁺_naive were grouped into “T_CD4⁺,” and T_CD8⁺_CD161⁺, T_CD8⁺_cytotoxic, and T_CD8⁺_naive were grouped as “T_CD8⁺.” Regulatory T cell subsets (T_TfR and T_Treg) were combined into a “T_other” cluster. The “Other” cluster included ILC, VSMC, mast cells, monocytes, and NKT cells. Cell types such as B_memory, B_naive, NK cells, and endothelial cells were retained as individual categories. In some analyses (SRS, spatial transcriptomic, and spatial epigenomic), we applied an alternative grouping to more specifically assess B and T cell heterogeneity. B_naive included both B_naive and B_preGC cells, while B_GC_LZ combined B_GC_LZ and B_GC_prePB. Cycling-like B , B_GC_DZ, B_plasma, and B_memory were kept as separate categories, whereas B_IFN and B_activated were merged as “B_activated.” For T cells, T_CD8⁺_cytotoxic, T_CD8⁺_CD161⁺, and NKT were combined as “T_CD8⁺ cytotoxic.” T_CD4⁺_TfH and T_CD4⁺_TfH_GC were merged as “T_CD4⁺_TfH,” while T_CD4⁺_naive and T_CD4⁺ were combined as “T_CD4⁺_naive.” Regulatory T cells (T_Treg, T_TIM3⁺, and T_TfR) were grouped as “T_reg,” and T_CD8⁺_naive was maintained as an individual category. For antigen-presenting cells, DC_cDC1, DC_cDC2, and FDC were grouped as “cDC,” while DC_pDC and DC_CCR7⁺ were combined as “pDC.” Macrophages_M1 and Macrophages_M2 were combined as “Macrophages,” whereas NK, ILC, endothelial cells (Endo), and VSMCs were kept as individual categories. Monocytes and mast cells were grouped as “Myeloid.”

### Proteomic analysis and integration

MaxFuse is utilized to integrate scRNA-seq reference data with CODEX data. To mitigate sparsity in scRNA-seq data, meta-cells of size 2, aggregated from cells with similar features, are constructed. Initially, all features from both scRNA-seq and CODEX data are used to construct a nearest-neighbor graph for each modality separately. MaxFuse then smooths the shared features between scRNA-seq and CODEX data to generate the initial cross-modal cell matching. For this initial matching, a weight of 0.3 is assigned to both modalities. The initial matching is iteratively refined using all features from both scRNA-seq and CODEX data, with one iteration chosen due to the low signal-to-noise ratio in the CODEX data. Following this refinement, cell pairs are filtered with a threshold of 0.5 to obtain refined pivots, resulting in approximately 60% of CODEX cells finding matches in the scRNA-seq data. The refined pivots are then used to propagate the matching to the remaining cells, with a cut-off of 0.3 applied to filter out poor matches. In the final result, more than 98% of CODEX cells are matched with those in the scRNA-seq data. Protein expression is calculated by summing the marker intensity within each cell and normalizing the total intensity by cell size. During preprocessing, total-protein normalization and log1p scaling are applied to generate the final protein expression matrix. To avoid batch effects between CODEX samples, senescence markers are analyzed separately for each sample, with specific senescence marker-positive cells defined as those with the highest 10% senescence marker expression. The proportions of these senescence marker-positive cells are then calculated and compared across all cell types, regions of interest (ROIs), and samples.

### Sequence alignment and generation of gene expression matrix

Read 2 from the FASTQ file was used to extract unique molecular identifiers (UMIs) and spatial barcodes A and B. Read 1, containing cDNA sequences, was aligned to either the mouse GRCm39 or human GRCh38 reference genome using STAR version 2.7.8a. Spatial barcodes were demultiplexed with ST_Pipeline version 1.8.1, based on the predefined coordinates of the microfluidic channels, and ENSEMBL IDs were converted to gene names. This process generated a gene-by-spot expression matrix for further analysis, with entries corresponding to spot positions lacking tissue being excluded.

### Gene data normalization and unsupervised clustering analysis

Spatial gene expression analysis was performed using the Seurat (4.2). Gene expression for each spot was first normalized and variance-stabilized using the SCTransform method, which is optimized for single-cell RNA sequencing (scRNA-seq) data. Linear dimensional reduction was applied with the "RunPCA" function, and the optimal number of principal components for further analysis was determined heuristically, visualized through an ’Elbow plot’ that ranked PCA components by their variance percentages. The "FindNeighbors" function then embedded spots into a K-nearest neighbor graph using Euclidean distances in PCA space, while the "FindClusters" function employed a modularity optimization technique to group the spots into clusters. The "RunUMAP" function was utilized to explore spatial heterogeneity via the Uniform Manifold Approximation and Projection (UMAP) algorithm. Lastly, differentially expressed genes (DEGs) characterizing each cluster were identified through the "FindMarkers" function, which conducted pairwise comparisons between groups of spots.

### High-resolution tissue architecture inference using iStar

iStar is a computational method that utilizes hierarchical image feature extraction to integrate spatial transcriptomics (ST) data with high-resolution histology images, enabling super-resolution predictions of spatial gene expression. The iStar algorithm consists of three key components: a histology feature extractor, a high-resolution gene expression predictor, and a tissue architecture annotator, as described in our previous work. In brief, H&E images were rescaled to standardize pixel dimensions, divided into hierarchical image tiles, and histology features were extracted and trained using a vision transformer. A feed-forward neural network, trained through weakly supervised learning, predicted super pixel-level gene expression based on the top 1,000 most variable genes from the Patho-DBiT expression matrix. Tissue segmentation was performed by clustering super-pixels according to gene expression, which was then co-registered with the H&E-stained images. The top 10 genes defining each cluster were selected for biological interpretation and further refined by a pathologist based on tissue morphology.

### Integration of CosMx and CODEX data

The alignment between the CosMx data and the CODEX data is performed using STalign with default parameters^69^. The CODEX data is normalized using centered log ratio transformation from Seurat^102^. The senescent cells are identified in the CODEX data as cells whose expression levels of p16 and p21 > 0.5. For the CosMx data, we focused on a subset of the region (24 FOVs) and normalized the data using log normalization from Seurat. We identified B cells through Louvain clustering (resolution = 1), and we computed the Euclidean distances of the senescent cells to these B cells using the aligned spatial coordinates and constructed a k nearest neighbor graph between the senescent cells and the B cells (k = 100). B cells are categorized into 3 groups: B cells that are the first nearest neighbors to the senescent cells (senescence-adjacent B cells), B cells that are not nearest neighbors to the senescent cells (distant B cells), and the remaining B cells (others). Differential gene expression analysis is performed between the senescence adjacent B cells and the distant B cells using a two-sample t-test from Seurat (p values are adjusted using Bonferroni correction). Top differentially expressed genes are identified with adjusted p value < 0.01.

### Epigenome analysis

The Gene Score model in ArchR was used to generate the gene accessibility score. A gene score matrix was generated for downstream analysis. The getMarkerFeatures and getMarkers function in ArchR (testMethod= “wilcoxon”, cutOff= “FDR<=0.05”) was used to identify the marker regions/genes for each cluster, and the marker genes were discussed in the manuscript because they were identified as one of the top differential genes between clusters. Gene-score imputation was implemented with addImputeWeights for data visualization. Spatial mapping of gene scores for selected marker genes in different clusters and the chromatin accessibility at select genes are highly tissue specific. Cell-type-specific marker peaks were identified using getMarkerFeatures with parameters (bias = c (“TSSEnrichment”, “log10(nFrags)”, testMethod = “wilcoxon”)) and getMarkers (cutOff = “FDR <= 0.05 & Log2FC >= 0.1”). Metadata from cell annotation was achieved through Knn_ on harmony and from a created Seurat object, peaks were called using MACS2 via the Callpeaks function in Signac. To compute per-cell motif activity, AddMotifs was applied followed by linking the peaks to each desired gene, using LinkPeaks.

## Data and code availability

The raw data for this study has been submitted to The Cellular Senescence Network (SenNet) Data Portal (https://data.sennetconsortium.org/) under the data provider group "TMC - Yale University-Fan". The PhenoCycler data was uploaded with upload IDs SNT223.SPFP.359 and SNT354.FTHQ.327 for FFPE and fresh frozen samples, respectively. The DBiT-Seq data was uploaded with upload IDs SNT347.QGGP.668 and SNT248.HNBH.289 for ATAC-Seq and RNA-Seq, respectively. Scripts for data analysis and visualization were written mostly in Python and R, available at https://github.com/MingyuYang-Yale/DBiT-seq, https://github.com/HaikuoLi/spatial_epigenome_FFPE.

---

**Table 1:** Summary of Human Lymph Node Tissue Procurement.

| Block numbers | Section | Age (yo) | Gender | Location | FF/FFPE | Modality |
| --- | --- | --- | --- | --- | --- | --- |
| YHLN-N6 | Yale- whole lymph node | 63 | M | right inguinal | FF | codex / spatial atac / spatial transcript |
| YHLN-N8 | Yale - whole lymph node | 29 | F | hilar | FF | codex / spatial atac / spatial transcript |
| YHLN-N9 | Yale - whole lymph node | 54 | F | submental | FF | codex / spatial atac / spatial transcript |
| YHLN-N17 | Yale - whole lymph node | 78 | F | left axillar | FF | codex / spatial atac / spatial transcript |
| YHLN-N2 | Yale - whole lymph node | 62 | M | left neck | FF | codex / spatial atac |
| YHLN-N22 | Yale - whole lymph node | 22 | F | left neck | FFPE | codex |
| YHLN-N27 | Yale - whole lymph node | 25 | F | right neck | FFPE | codex / spatial transcript |
| YHLN-N4 | Yale - whole lymph node | 70 | M | right inguinal | FF | - |
| YHLN-N1 | Yale - whole lymph node | 73 | F | left groin | FF | - |
| YHLN-N3 | Yale - whole lymph node | 68 | M | left neck | FF | codex / spatial atac |
| YHLN-N5 | Yale - whole lymph node | 62 | F | left tonsil | FF | - |
| YHLN-N7 | Yale - whole lymph node | 75 | F | left axillar | FF | - |
| YHLN-N12 | Yale - whole lymph node | 20 | M | right neck | FF | codex / spatial atac / spatial transcript |
| YHLN-N13 | Yale - whole lymph node | 71 | F | left inguinal | FF | codex / spatial atac / spatial transcript |
| YHLN-N14 | Yale - whole lymph node | 74 | M | left neck | FF | codex / spatial atac |
| YHLN-N15 | Yale - whole lymph node | 84 | F | right axillary | FF | - |
| YHLN-N16 | Yale - whole lymph node | 86 | M | left neck | FF | codex / spatial atac |
| YHLN-N18 | Yale - whole lymph node | 62 | M | right neck | FFPE | - |
| YHLN-N19 | Yale - whole lymph node | 81 | M | right base of tongue | FFPE | - |
| YHLN-N20 | Yale - whole lymph node | 66 | M | right axillary | FFPE | codex / spatial transcript |
| YHLN-N21 | Yale - whole lymph node | 45 | F | left axillar | FFPE | - |
| YHLN-N23 | Yale - whole lymph node | 52 | M | right neck | FFPE | codex / spatial transcript |
| YHLN-N24 | Yale - whole lymph node | 74 | F | right inguinal | FFPE | - |
| YHLN-N25 | Yale - whole lymph node | 74 | F | right inguinal | FFPE | - |
| YHLN-N26 | Yale - whole lymph node | 86 | M | left axillary | FFPE | codex / spatial transcript |
| YHLN-N28 | Yale - whole lymph node | 55 | M | right neck | FFPE | - |
| YHLN-N29 | Yale - whole lymph node | 34 | M | left neck | FFPE | codex / spatial transcript |
| YHLN-N30 | Yale - whole lymph node | 55 | F | left neck | FFPE | codex / spatial transcript |
| YHLN-N31 | Yale - whole lymph node | 22 | M | right axilla | FFPE | codex |
| YHLN-N32 | Yale - whole lymph node | 61 | M | right groin | FFPE | - |
| YHLN-N33 | Yale - whole lymph node | 63 | F | right axilla | FFPE | - |
| YHLN-N34 | Yale-TMA | 53 | F | left axillary lymph node | FFPE | codex |
| YHLN-N35 | Yale-TMA | 65 | F | right neck mass | FFPE | codex |
| YHLN-N36 | Yale-TMA | 46 | M | midline neck | FFPE | codex |
| YHLN-N37 | Yale-TMA | 57 | F | submental mass | FFPE | codex |
| YHLN-N38 | Yale-TMA | 56 | F | left level 2 lymph node | FFPE | codex |
| YHLN-N39 | Yale-TMA | 17 | M | left neck mass | FFPE | codex |
| YHLN-N40 | Yale-TMA | 47 | F | right neck mass | FFPE | codex |
| YHLN-N41 | Yale-TMA | 8 | M | left neck mass | FFPE | codex |
| YHLN-N42 | Yale-TMA | 45 | F | right external jugular lymph | FFPE | codex |
| YHLN-N43 | Yale-TMA | 16 | M | right sided submental mass | FFPE | codex |
| YHLN-N44 | Yale-TMA | 29 | F | right neck level 5 lymph node | FFPE | codex |
| YHLN-N45 | Yale-TMA | 29 | F | submental lymph nodes for lymphoma | FFPE | codex |
| YHLN-N46 | Yale-TMA | 32 | F | left level 2 lymph node | FFPE | codex |
| YHLN-N47 | Yale-TMA | 47 | M | right level 2 neck mass | FFPE | codex |
| YHLN-N48 | Yale-TMA | 50 | F | left groin lymph node 2 | FFPE | codex |
| YHLN-N49 | Yale-TMA | 6 | F | left cervical lymph node | FFPE | codex |
| YHLN-N50 | Yale-TMA | 83 | M | colon | FFPE | codex |
| YHLN-N51 | Yale-TMA | 60 | F | right colon | FFPE | codex |
| YHLN-N52 | Yale-TMA | 85 | M | level 7 lymph node | FFPE | codex |
| YHLN-N53 | Yale-TMA | 81 | F | right 10 node | FFPE | codex |
| YHLN-N54 | Yale-TMA | 58 | M | mesenteric lymph node | FFPE | codex |
| YHLN-N55 | Yale-TMA | 71 | M | pancreas and spleen | FFPE | codex |
| YHLN-N56 | Yale-TMA | 68 | F | left axilla lymph node | FFPE | codex |
| N57 | Commercial-TMA | 30 | M | NA | FFPE | codex |
| N58 | Commercial-TMA | 34 | M | NA | FFPE | codex |
| N59 | Commercial-TMA | 30 | M | NA | FFPE | codex |
| N60 | Commercial-TMA | 40 | M | NA | FFPE | codex |
| N61 | Commercial-TMA | 32 | M | NA | FFPE | codex |
| N62 | Commercial-TMA | 35 | M | NA | FFPE | codex |
| N63 | Commercial-TMA | 28 | M | NA | FFPE | codex |
| N64 | Commercial-TMA | 38 | M | NA | FFPE | codex |
| N65 | Commercial-TMA | 30 | M | NA | FFPE | codex |
| N66 | Commercial-TMA | 35 | M | NA | FFPE | codex |
| N67 | Commercial-TMA | 36 | M | NA | FFPE | codex |
| N68 | Commercial-TMA | 31 | M | NA | FFPE | codex |
| N69 | Commercial-TMA | 28 | M | NA | FFPE | codex |
| N70 | Commercial-TMA | 31 | M | NA | FFPE | codex |
| N71 | Commercial-TMA | 33 | M | NA | FFPE | codex |
| N72 | Commercial-TMA | 40 | M | NA | FFPE | codex |
| N73 | Commercial-TMA | 28 | M | NA | FFPE | codex |
| N74 | Commercial-TMA | 32 | M | NA | FFPE | codex |
| N75 | Commercial-TMA | 32 | M | NA | FFPE | codex |
| N76 | Commercial-TMA | 45 | M | NA | FFPE | codex |
| N77 | Commercial-TMA | 35 | M | NA | FFPE | codex |
| N78 | Commercial-TMA | 79 | F | NA | FFPE | codex |
| N79 | Commercial-TMA | 42 | M | NA | FFPE | codex |
| N80 | Commercial-TMA | 56 | M | NA | FFPE | codex |

**Supplementary Table 2:** CODEX Antibody panel.

| Antibody | Antibody_rrid | Dilution_factor | Reporter_cat_number |
| --- | --- | --- | --- |
| Bcl-2 | AB_2936078 | 200 | 4550089 |
| Beta-actin | AB_3478101 | 200 | 4450040 |
| CD107a | AB_3474468 | 200 | 4550098 |
| CD11c | AB_3083459 | 200 | 4550114 |
| CD14 | AB_3083457 | 400 | 4450047 |
| CD141 | AB_3082975 | 200 | 4250097 |
| CD163 | AB_2935895 | 200 | 4250079 |
| CD20 | AB_3094498 | 200 | 4450094 |
| CD21 | AB_3474900 | 400 | 4450027 |
| CD31 | AB_2915935 | 100 | 4450017 |
| CD34 | AB_2909512 | 100 | 4250057 |
| CD38 | AB_3082976 | 200 | 4250080 |
| CD39 | AB_3096410 | 200 | 4250076 |
| CD3e | AB_2936080 | 200 | 4550119 |
| CD4 | AB_3094499 | 200 | 4550112 |
| CD44 | AB_2936081 | 400 | 4450041 |
| CD45 | AB_2915946 | 400 | 4450042 |
| CD45RO | AB_2895053 | 200 | 4250023 |
| CD66 | AB_3475664 | 200 | 4550001 |
| CD68 | AB_2935894 | 400 | 4550113 |
| CD8 | AB_2915960 | 200 | 4250012 |
| CDKN2A/p16 | AB_2078303 | 75 | 10883-1-AP |
| p21 | AB_3099451 | 200 | 19399 |
| CXCL13 | AB_2746222 | 400 | 38738A03 |
| CXCR5 | AB_394324 | 200 | 552032 |
| Collagen IV | AB_2927676 | 200 | 4550122 |
| E-cadherin | AB_2895057 | 200 | 4250021 |
| EpCAM | AB_2935888 | 200 | 4550088 |
| FOXP3 | AB_2927679 | 200 | 4550071 |
| Granzyme B | AB_3472025 | 200 | 4250055 |
| HLA-A | AB_3094501 | 200 | 4250100 |
| HLA-DR | AB_3080864 | 200 | 4550118 |
| HLA-E | AB_3478569 | 200 | 4250065 |
| HMGB1 | AB_2049739 | 150 | ab92310 |
| ICOS | AB_3096408 | 200 | 4550117 |
| IDO1 | AB_3476035 | 200 | 4550123 |
| IFN-G | AB_3476455 | 200 | 4250062 |
| Ki67 | AB_3094497 | 200 | 4450096 |
| LAG3 | AB_3096409 | 100 | 4550058 |
| LMP1 | AB_1566182 | 50 | ab78113 |
| MPO | AB_2927678 | 200 | 4250083 |
| Mac2/Galectin-3 | AB_3477119 | 200 | 4450034 |
| PCNA | AB_2936083 | 200 | 4550124 |
| PD-L1 | AB_3096406 | 100 | 4550072 |
| PD1 | AB_3096407 | 200 | 4550038 |
| Pan-Cytokeratin | AB_3083456 | 200 | 4450020 |
| Podoplanin | AB_3082979 | 200 | 4250094 |
| SMA | AB_2936084 | 200 | 4450049 |
| TOX | AB_3477620 | 200 | 4250067 |
| VISTA | AB_3479195 | 150 | 4250063 |
| Vimentin | AB_2935889 | 200 | 4450050 |
| yH2AX | AB_315795 | 300 | 613402 |

**Supplementary Table 3:**
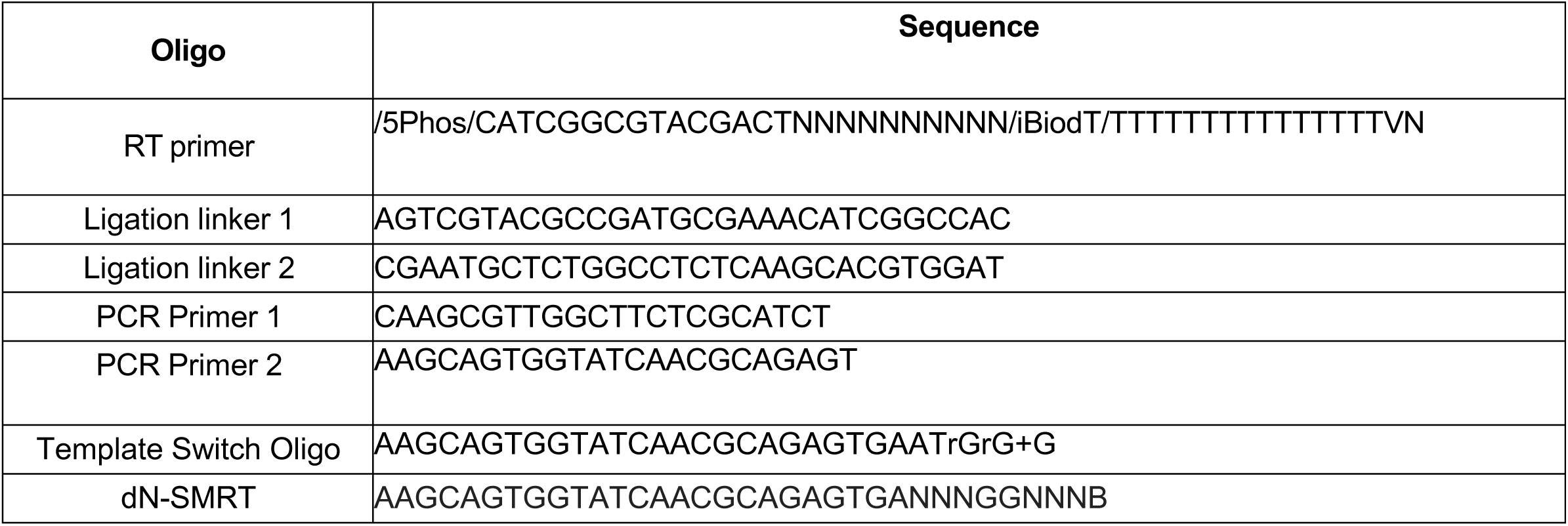
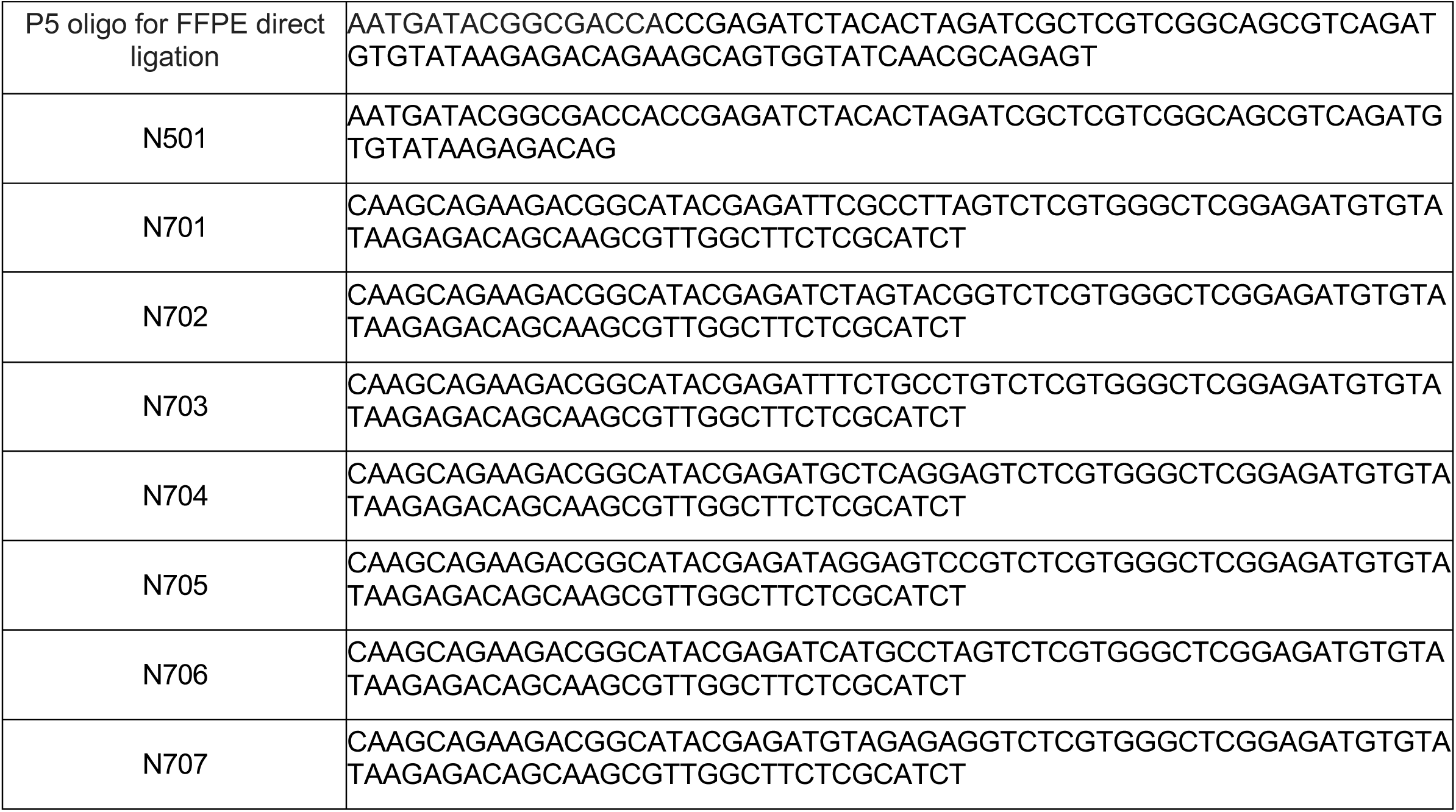
DNA oligos for PCR, ligation and library preparation- spatial transcriptome.

**Supplementary Table 4:**
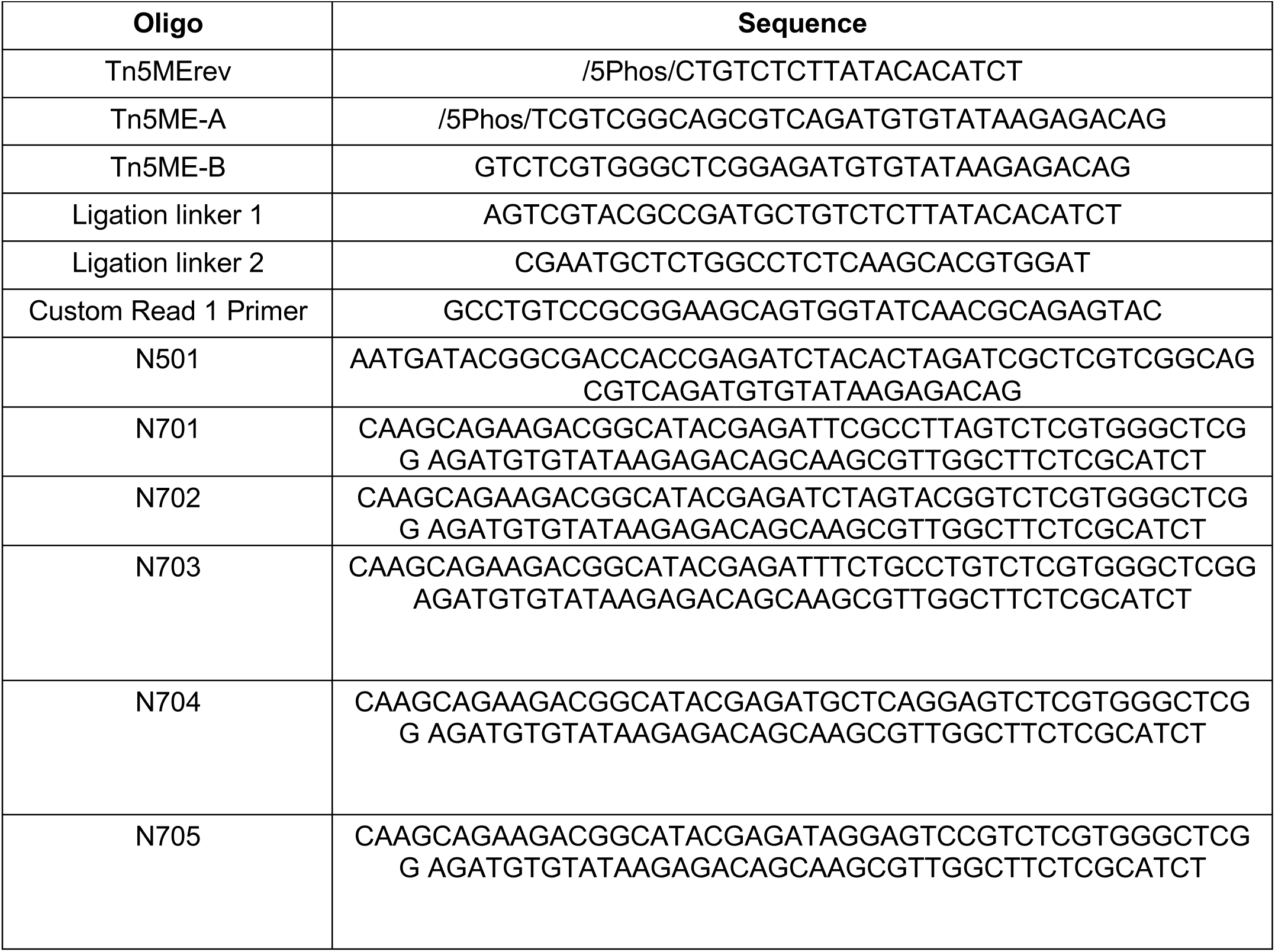
DNA barcode B sequences- spatial ATAC-seq.

**Supplementary Table 5:**
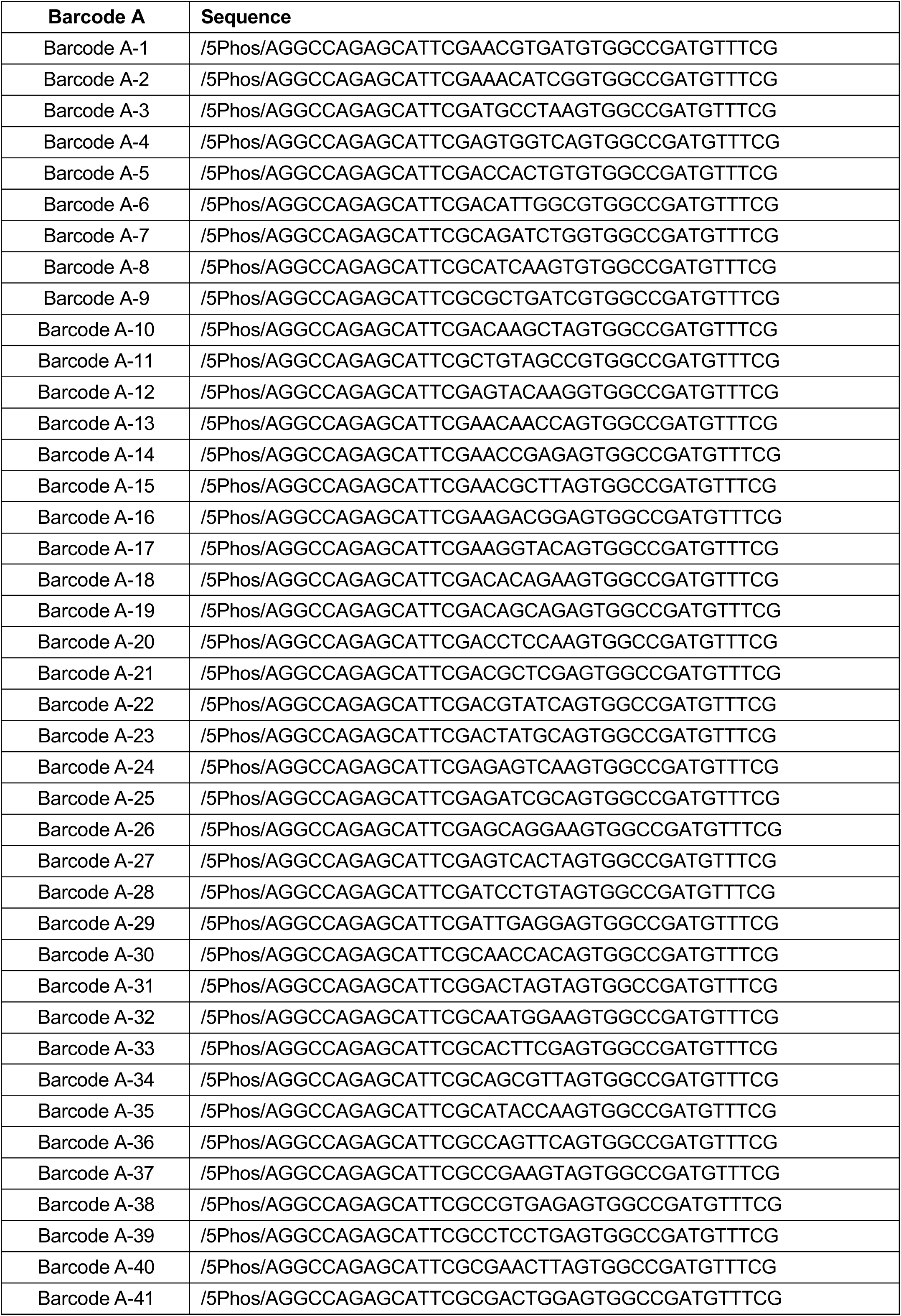
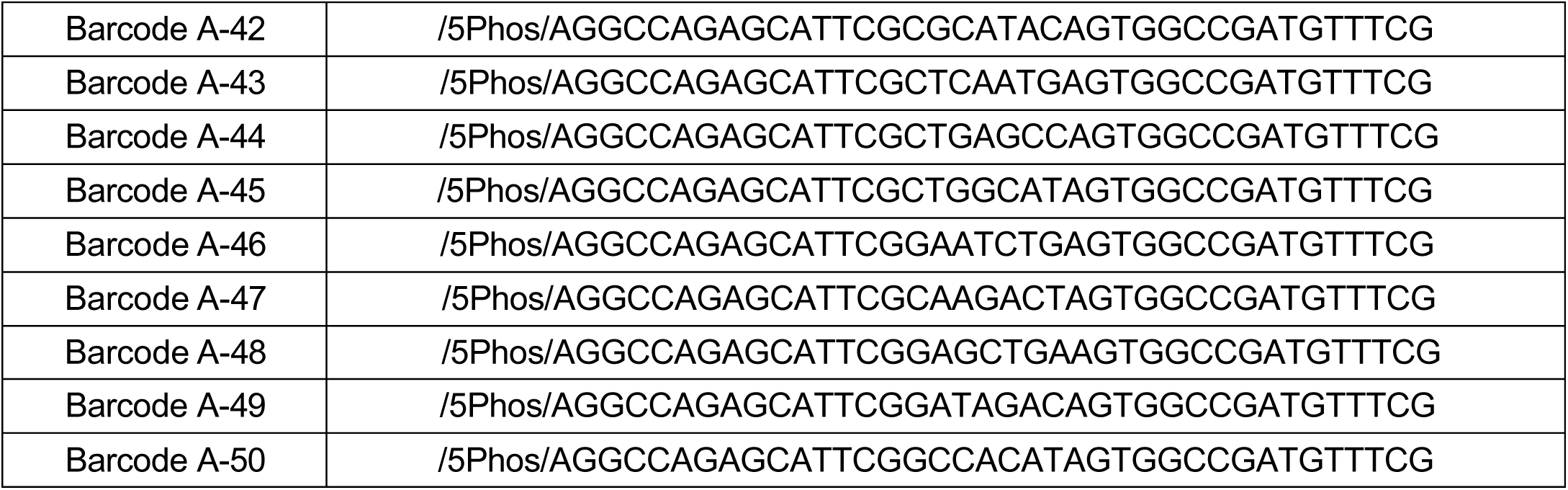
DNA barcode A sequences.

**Supplementary Table 6:**
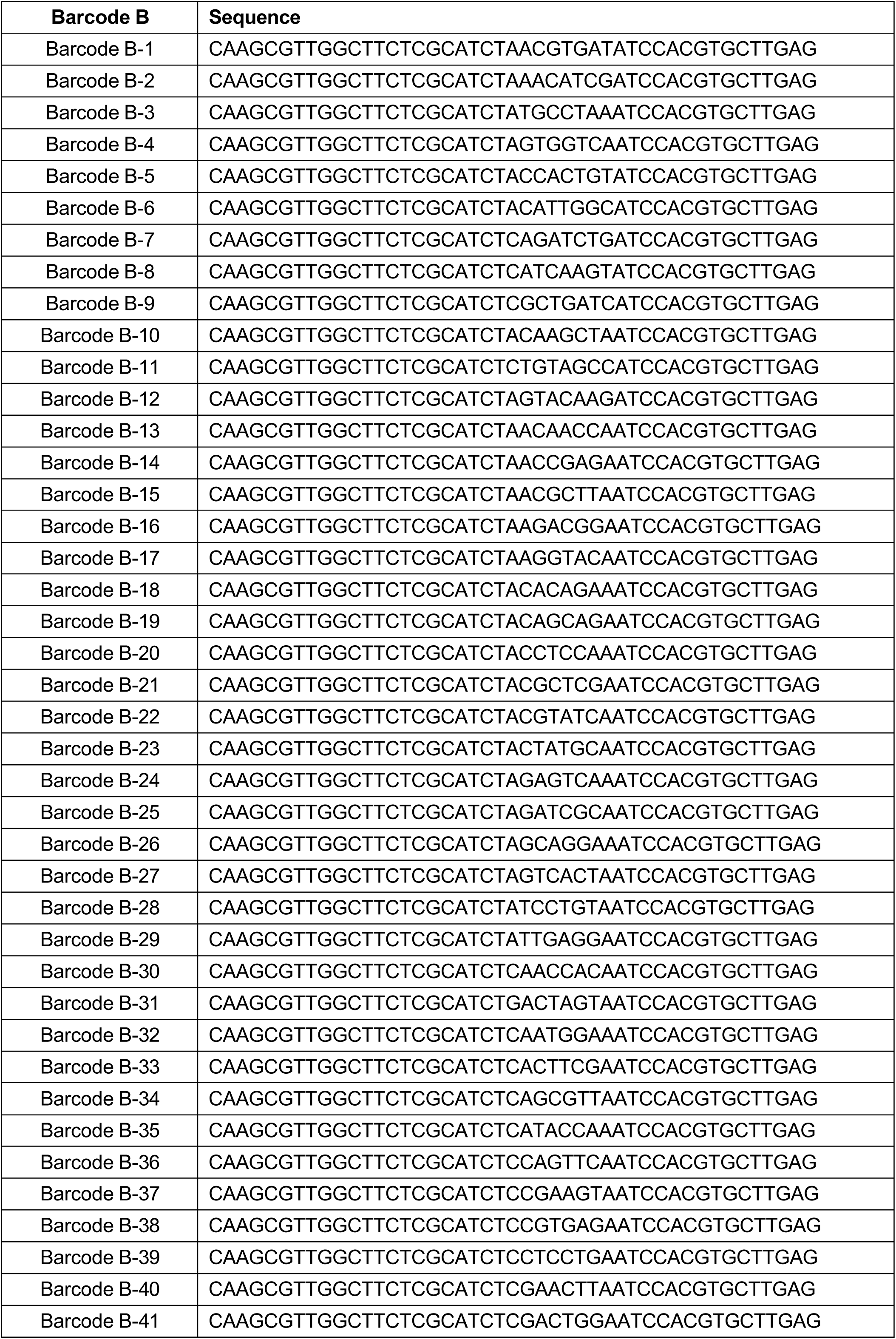
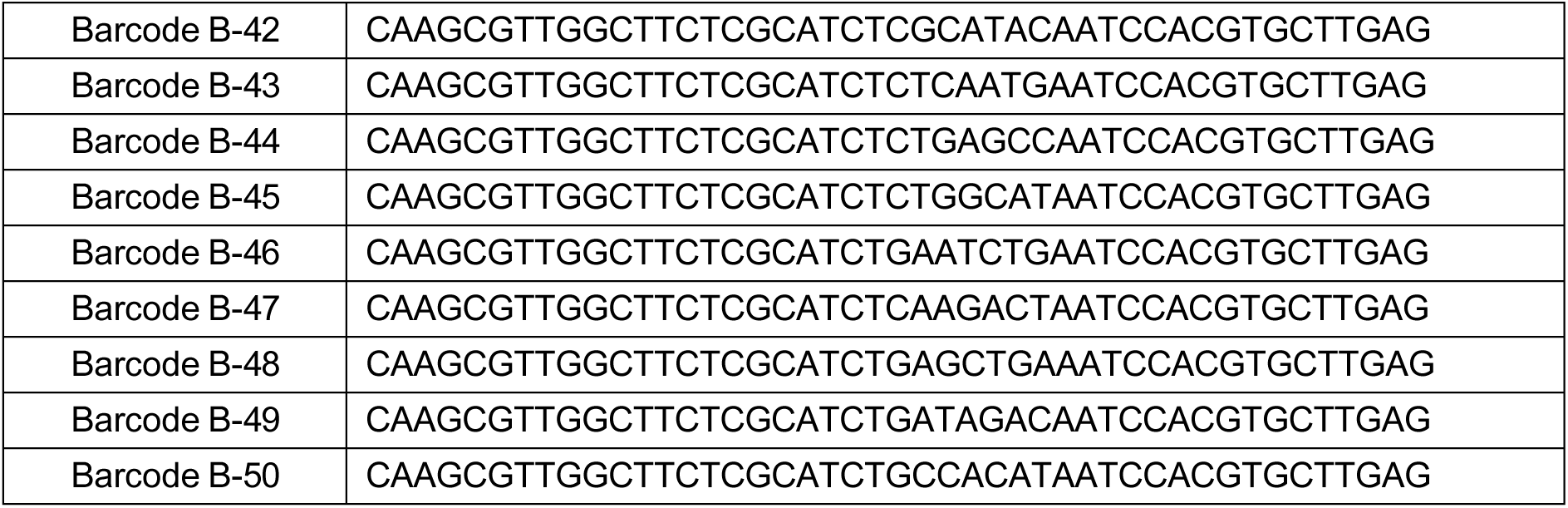
DNA barcode B sequences.

**Supplementary Table 6:**
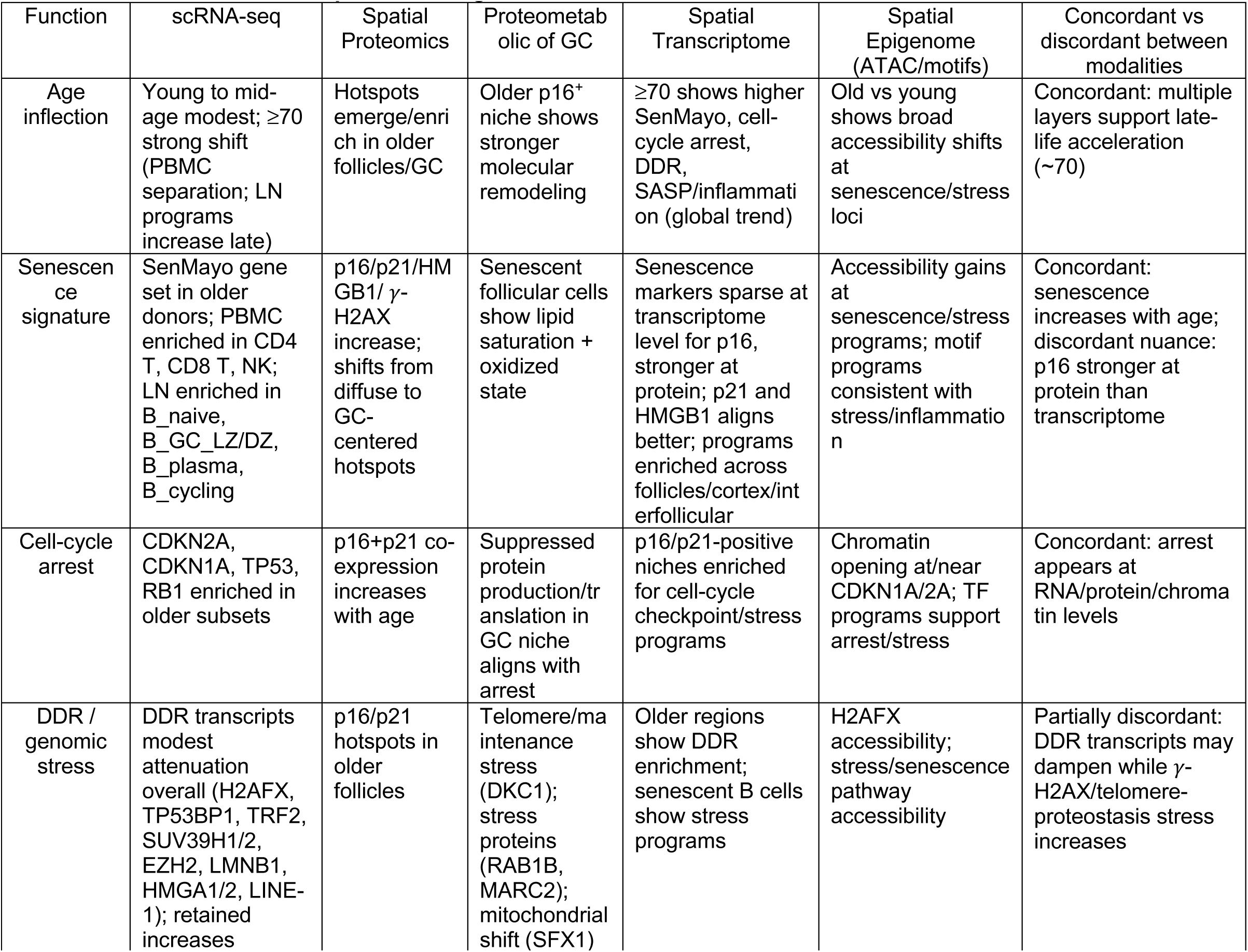
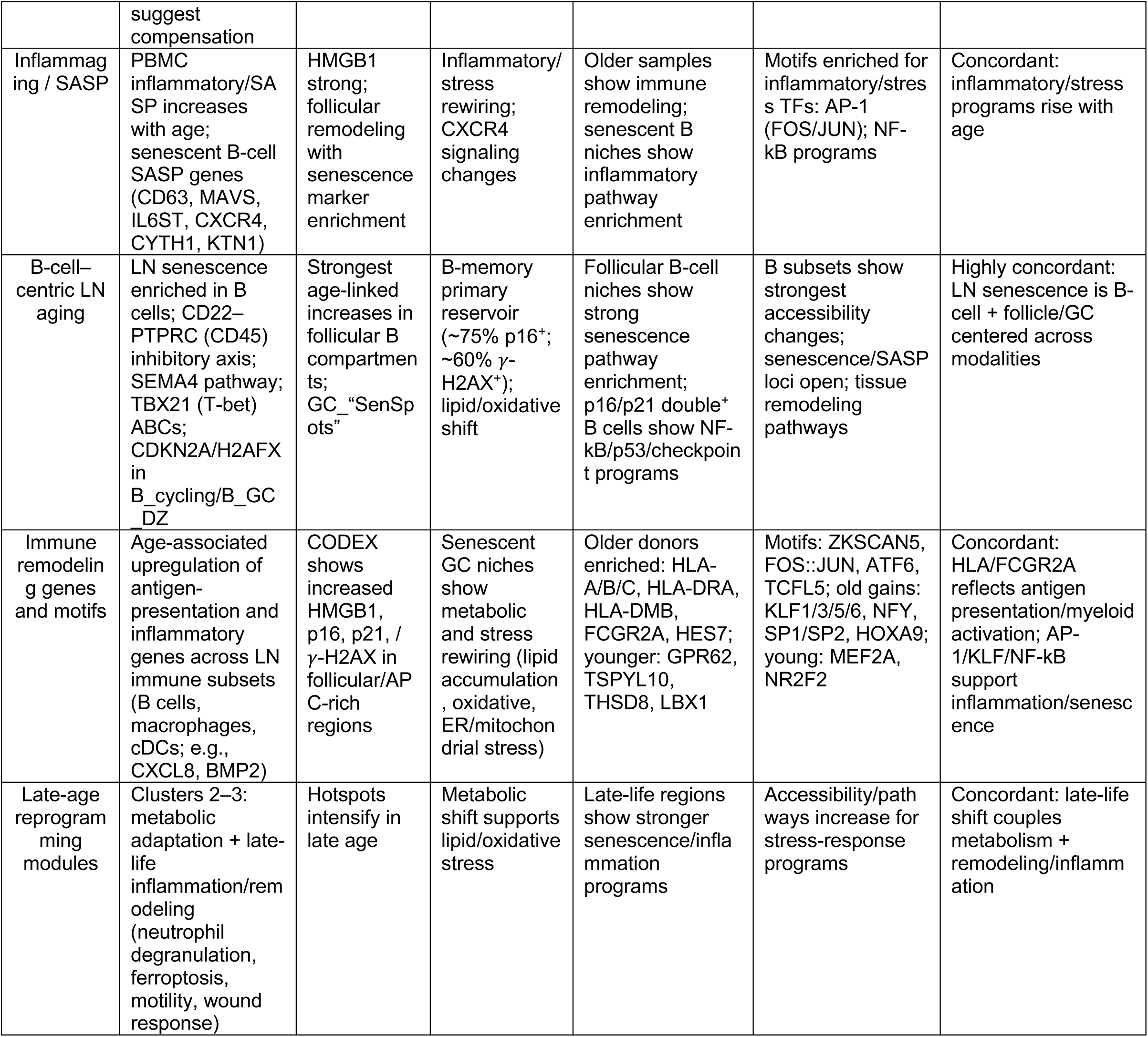
Concordant and discordant features of senescence and aging programs across multi-modal spatial and single-cell datasets.

**Figure S1.**
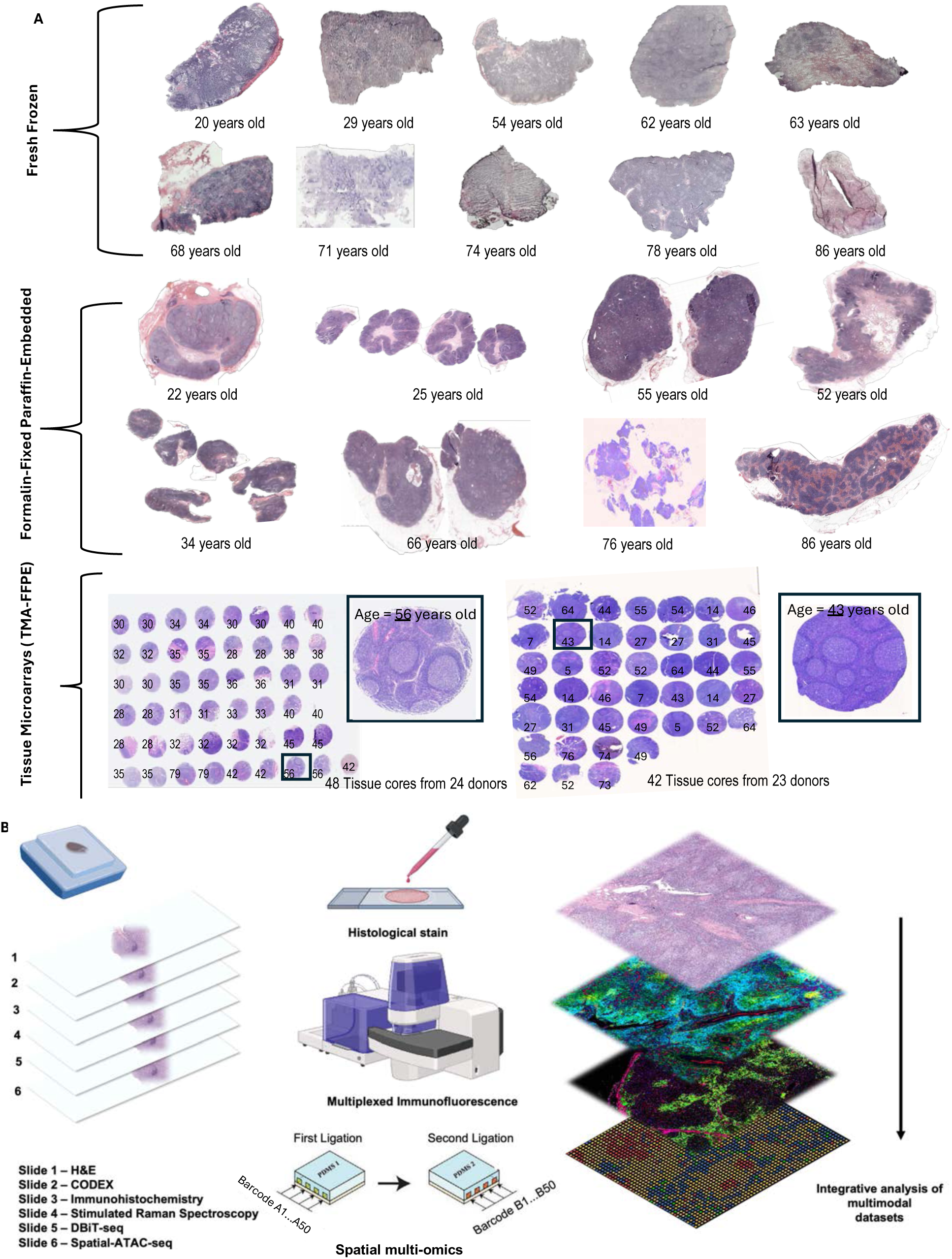
Overview of human lymph node samples and multimodal spatial profiling workflow. (A) Representative hematoxylin and eosin (H&E)-stained images of human lymph node sections collected from donors of varying ages, displayed by preservation method. Fresh frozen samples (top) include donors aged 20–86 years. Formalin-fixed paraffin-embedded (FFPE) samples (middle) include donors aged 22–86 years. Tissue microarrays (TMAs, bottom) consist of multiple cores from 47 donors (ages indicated), with representative enlarged cores. (B) Schematic of the integrative multimodal spatial profiling workflow. Serial tissue sections were processed for histological staining (Slide 1: H&E), multiplexed immunofluorescence imaging (Slide 2: CODEX), immunohistochemistry (Slide 3), stimulated Raman spectroscopy (Slide 4), DBiT-seq (Slide 5), and spatial-ATAC-seq (Slide 6). Spatial barcoding and ligation enable multi-omics data integration for comprehensive analysis of cellular and molecular features within human lymph node tissues.

**Figure S2.**
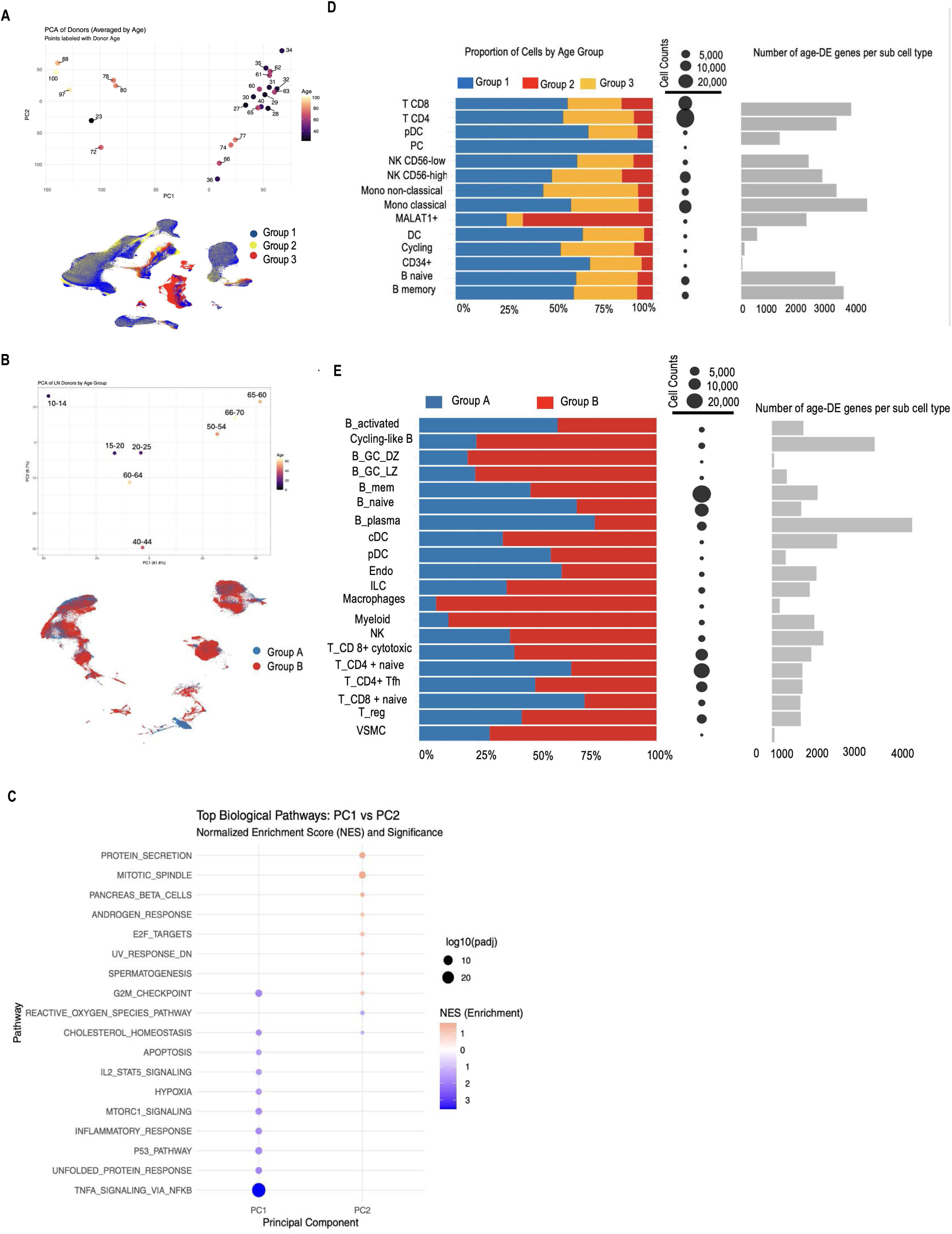
Single cell-RNA-seq data set with district sub cell types and across different age group. (A,B) Principal component analysis (PCA) of PBMC and LN donors across ages shows segregation along PC1 and PC2 , with age-associated separation across samples. UMAP projections colored by age group (Gorup A and B in LN , Group 1, Group 2, Group 3 in pbmc) illustrate the distribution of cells in each group. (C ) Dot plot showing top biological pathways enriched along PC1 and PC2 based on normalized enrichment score (NES). Dot size represents statistical significance (log10 adjusted p-value), and color indicates enrichment direction and magnitude. (D&E)Stacked bar plots display the proportion of cell types for each dataset across age groups (Group 1–3). Dot size indicates total cell counts per cell type, and horizontal bar plots show the number of age-associated differentially expressed genes (DEGs) per sub–cell type.

**Figure S3.**
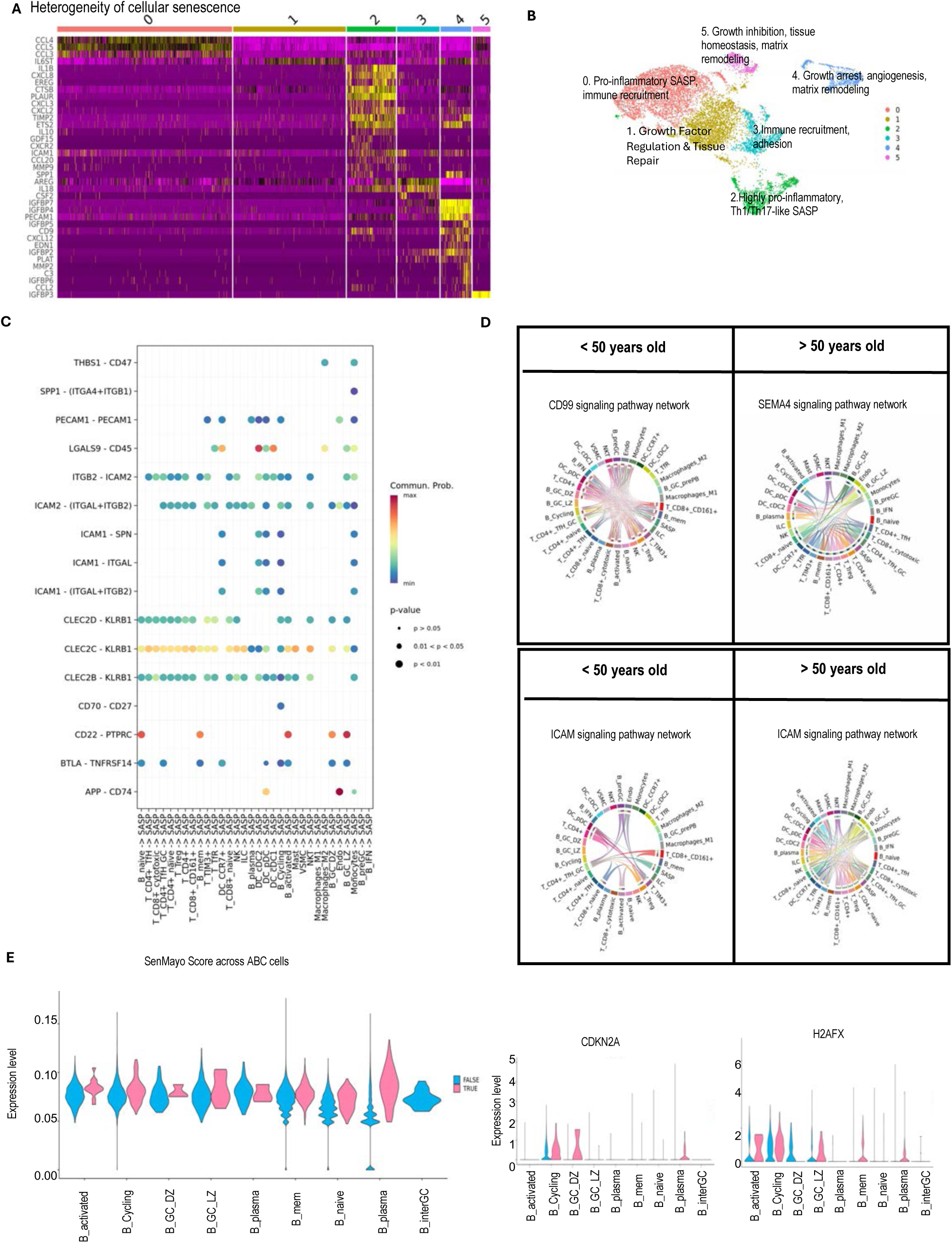
Cellular heterogeneity, ligand-receptor interactions, and age-associated changes in senescent B cell networks. (A) Heatmap showing heterogeneity of cellular senescence across clusters, highlighting differential expression of senescence-associated genes in distinct immune cell populations. (B) UMAP plot depicting six major senescent cell clusters annotated by their dominant functional signatures. (C) Dot plot showing significant ligand–receptor interactions between SASP cells and other immune cell types in donors aged more than 50 years. The color scale represents communication probability, and dot size corresponds to p-value significance. (D) Chord diagrams comparing age-associated differences in signaling pathway networks (<50 vs. >50 years old), illustrating enrichment of CD99, SEMA4, and ICAM signaling pathways in each group. (E) Violin plots displaying SenMayo module scores and expression levels of CDKN2A (p16) and H2AFX across age-associated B cell subsets, showing increased senescence signatures in specific B cell populations in aged individuals.

**Figure S4.**
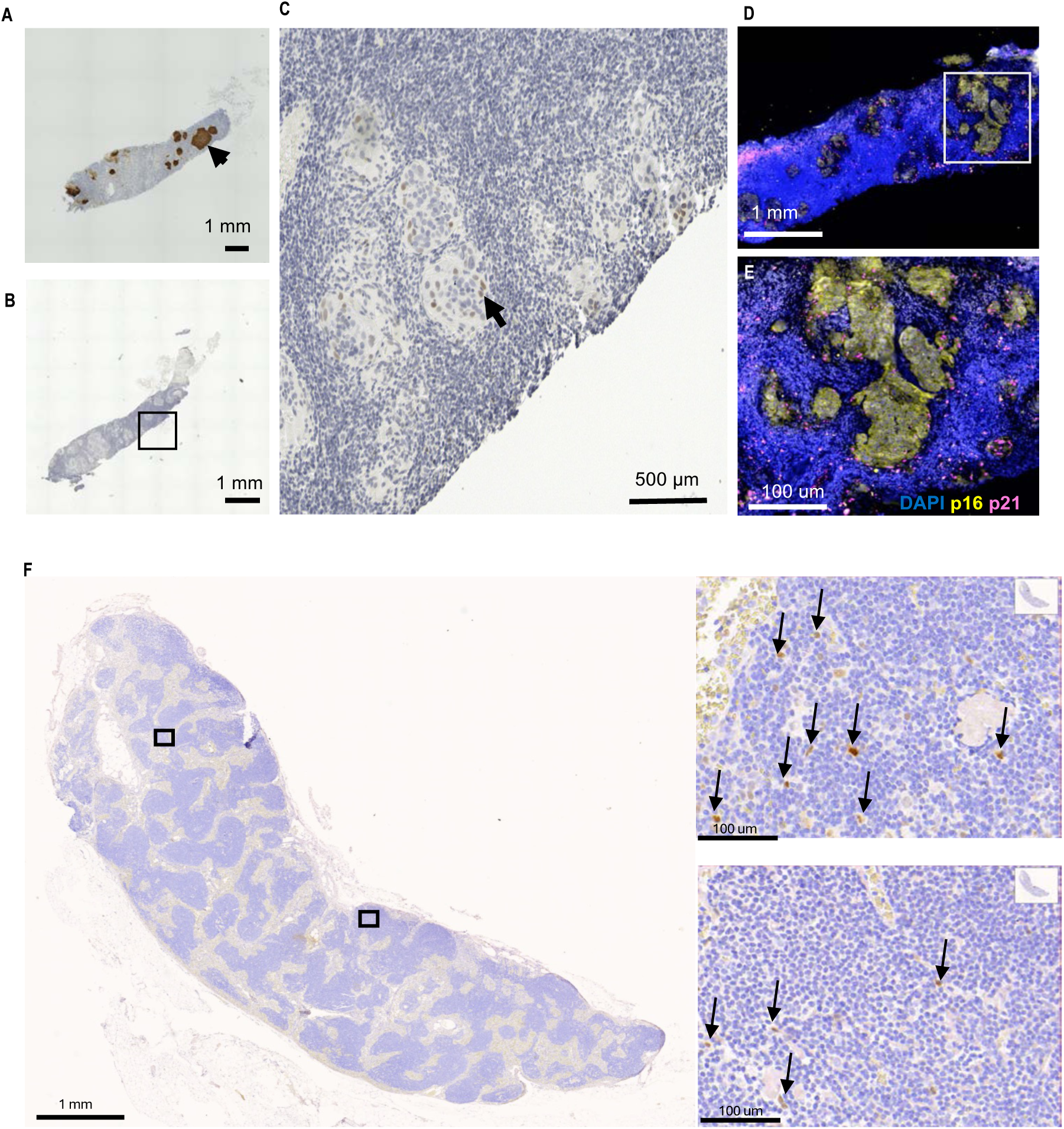
Validation of p16 and p21 antibodies on a p16 positive nasopharyngeal carcinoma and human lymph node samples. (A–C) Immunohistochemistry (IHC) images showing p16⁺ cells (brown, arrows) within lymph node sections at low (A, B) and higher (C) magnification. (D–E) Multiplex immunofluorescence images displaying co-localization of p16 (yellow) and p21 (magenta) within tumor regions, counterstained with DAPI (blue). (F) Representative IHC images of p21 expression across the entire human lymph node section of an old donor-86 years old (left), with magnified insets (right) showing clusters of p21⁺ cells (arrows) localized predominantly in germinal center and perifollicular regions. Scale bars are indicated.

**Figure S5.**
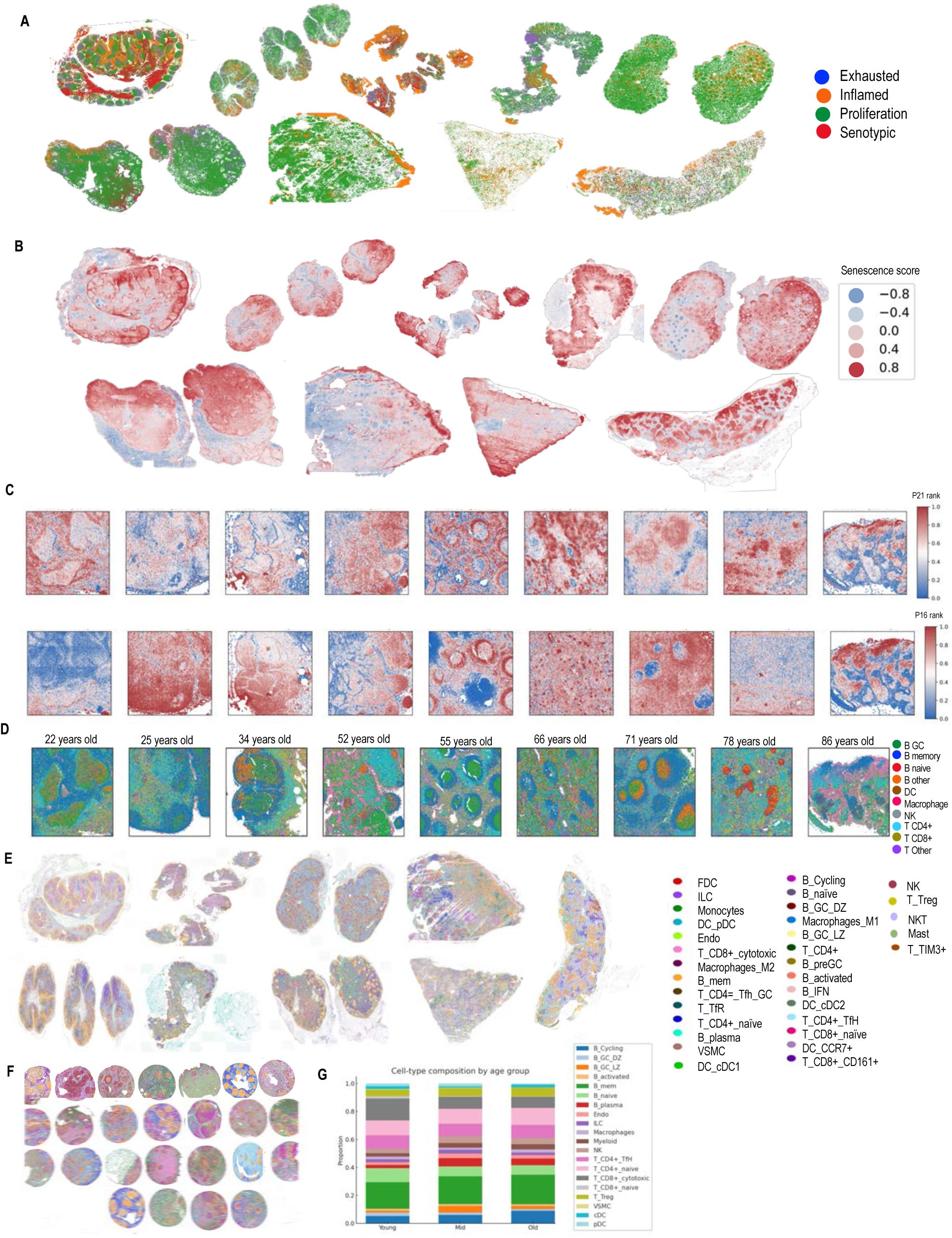
Spatial proteomic mapping of senescence, proliferation, and immune cell composition in human lymph nodes across ages. (A) Spatial distribution of cellular states (exhausted, inflamed, proliferative, and senescent) within lymph node sections, highlighting heterogeneity across donors. (B) Spatial proteomic maps senescence score (red) and relative rank across donors, with localized to germinal center regions in 50-70 and >70 years old donors. (C) Representative high-magnification images illustrating spatial patterns of CDKN2A (p16) and CDKN1A (p21) expression within zoomed in follicles of individual lymph nodes. (D) CODEX multiplexed immunofluorescence images of lymph node sections from donors aged 22–86 years, displaying cell-type distribution and senescent cell localization. (E) Integrated cell-type annotation of lymph node sections combining multiplexed protein and scRNA-seq reference data, identifying major immune cell populations.

**Figure S6.**
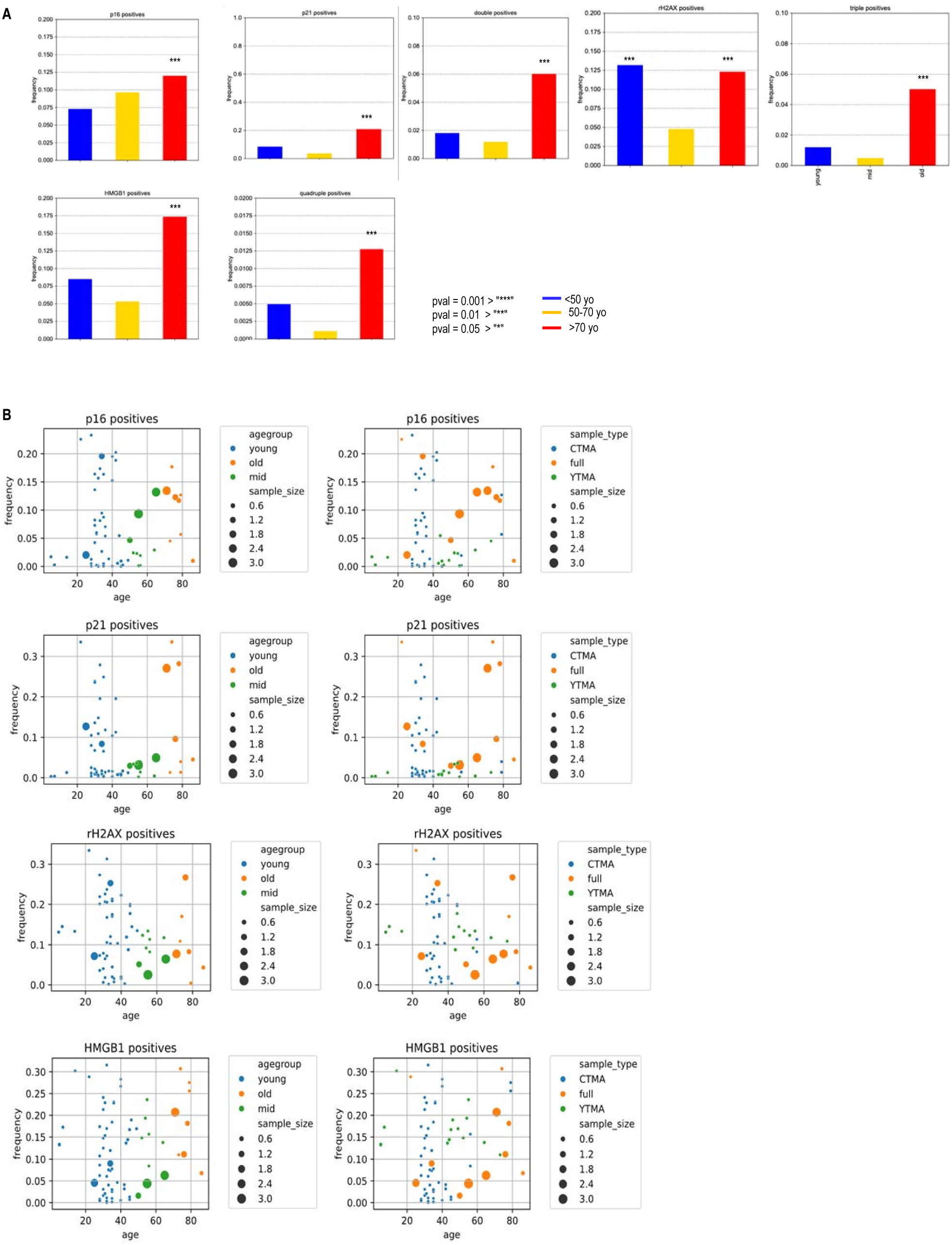
Quantification of senescence marker-positive cells in human lymph nodes across age groups using CODEX. (A) Bar plots showing the frequency of cells positive for p16, p21, double-positive (p16+p21), 𝛾-H2AX, triple-positive (p16+p21+ 𝛾-H2AX+or HMGB1+), HMGB1, and quadruple (p16+p21+ 𝛾-H2AX+HMGB1+) across three age groups (<50 , 50-70, and >70 years old). Statistical significance is indicated (**p ≤ 0.01; ***p ≤ 0.001). (B) Scatter plots depicting the frequency of p16⁺, p21⁺, 𝛾-H2AX⁺, and HMGB1⁺ cells as a function of donor age. Data points are color-coded by age group (<50 , 50-70, and >70 years old). and shaped by sample type (CTMA, full, YTMA), with point size indicating sample size (cells in millions).

**Figure S7.**
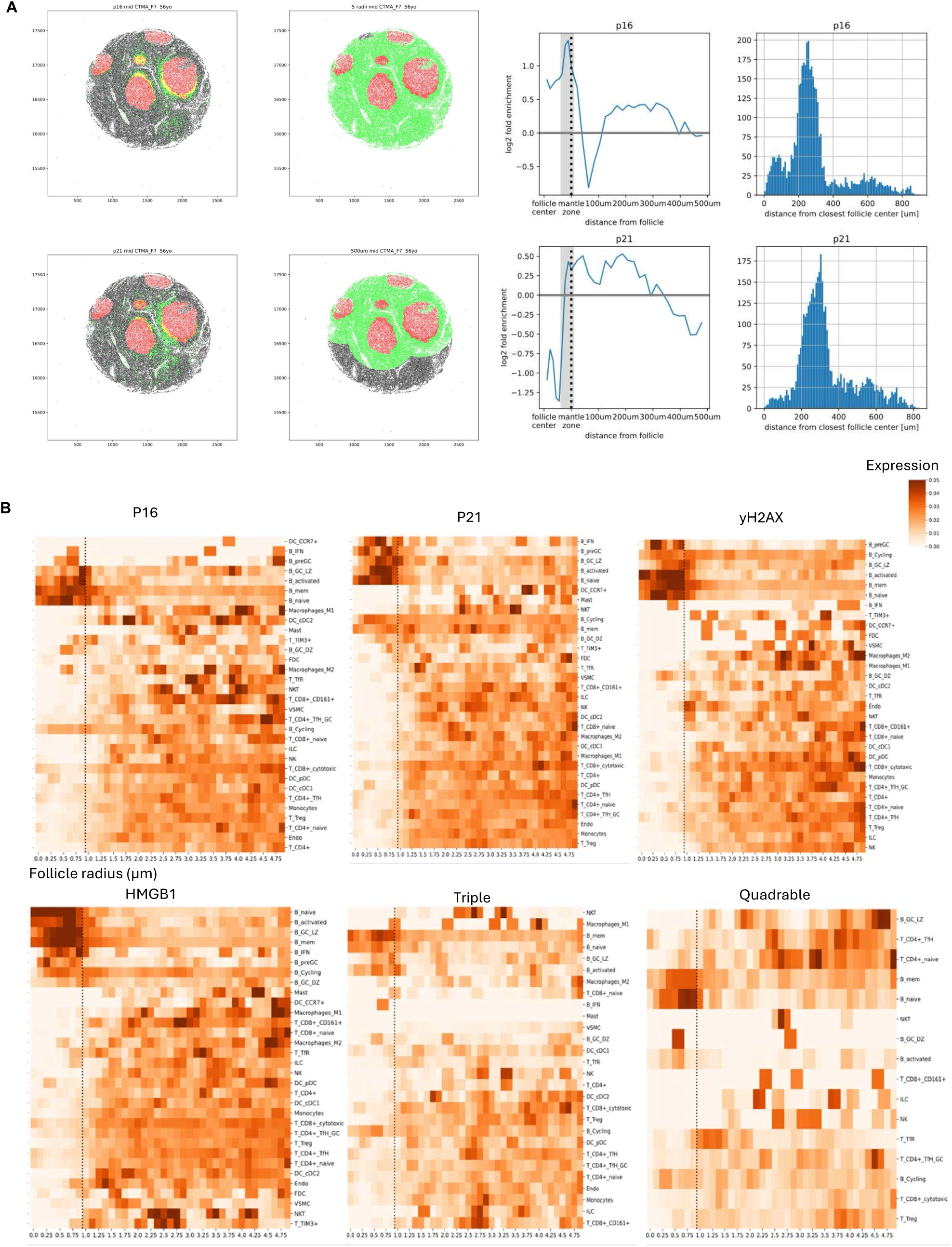
Spatial enrichment of senescence marker-positive cells relative to follicular structures. (A) Representative images showing follicle identification on a mid-age TMA (56 years old) based on sample-specific CD20 and CD21 expression. For each follicle, a Gaussian mixture model was applied to determine centroid location and morphology. Line plots depict fold-enrichment of p16⁺ and p21⁺ cells relative to distance from follicle centers in both Mahalanobis and Euclidean space. Histograms on the right show the distribution of distances of p16⁺ and p21⁺ cells from the nearest follicle center. (B) Heatmaps depicting the abundance of cells positive for p16, p21, 𝛾-H2AX, HMGB1, triple-positive, and quadruple-positive markers as a function of follicular radius (x-axis) across various cell types (y-axis).

**Figure S8.**
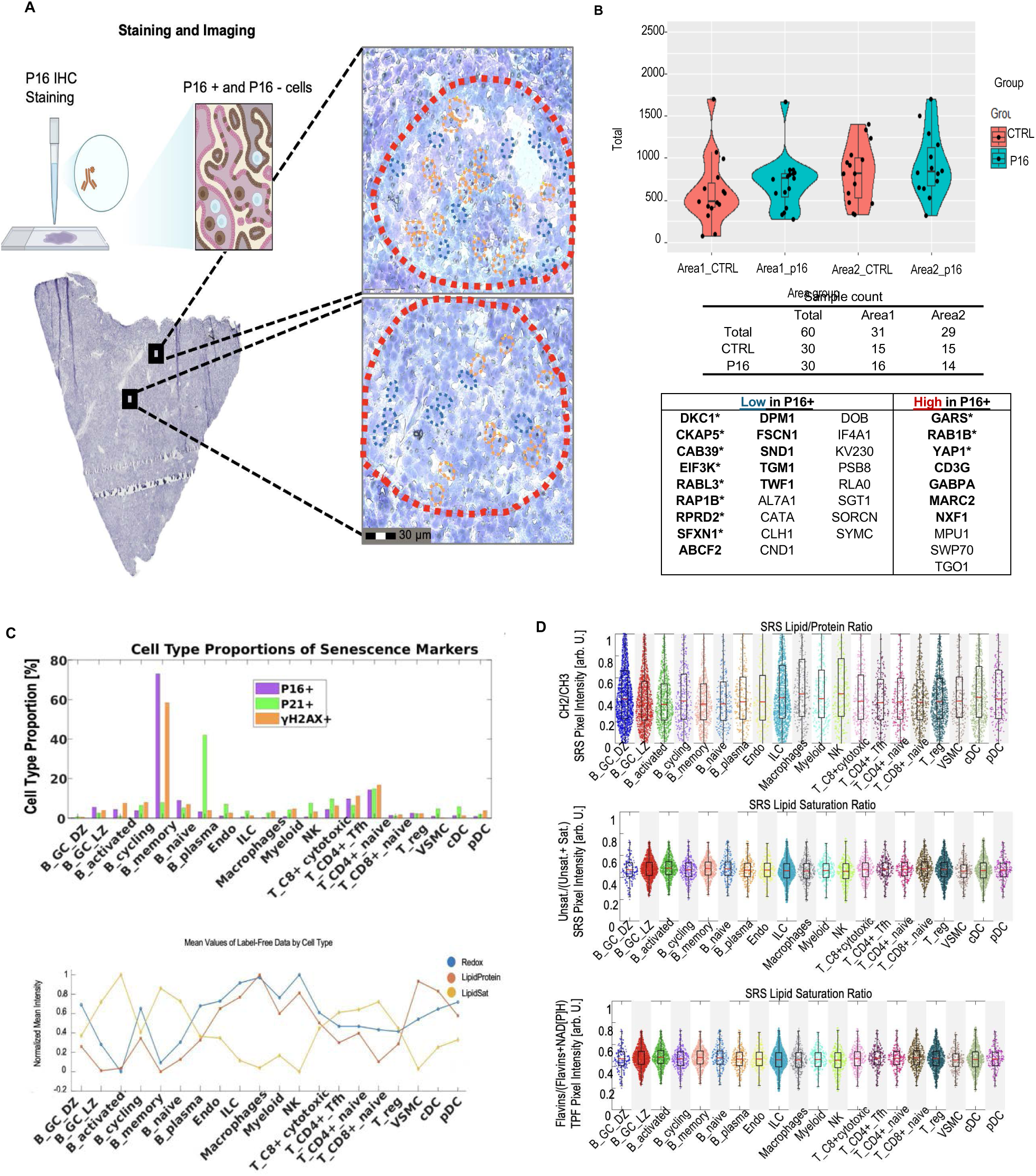
Integrated spatial single-cell proteomics and SRS+CODEX profiling of senescent cells in human lymph nodes. (A) Human lymph node tissue section from a 78-year-old donor was mounted on a Polyethylene Naphthalate (PEN) membrane for spatial single-cell proteomics. Workflow for isolating single p16⁺ and p16⁻ cells from a fresh frozen human lymph node section (78-year-old donor) mounted on a PEN membrane. Sections were stained with p16-IHC, and target follicles were identified by overlapping p16-IHC images with serial sections stained using a multiplex immune panel (including p16 and p21). Single cells were captured by laser ablation and deposited into nanoPOTS wells, followed by proteomic preparation and LC-MS/MS analysis. Representative images show selected follicles with p16⁺ (orange) and p16⁻ (blue) cells. (B) Violin plots comparing the median number of quantified protein groups per cell for p16⁺ and p16⁻ cells across two follicular regions. The table lists proteins with significantly different abundance between p16⁺ and p16⁻ cells, as identified by ANOVA, with key proteins enriched or depleted in p16⁺ cells highlighted. (C) Bar plots showing cell type proportions of p16⁺, p21⁺, and 𝛾-H2AX⁺ senescent cells. The line plot below shows mean label-free SRS-derived metabolic features (lipid-to-protein ratio, lipid saturation, and lipid unsaturation) across cell types. (D) Violin plots displaying SRS-derived lipid/protein ratio, lipid saturation ratio, and lipid unsaturation ratio for major immune cell subsets, overlaid with CODEX-defined cell types.

**Figure S9.**
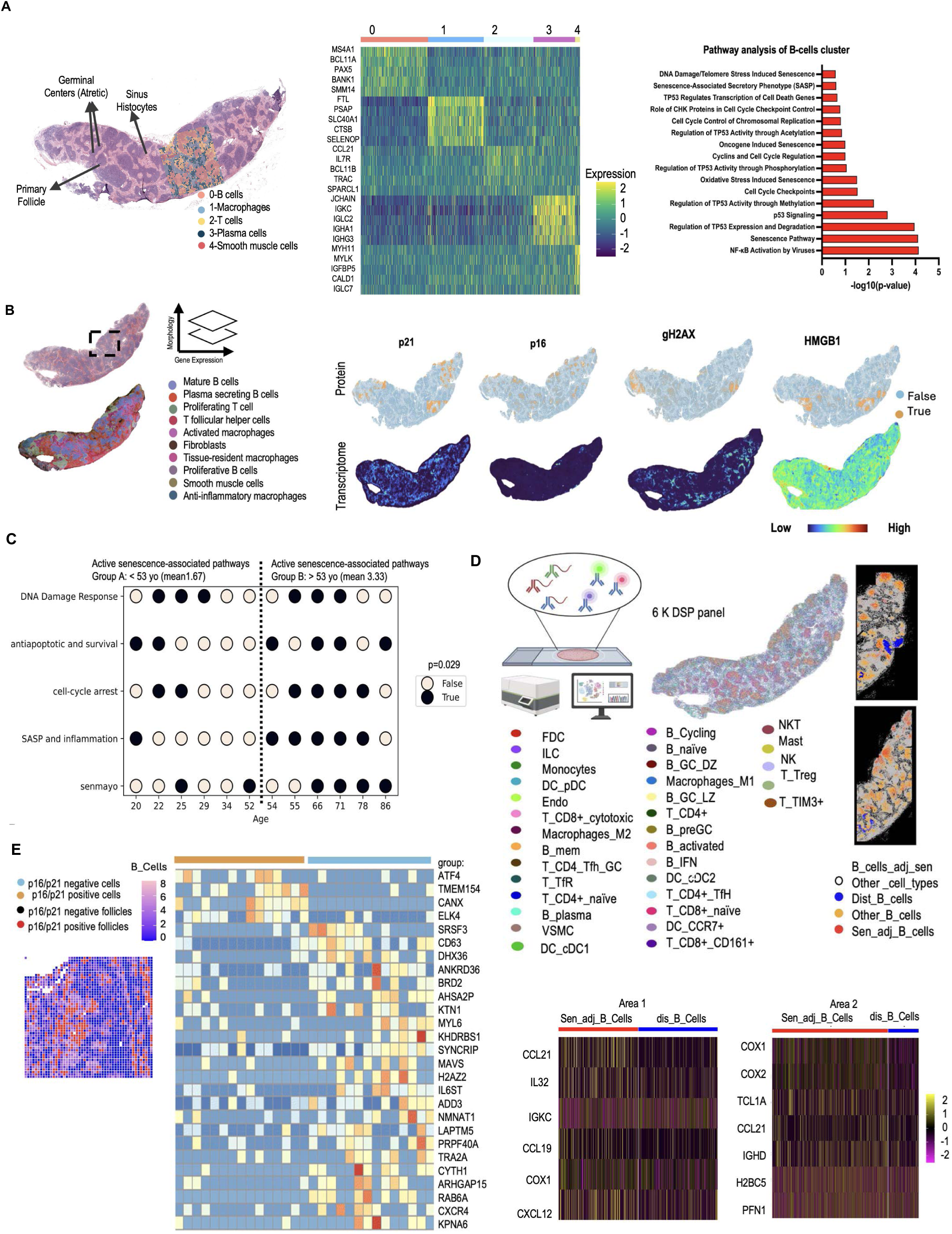
Integration of DBiT-seq, CosMx, and CODEX for spatial and transcriptomic characterization of senescent B cells. (A) H&E image annotation of lymph node regions (germinal centers, sinuses, primary follicles) overlaid with DBiT-seq transcriptomic profiling. The heatmap shows gene expression in clusters, and pathway analysis highlights pathway enrichment in the B-cells cluster. (B) Super-resolved transcriptome-wide expression was generated from the DBiT-seq profile using iSTAR and compared with the multiplexed protein profile. Expression patterns of p21, p16, 𝛾-H2AX, and HMGB1 were analyzed across the entire lymph node from an 86-year-old donor. (C) Dot plot summarizing pathway-level senescence activity across individual donors ordered by age. For each donor, five senescence-related programs (DNA damage response, anti-apoptotic/survival signaling, cell-cycle arrest, SASP/inflammation, and SenMayo) were binarized as active (True, dark) if the pathway score ranked above the cohort median, or inactive (False, light) otherwise. (D) Integration of CosMx spatial transcriptomics with scRNA-seq reference using MaxFuse, resulting in single-cell level annotation of immune populations. CosMx spatial analysis comparing p16/p21-positive and -negative B cells within follicles. DE gene heatmaps reveal distinct transcriptional profiles of senescent versus non-senescent B cells. (E) Identification of p16/p21-positive B cells in DBiT-seq regions of interest (ROIs). Heatmaps show differentially expressed genes between p16/p21-positive and -negative pixels within follicular regions.

**Figure S10.**
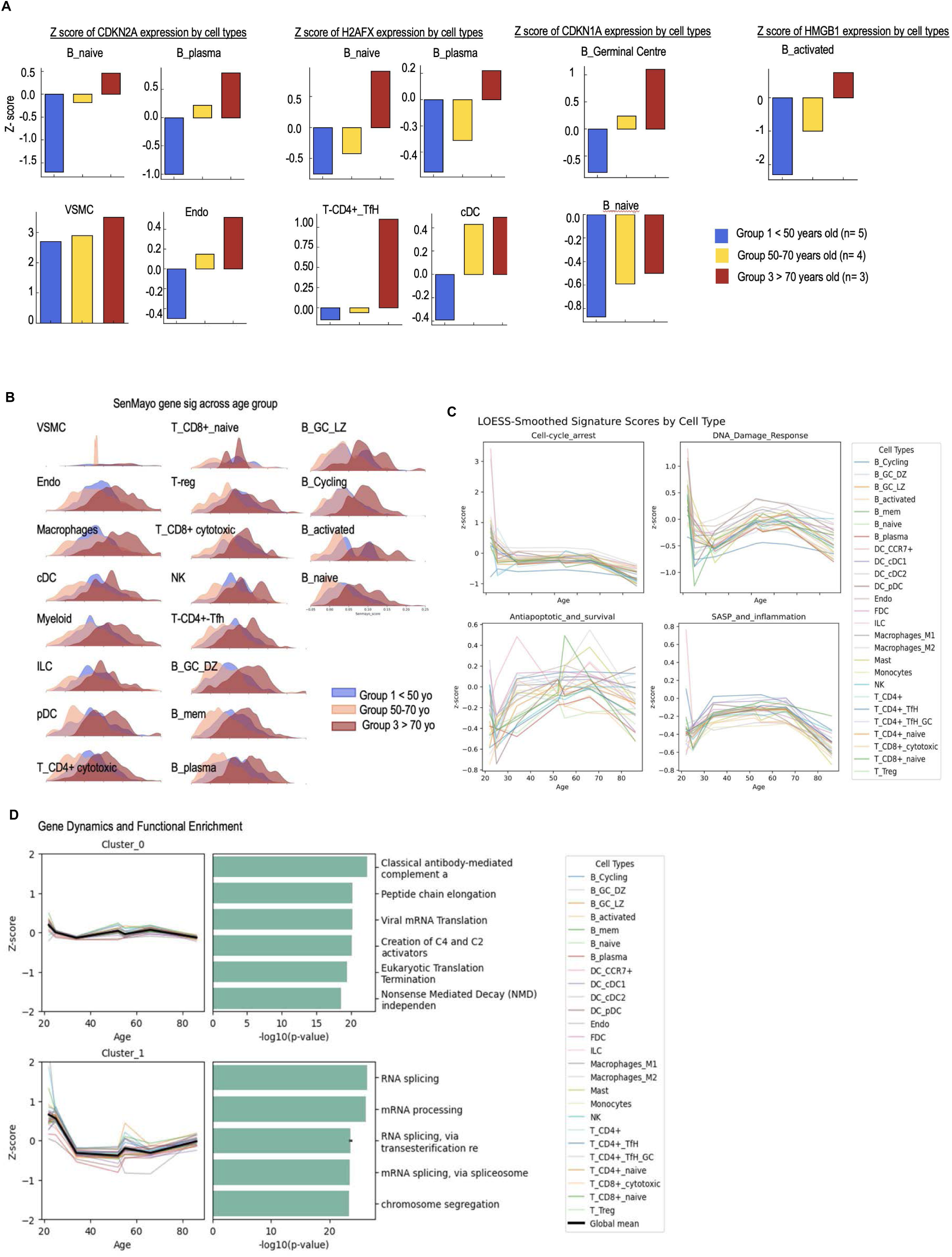
Age-associated transcriptional dynamics and pathway enrichment in lymph node cell types using integrated DBiT-seq and scRNA-seq analysis. (A) Bar plots showing Z-scores of CDKN2A, H2AFX, CDKN1A, and HMGB1 expression across selected immune and stromal cell types (e.g., B_naive, B_plasma, B_GerminalCentre, VSMC, Endothelial, T_CD4+_TH, cDC) stratified by age group (<50 , 50-70, and >70 years old). (B) Density plots of SenMayo gene set signature scores across cell types and age groups, highlighting increased senescence-associated transcriptional burden in specific B cell and other immune cell populations with age. (C) Line plots illustrating age-associated dynamics of DNA damage response, cell cycle arrest, anti-apoptotic/survival, and SASP/inflammation gene programs across cell types. (D) Gene dynamics and functional enrichment analyses for two identified gene clusters. Cluster_0 is enriched for antibody-mediated complement activation and viral mRNA translation, while Cluster_1 is enriched for RNA splicing, mRNA processing, and chromosome segregation pathways.

**Figure S11.**
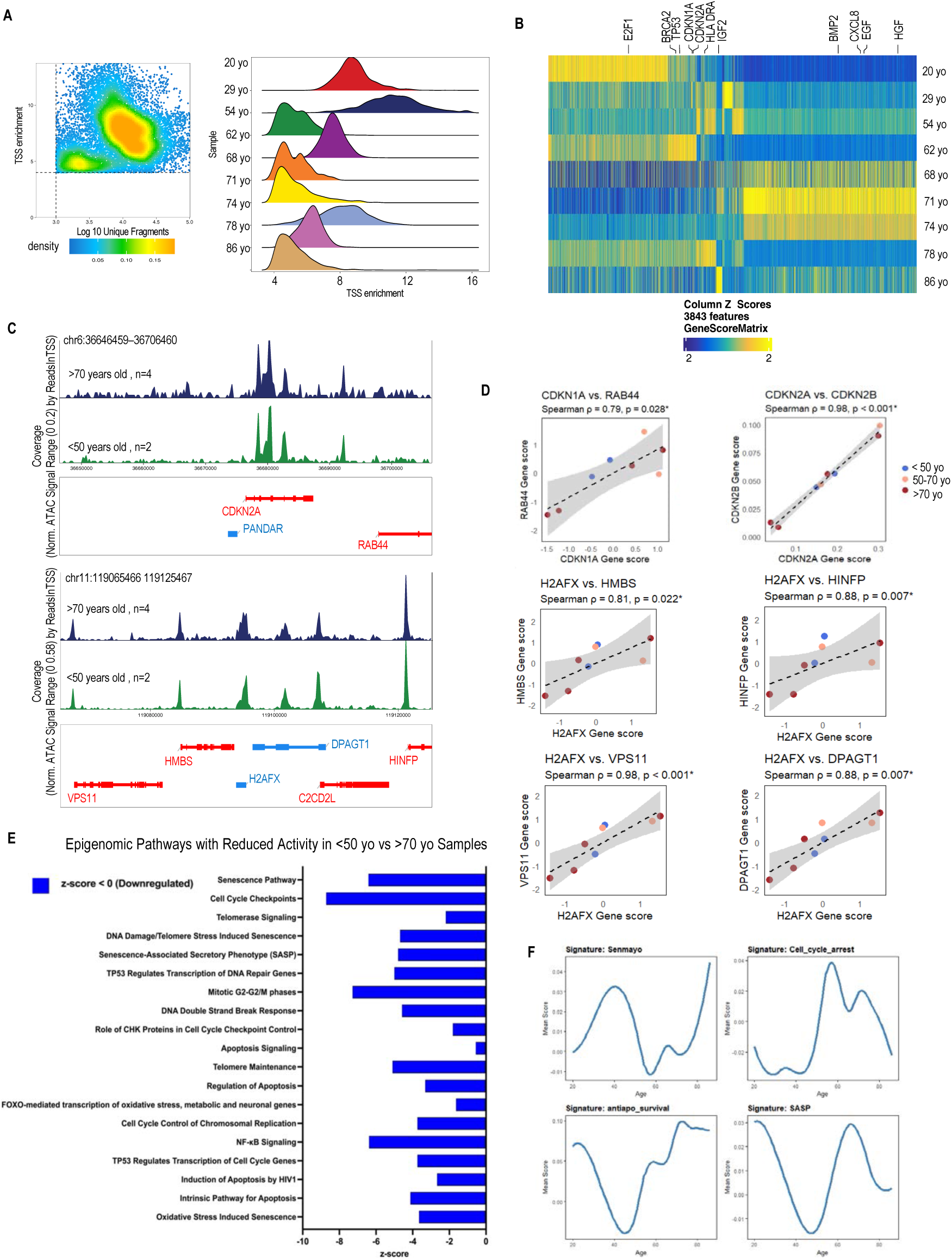
Spatial ATAC-seq analysis reveals age-associated chromatin accessibility changes in senescence-related genes. (A) Representative density and distribution plots of chromatin accessibility peaks across all cell types identified in spatial ATAC-seq data. (B) Heatmap of Z-scored chromatin accessibility for senescence-related genes (e.g., CDKN2A, CDKN1A, H2AFX, HMGB1) across donors of different ages. (C) Genome browser tracks depicting chromatin accessibility peaks at the CDKN2A and H2AFX loci in <50 and >70 years old samples. (D) Scatter plots showing the correlation between H2AFX or CDKN1A expression and that of neighboring genes (e.g., RAB44, HMBS, HINFP, DPAGT1, VPS11), with Spearman correlation coefficients and associated p-values indicated. Scatter plots showing correlations between H2AFX or CDKN1A gene scores and expression of and associated genes (e.g., HMBS, HINFP, DPAGT1, VPS11, PANDAR, RAB44) gene expression, with Spearman correlation coefficients and p-values indicated. (E) Bar plot showing pathways that are significantly downregulated in older compared to <50 years old samples based on chromatin accessibility analysis. The x-axis represents the –log10(p-value) of pathway enrichment. (F) line plots illustrate anti-apoptotic/survival, cell cycle arrest, SASP and SenMayo signature scores across different ages.

